# Pan-cancer Analysis Predicts Kindlin-associated Global Mechanochemical Perturbation

**DOI:** 10.1101/2022.10.31.514453

**Authors:** Debojyoti Chowdhury, Ayush Mistry, Riti Bhatia, Simran Wadan, Soham Chakraborty, Shubhasis Haldar

**Author notes:** Contributed equally to this work (Debojyoti Chowdhury, Ayush Mistry).

## Abstract

Kindlins are mechanosensitive adapter proteins that connect extracellular mechanical cues to intracellular chemical events. Any alterations in these proteins thus alter cellular signaling, which could result in cancer progression. However, their involvement in global mechanochemical signals remains elusive in cancers. Here we analyze pan-cancer samples to decipher how kindlin alterations aid cancer progression. We show that kindlin alterations, at both the genetic and mRNA level, dysregulates cellular behavior which significantly correlate with poor survival. We find that while these alterations are cancer-specific, they are prevalent in advanced tumor stages and metastatic onset. We observe that kindlins co-alter with a substantial fraction of human mechanochemical proteome in various tumors. Our analysis suggests how kindlin alterations aid tumor-promoting signals with a synergistic effect from alterations of cancer-hallmark genes. Notably, we demonstrate a consistent alteration of epithelial-mesenchymal-transition markers with kindlin activity. Overall, our study highlights how kindlin alterations could affect metabolism, genomic instability, and signal disruption via their interactome network, causing cancer and suggests targeting them as a therapeutic strategy.

## Introduction

Cancer is a multiplex physiological disorder projected to be nearly 1.5 times deadlier in the coming decades^1^. Severity of this disease originates from the rapid migration, protrusion, tissue invasiveness, metastatic capacity, and chemotherapy-resistant properties of malignant cells^2–4^. These processes are constrained by many internal and external cues, including mechanical force transmitted from extracellular matrix (ECM) proteins^5^. These Cell-ECM interactions are driven by a specialized transient structure called focal adhesion, which forms the mechanosensory hub by recruiting hundreds of force-sensing proteins to orchestrate intracellular rearrangement according to external mechanical cues^6^. Due to the presence of these proteins, the site-specific mechanical cues are translated into chemical signals within cells, further rewiring the genetic and epigenetic landscape, favoring cancer growth^7–9^. One such family of mechanosensing adapter proteins, the kindlins, convey the extracellular message of integrin outside-in signals via physical interactions to various structural proteins, receptors, and transcription factors, producing a plethora of chemical effects^10–13^. These proteins are explicitly related to almost every cancer-hallmark protein, showing their importance in cancer onset, progression and reccurance^14^. Kindlins are known to play important roles in signaling pathways corresponding to tumor-microenvironment manipulation^15^, cellular metabolism^16^, cell cycle progression^17^, transcriptional regulation^18^, and cancer stem cell regulation^19^. Any alteration or structural mutations in these proteins might affect their mechanochemical signaling in a global scale leading to altered homeostasis.

Mutations in the regulatory region of a gene either up or downregulate its expression, while coding mutations can alter the proteins’ stability, flexibility, and binding affinity for the respective ligands, thereby hindering signal transduction. However, the current data are insufficient to model the actual effect of mutations on the intrinsic properties, especially for the mechanosensitive proteins. The stochastic nature of tumor heterogeneity, due to the involvement of signal crosstalk and co-occurrent mutations in involved proteins, also makes the cancer systematics intricate and complicated to regard the effect of genomic and proteomic alterations^20,21^. Moreover, the current way of tracing a high-impact alteration solely based on a statistical analysis of clinical data belies the structural and chemical consequences^22^. Oversight of these aspects thus encumbers the coherent understanding of the intricate mechanisms of cancer progression.

In the present study, we have conducted a pan-cancer analysis of FERMT genes based on the clinical samples of International Cancer Genome Consortium (https://dcc.icgc.org), The Cancer Genome Atlas (https://www.cancer.gov/tcga), Catalogue of Somatic mutations in cancer (https://cancer.sanger.ac.uk/cosmic), and PCAWG^23^ study, utilizing genetic alterations, protein structural analysis tools, and gene ontology datasets to study the pan-cancer alteration effects of kindlin family proteins in mechanochemical signaling. While we show that kindlins are responsible for tumor progression, and the onset of metastasis, our analysis also reveals a prominent role of kindlins in epithelial-mesenchymal transition correlating to alterations in major mechanosensitive proteins. This study plausibly reveals an association of kindlin dysfunction with poor disease-free survival. Furthermore, using normal mode analysis (NMA), we have predicted the impact of cancer-specific mutations on the structural stability of kindlin proteins and how these mutations perturb their signal receiving and transducing ability, which in turn could regulate their downstream signal transduction pathways. Finally, this structural genomics approach successfully correlates the alteration effects to the clinical parameters substantiating the mechanochemical role of kindlins in different stages and subtypes of cancer.

## Results

### Kindlin alterations are found across multiple cancer types

Using our protocol (Fig. S1), three types of kindlin family genes were found altered in 45 different cancer types (sample size, n=2922), with FERMT1 (kindlin1 gene) being the major contributor (8%) followed by FERMT2 (kindlin2 gene, 6%) and FERMT3 (kindlin3 gene, 5%) (Fig. S2) above a z-score cut-off ±1.96 (p-value < 0.05). Majority of these alterations are either mRNA expression or gene amplification (Fig: 1a). Alteration frequencies are high in the case of non-solid (e.g., leukemia, lymphoma) or soft-tissue cancers (e.g., leiomyosarcoma) (Fig: 1a). FERMT1 alterations are found in 21 types of cancers whereas FERMT2 is altered in 15 types (Fig. S2a and b).

Kindlin expression level associates with stiffness-induced breast-cancer invasiveness and metastasis^24^. In our sample cohort, we found that both FERMT1 and FERMT3 mRNA are overexpressed in tumor samples. However, the increase in mRNA expression is not significant for FERMT3 when compared to FERMT1 (Fig. 1b). Interestingly, FERMT2 expression is significantly decreased in these tumors when compared to the normal samples (Fig. 1b, Fig. S3). Among blood cancers, expression of FERMT1 and FERMT2 is highest in T-cell lymphoma, unlike FERMT3 expression, which is lowest compared to other subtypes (Fig. S4-S6). Similarly, both FERMT1 and FERMT2 expression are higher in luminal breast cancer and MSS form of colon cancer (Fig. S4, and S5), while in gastric MSS, FERMT3 expression is higher (Fig. S6). FERMT1 expression increases in higher stages of cancer progression in BLCA, COAD, LUSC, and STAD (Fig. S7). Stage-specific decrease in FERMT2 mRNA expression is not pronounced except in renal cancer, LUAD, and BRCA (Fig. S7). The increase in FERMT3 mRNA expression shows significant stage-specific increase only in renal cancer and uveal melanoma (Fig. S7). These expression changes in kindlins are related to the overall mutational load in samples. With increasing mutations in cancer samples, FERMT1 and FERMT3 expression levels increase significantly, while FERMT2 shows an inverse trend (p^FERMT1^=4.184 x 10^-15^, p^FERMT2^=1.356 x 10^-14^, p^FERMT3^=6.687 x 10^-07^), albeit not so strong correlation (ρ_FERMT1_∼0.25, ρ_FERMT2_∼ -0.25, ρ_FERMT3_∼0.16) (Fig. 1d). To check whether the mRNA expression is consistent to the protein abundance, we analyzed the CPTAC pan-cancer proteome data from TCGA samples by comparing tumor and tumor-adjacent normal tissue. In contrast to the mRNA expression level, FERMT1 protein expression decreases in maximum cancer samples. However, FERMT3 protein expression shows a mixed nature of up-and down-regulation in a cancer-specific manner (Fig. 1c). To find the plausible cause, we examined the expression level of the kindlin-associated miRNAs from cancer samples (Table S1-S3). These results not only indicate miRNA-mediated kindlin expression in cancer-specific manner but also suggest a feedback-like looping between kindlin expression and miRNA expression. This connection of kindlin/miRNA axis to cancer progression and chemoresistance is consistent with experimental evidences^25,26^.

**Fig 1.**
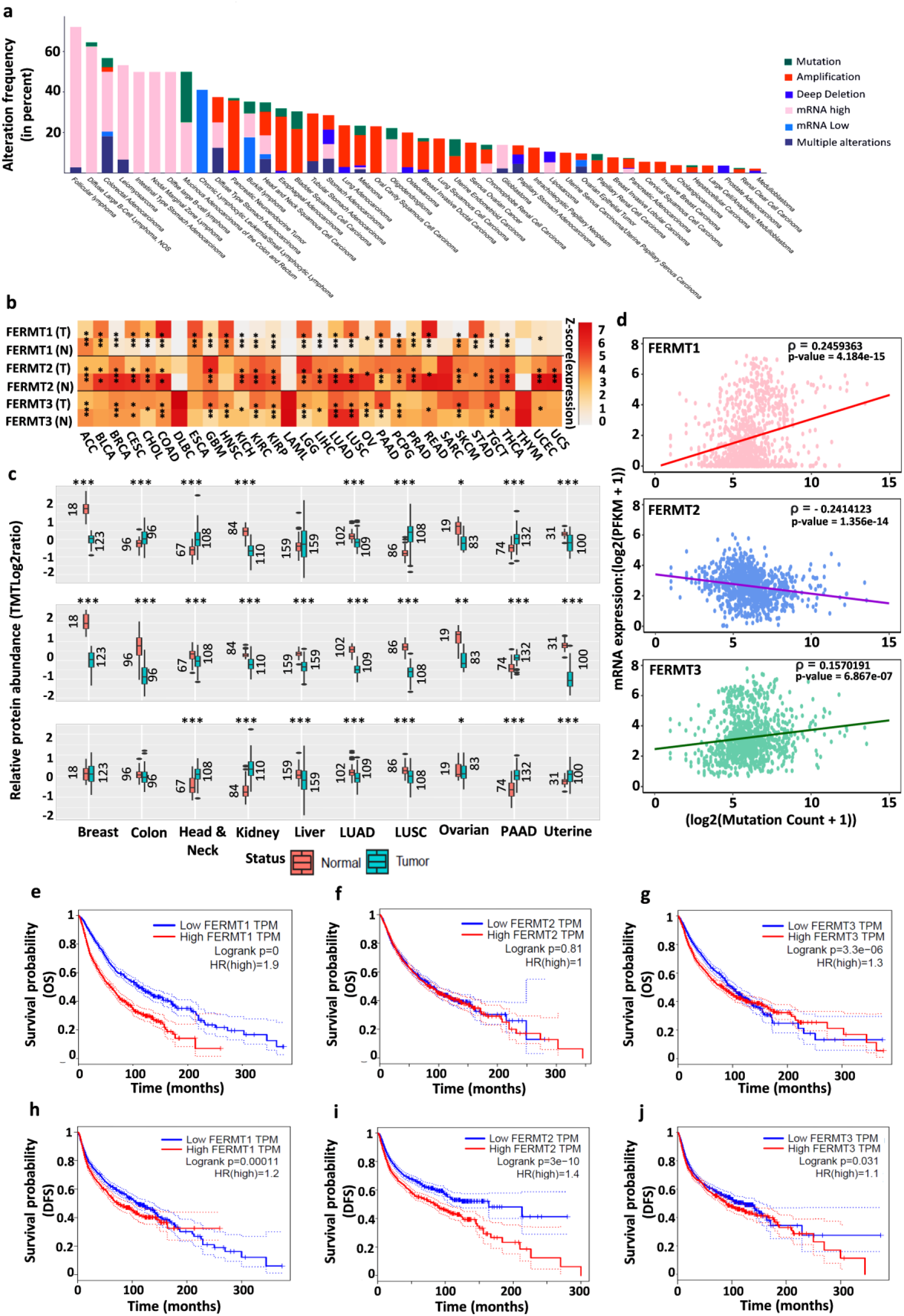
Pan-cancer framework of various alterations and expression change of Kindlin family proteins along with their clinical significance. a. Prevalence of kindlin proteins alterations in different types of cancers. The color codes indicate types of alterations as shown in the figure. b. Expression levels of kindlin mRNA in different types of cancer samples with respect to the corresponding normal tissues in terms of Z-score (±1.96 cut-off, p-value < 0.05). ***, p-value <0.0005; **, p-value <0.005; *, p-value <0.05; absence of * = no significance or absence of sample in a cohort, paired p-values were obtained by student t-test. c. Expression levels of kindlin proteins in different types of cancer samples with respect to the corresponding adjacent normal tissues in terms of Z-score (±1.96 cut-off, p-value < 0.05). ***, p-value <0.0005; **, p-value <0.005; *, p-value <0.05; absence of * = no significance or absence of sample in a cohort, paired p-values were obtained by student t-test. d. Changes in mRNA expression level with sample specific mutation count. The solid lines represent the regression line each case. Spearman’s correlation: FERMT1, ∼0.25 (P-value= 4.184 x 10^-15^); FERMT2, ∼ -0.24 (P-value= 1.356 x 10^-14^); FERMT3, ∼ 0.16 (P-value= 6.867 x 10^-7^). (e-g). Comparative quartile overall survival probability curve as Kaplan-Meier plot of high and low mRNA expression sample group in case of e. FERMT1 (Logrank P value, 0; Hazard ratio, 1.9); f. FERMT2 (Logrank P value, 0.81; Hazard ratio, 1); g. FERMT3 (Logrank P value, 3.336 x 10^-6^; Hazard ratio, 1.3). (h-j). Comparative quartile disease-free survival probability curve as Kaplan-Mayer plot of high and low mRNA expression sample group in case of h. FERMT1 (Logrank P value, 0.00011; Hazard ratio, 1.2); i. FERMT2 (Logrank P value, 3.0 x 10^-10^; Hazard ratio, 1.4); j. FERMT3 (Logrank P value, 0.031; Hazard ratio, 1.1). Quartile cut-off high: 75%, low: 25%; dotted lines corresponding to the survival probability lines represent 95% CI. 100-month cut-off survival time is designated as dotted line (parallel to y-axis) is shown in each image.

To investigate whether kindlin mRNA expression can correlate with cancer prognosis, we performed the expression-specific analysis of overall survival and disease-free survival (Fig. 1e-j). High expression of FERMT1 can be associated with poor overall survival (p < 0.001, hazard ratio (HR) = 1.9) (Fig. 1e), while overall survival appears to be independent of FERMT2 (Fig. 1f) and FERMT3 (Fig. 1g) expression. Interestingly, higher FERMT2 expression can be correlated with lower disease-free survival (Fig. 1i), unlike the other two FERMT genes (Fig 1h, 1j), suggesting its role in chemoresistance or cancer recurrence, as evidenced in a study by Ning et al. ^27^ Individual cancer analyses show FERMT1 overexpression as a prognostic marker in PAAD (p=0.03, HR=1.6) and SKCM (p<0.001, HR=1.7) (Fig. S8). Underexpression of FERMT2 may correlate with lower survival in BLCA (p=0.0036, HR=1.6) and STAD (p=0.034, HR=1.4) (Fig. S9). Interestingly, FERMT3 overexpression is prognostic for LAML (p=0.001, HR=2.5), while under-expression is prognostic for SKCM (p=0.0019, HR=1.52) (Fig. S10). FERMT1 overexpression corresponds to poor disease-free survival (DFS) in PAAD (p=0.0019, HR=2) (Fig. S11), while the same in the case of FERMT2 in ACC (Logrank p=0.034, HR=2.1) and COAD is noticeable (p=0.033, HR=1.7) (Fig. S12). FERMT3 underexpression in CHOL (p=0.0066, HR=4.0) and overexpression in UVM (p=0.00067, HR=6.7) are strongly associated with poor DFS (Fig. S13).

Apart from expression, we analyzed the copy number variation (CNV) of FERMT genes in 33 types of cancer (Fig. S14). Except for LAML, THCA, and PRAD, other cancer types show significant CNV of FERMT genes in the form of heterozygous CNV (Fig. S14a). FERMT1 and FERMT3 mostly show heterozygous amplification across cancer types as opposed to FERMT2, which shows mostly heterozygous deletion (Fig. S14b). DNA methylation, another key regulator of Kindlin gene expression, is also significantly altered in 14 cancer types (Fig. S15). FERMT2 hypermethylation is significant in tumors showing the most prominent effect in KIRP. Although FERMT1 is hypomethylated in most tumor types, it is significantly hypermethylated in BRCA. Independent of hyper- or hypo-methylation, changes in methylation amount in kindlin genes anti-correlates with gene expression suggesting a downregulation most saliently for FERMT3 (Fig. S15b). FERMT1 and FERMT2 hypermethylation are markers of survival risk in LGG (Logrank p < 10^-10^), ACC (Logrank p < 10^-5^) and KIRC (Logrank p < 0.05), SARC (Logrank p < 0.05) respectively.

### FERMT mutations are linked with tumor progression and metastasis

FERMT mutations are found in 31 cancer types with mutation frequencies of 15%, 14%, and 12% for FERMT1, FERMT3, and FERMT3, respectively (Fig 2a). Cancer-specific coding somatic mutations are mostly missense mutations, followed by silent and frameshift insertion or deletion (Indel) mutations (Fig. S16). These mutations are distributed throughout their sequences (Fig 2b). However, the mutation frequency within the FERM domain of FERMT3 is considerably high. In the case of FERMT2, we observe a mutational hotspot partially within the F1 domain. We have mapped the potential impact of mutations in the regulatory region on mRNA expression through the extent of loss of function, gain of function, or change of function from wild-type functionality (inferred from recurrence and multiplicity in tumor samples^28^ (Fig. 2c). Across FERMT1 and FERMT2, most of the high-impact mutations originate from the 5’UTR and upstream sequence. Because of these regulatory mutations, the expression of FERMT1 and FERMT2 has decreased consistently, whereas it increases for FERMT3. In addition, the impact of regulatory upstream or downstream mutations across FERMT3 has remained low contrasting to some start-loss, stop-loss, stop-gained, splice-region and intronic mutants which remain high-impact. The Kaplan-Meier curve shows a significant risk of survival conferred by FERMT mutations (p-value = 0.0003, HR = 1.932) (Fig. 2d). Upon further analysis, we found that FERMT1 and FERMT3 mutations have almost the same survival risk, and are higher compared to the FERMT2 mutation-associated risk (Kruskal-Wallis rank sum p-value: FERMT1-FERMT2, 0.0058; FERMT2-FERMT3, 0.0002; FERMT1-FERMT3, 0.2945) (Fig. 2e). Tumor stage-specific mutation analysis shows an almost similar trend for all kindlins, as most of these mutations are found at tumor stages T2 and T3, indicating their effect on tumor progression rather than onset (Fig. 2f). FERMT mutations are also significant in the metastatic M0 stage compared to later stages, suggesting their role before metastatic onset (Fig. 2g).

**Fig 2.**
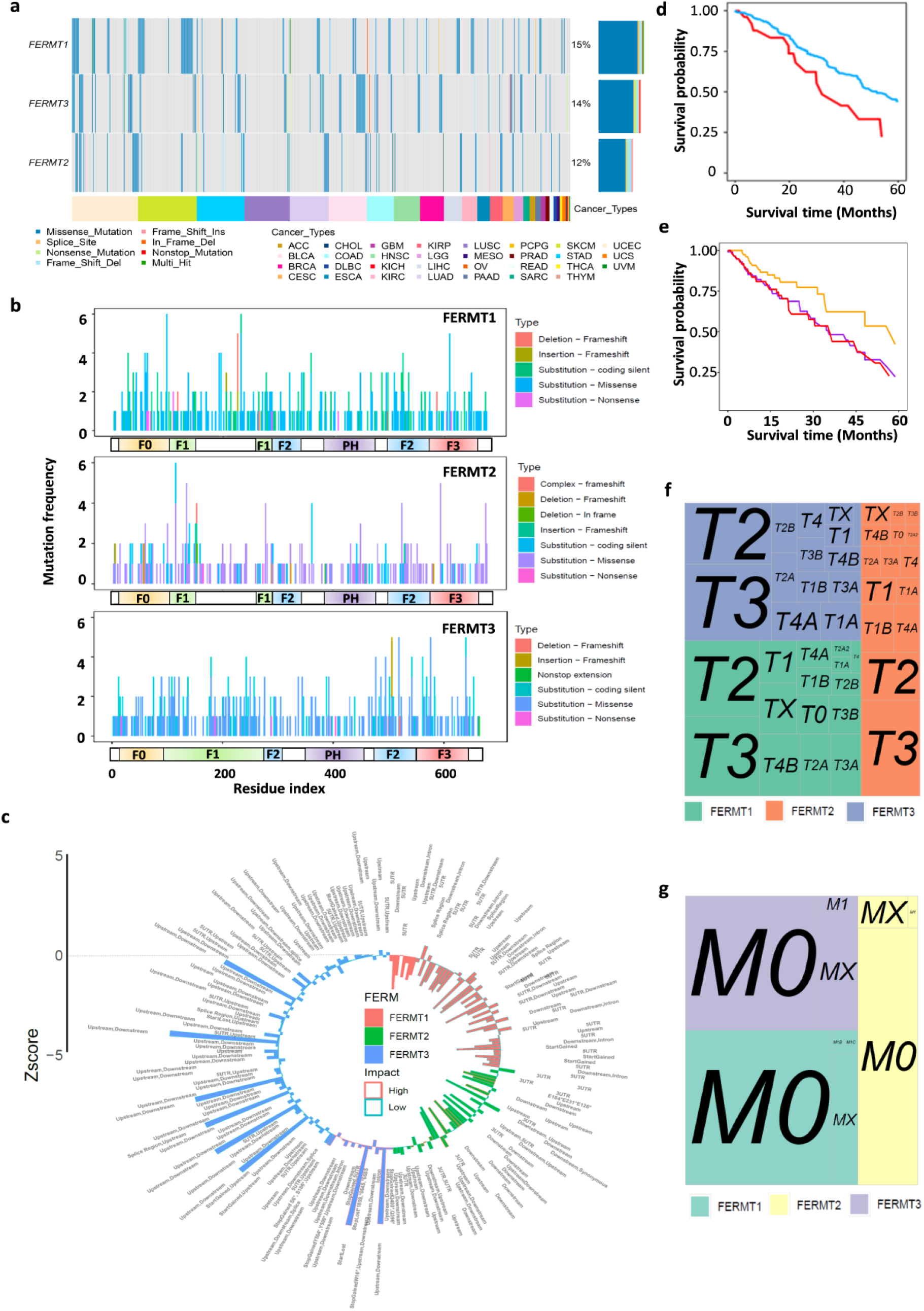
Characterization of Kindlin mutations in patient cohort. a. Mutation frequency (in % samples) of kindlin coding mutations across various types of tumor samples (Colors are representative of types of cancer in lower horizontal bar). Different mutation types are also represented with different colors in respective patient samples. b. Residue specific distribution of somatic mutations across all three kindlins. Types of mutations and their respective color coding are presented with respective images. Below the corresponding sequence-specific mutation frequency graphs, kindlin domain organization is given. Each color in these represents specific domains. c. Comparative plot of Z-score values for differentially expressed mutated (non-coding) kindlin transcripts (significance cut-off z-score ±1.96, p-value < 0.05). Different color-coded bars indicate different kindlins. Cancer-specific impact of the mutations are denoted by contrasting border colors of the bars. Positions of the non-coding mutations are indicated in text corresponding to each bar. d. Comparative Survival time versus survival probability curve for kindlin mutated (red) and non-mutated (blue) sample cohort. (p value = 0.0003, Hazard ratio, 1.932). e. Comparative Survival time versus survival probability curve for different kindlin mutated sample cohort. FERMT1, pink; FERMT2, yellow; FERMT3, purple; (Kruskal-Wallis rank sum p value: FERMT1-FERMT2, 0.0058; FERMT2-FERMT3, 0.0002; FERMT1-FERMT3, 0.2945) f. Tumor stage specific mutations in kindlins. The area of each quadrilateral and corresponding size of tumor stage name text indicates the amount of alteration. g. Metastasis stage specific mutations in kindlins. The area of each quadrilateral and corresponding size of metastatic stage name text indicates the amount of kindlin alterations involved.

### Mutations affects structure-function dynamics of kindlins

Kindlin family proteins show ∼49-58% sequence identity and ∼67-73% sequence similarity, but they are structurally similar (Fig. S17). To study the effects of mutations on structural stability of kindlins, we calculated the ΔΔG values of all the cancer-specific mutated conformations of all kindlin types. Since the mechanochemical activity of kindlins comes from their domain-specific flexibility, we calculated the vibrational entropy change (ΔΔS) of mutants concerning the wild type versions. Our analysis reveals four different populations of these mutants: both high flexibility and stability (Q1), low flexibility and high stability (Q2), both low flexibility and stability (Q3), and high flexibility and low stability (Q4) (Fig. 3a-c). We also get a trend towards decreasing stability with increasing flexibility (p<2.2e-16; σ_FERMT1_= -0.7470601; σ_FERMT2_=-0.8190608; σ_FERMT3_=-0.7077385). Furthermore, we classified the mutants into five categories: very high, high, moderate, low, and slight for each stabilizing and destabilizing cohort (Table S4-S6). Almost 50% cancer-specific mutants of all kindlins are loss-of-function and highly disease causing, while the rest ∼50% are tolerable in cells (Fig. 3d). Most of the loss of function mutants are from both very high stability/low flexibility and very high flexibility/very low stability region i.e., Q2 and Q4, respectively. ΔΔG analysis of respective dimers also shows a common destabilizing effect for all the mutant dimers (Fig. 3e-3g). Mutation-induced protein structural alterations can lead to changes in chemical properties due to altered intramolecular interactions (Fig. S18-S23). Our analysis predicts pronounced changes in hydrophobicity in mutant monomers and dimers of all kindlins (Fig. S24, S25). However, no significant changes in intramolecular salt bridge interactions are seen considering minimally altered κ-values, a parameter indicating extent of intramolecular salt-bridge formation (Fig. S26, S27).

**Fig 3.**
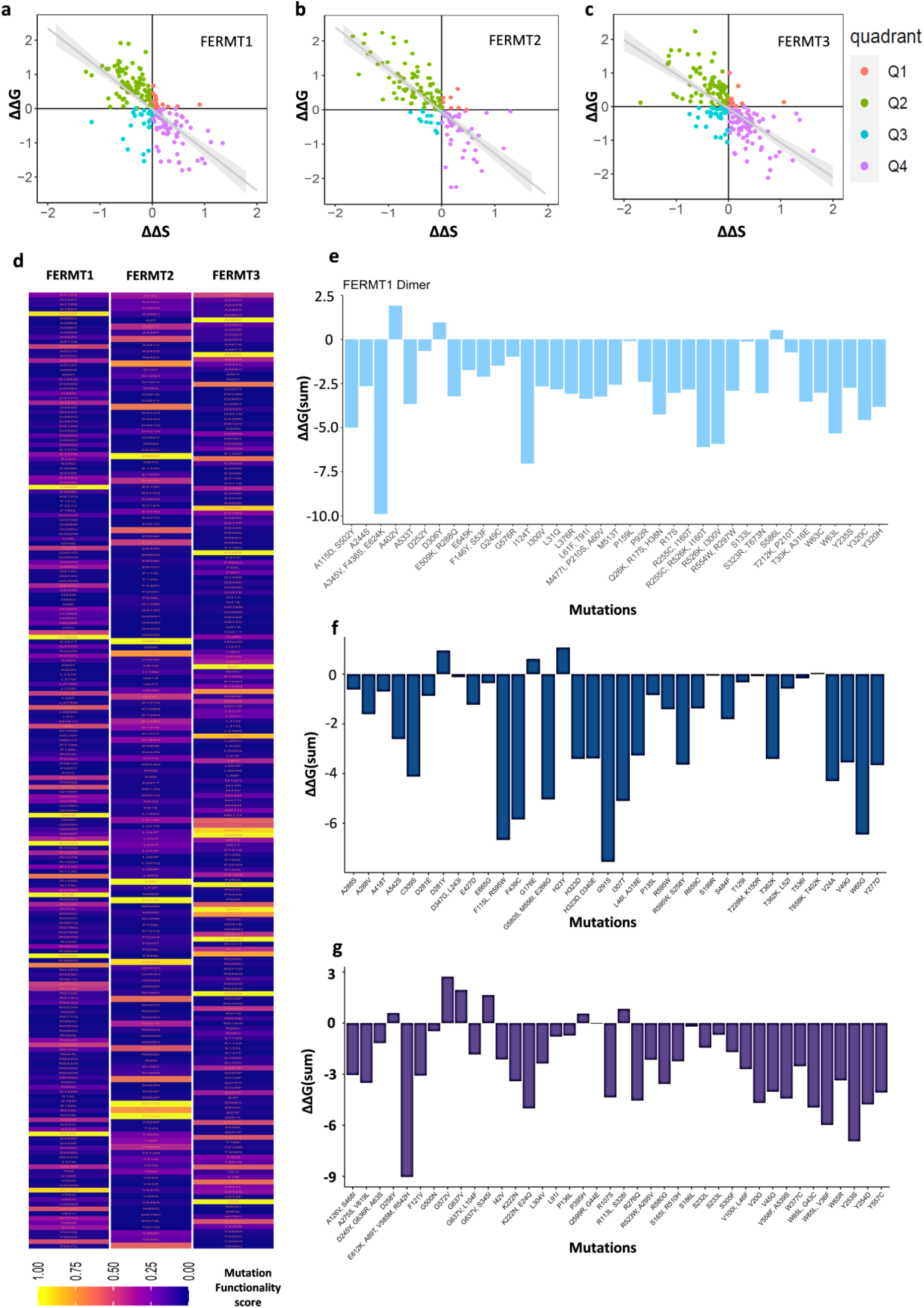
Stability analysis of mutated Kindlins. Only missense mutations (either single or multiple mutation variants) are considered here. (a-c). ΔΔG vs ΔΔS plots to determine stability of the mutants against their flexibility for a. kindlin1 (FERMT1) b. kindlin2 (FERMT2) and c. kindlin3 (FERMT3). ΔΔG values and ΔΔS values are presented in KJ/mol and KJ/(mol.K) unit respectively. The plots are divided into four quadrants (Q1-Q4) according to the nature of four different mutant population. Q1, increased stability and flexibility (red); Q2, increased stability and decreased flexibility (green); Q3, decreased stability and flexibility (blue); Q4, decreased stability and increased flexibility (purple). Regression lines are shown in gray. d. Mapping kindlin mutations according to their plausible impact on functionalities. Values <0.05 indicates loss-of-function mutants, rest are neutral mutants. (e-f) Stability changes in mutated protein dimer structures with respect to wildtype in Sum ΔΔG values (KJ/mol). d. kindlin1 (FERMT1) e. kindlin2 (FERMT2) and f. kindlin3 (FERMT3).

Phosphorylation is another important aspect of kindlin functionality, which has been validated experimentally at T8 and T30 position for kindlin1; Y193, S159, S181, and S666 for kindlin2; T482 and S484 for kindlin3^29^. Computational predictions indicate a complete loss of T8 and S484 mutation sites in kindlin1 and kindlin3, respectively. For FERMT2, all frameshift mutants show a complete loss of Y193 and S666 phosphorylation sites (Table S7-S9). These structural effects on phosphorylation correlate with patient-specific phosphorylation level. From phosphoproteomic tandem mass tag (TMT) data, an overall decrease in phosphorylation is observed for all three kindlins unlike FERMT2, which shows that elevated phosphorylation levels (Fig. S28) might be due to altered phosphorylation sites. Overall, this affected phosphorylation might be attributed to the altered kinase activity of ser-thr/ tyr kinases, a signature of tumor cells^30^.

With all kindlins, insertion-deletion (in-del) frameshift mutations lead to the loss of one or more biologically active domains in the protein (Fig. 4). Most of the FERMT3 in-del mutants except D231del, retain all domains with either altered sequence or altered three-dimensional domain structure. Similarly, K154del kindlin2 mutant retains all domains as wild type but with distorted dimerization domain, FERM and PH domain, indicating a loss of important functionalities (Fig. 4b and d inset).

**Fig 4.**
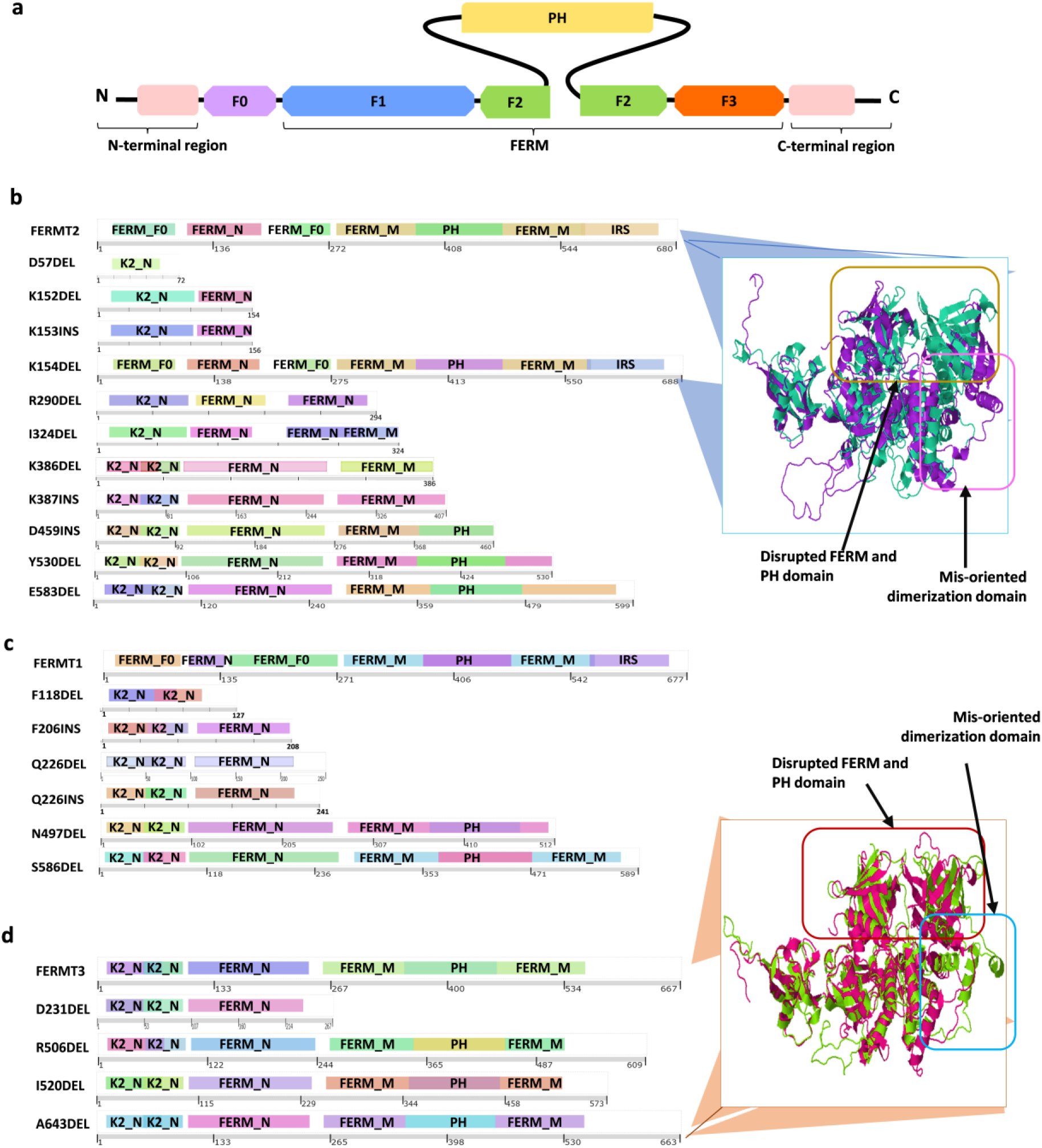
Functional impact of frameshift mutations of Kindlins. a. Common domain organization of kindlins as a schematic diagram. b. Comparison of domain organization of wildtype kindlin2 (FERMT2) with its cancer-specific indel variants (Inset: Superimposed structure of WT and K154del mutant containing all the domains show an altered orientation of various domains). c. Comparison of domain organization of wildtype kindlin1 (FERMT1) with its cancer-specific indel variants. d. Comparison of domain organization of wildtype kindlin3 (FERMT3) with its cancer-specific indel variants (Inset: Superimposed structure of WT and A643del mutant containing all the domains show an altered orientation of various domains). Differently colored domains indicate their similarity with pfam annotated domains from different organisms.

### Kindlin mutations cause alterations in mechanochemical signal transmission

Kindlins are known to carry out their mechanochemical signal transduction by their domain-specific movement (Fig. 5a). This mechanotransduction and its perturbations can be measured by normal mode analysis (NMA)^31^. We have performed NMA of the mutant and the wild-type kindlins in the presence of a computationally predicted force range of 87 pN^31^. Previously, it was shown that force transmitted to kindlins through integrin outside-in signaling strengthens the dimerization of kindlins reinforcing integrin condensation at focal adhesion sites^31^. Our analysis reveals force-induced movements of three specific regions- the F0 domain that interacts with the F-actins and paxillin^32,33^; the PH domain that connects plasma membrane via PIP2 and might respond to membrane-mediated signaling events^34,35^; and F3 domain that directly binds to integrin, TGFβRI, and RTKs through F3 domain^36,37^ (Fig 5b). B-factor analysis of force-induced root mean squared fluctuations (RMSF) predicts any changes in atomic fluctuation of cancer-specific mutants in all three kindlins (Fig. S29), with significant changes in the very highly stabilizing/destabilizing and multiple mutants, as might be expected. This change is evident for kindlin1 and kindlin2 in their dimerization domain, PH, and F3 domain (Fig. 5b). In the W65G mutant, the kindlin2 dimerization region shows a high RMSF, plausibly destabilizing dimerization, while C309S loses its flexibility in the F-actin/paxillin binding region (Fig. 5b). The multiple mutant R255C-R526K-I160T variant and the R526K-I300V variant of kindlin1 appear to lose the flexibility of the F3 domain, indicating a loss of the spring-like force-transmitting property (Fig. 5b). F3 mutants show minimal changes in domain-specific RMSF (Fig. 5b). The changes in the dimerization properties of the mutants are also evident from the prediction of the dimerization affinity (Fig. 6a-6c). Kindlin2 completely loses its dimerization affinity in all mutants (Fig. 6b). In contrast, kindlin3 mutants appear more stably dimerized (Fig. 6c), while kindlin1 mutants show a mixed effect on the dimerization trait (Fig. 6a) suggesting altered kindlin-dimer functionalities.

**Fig 5.**
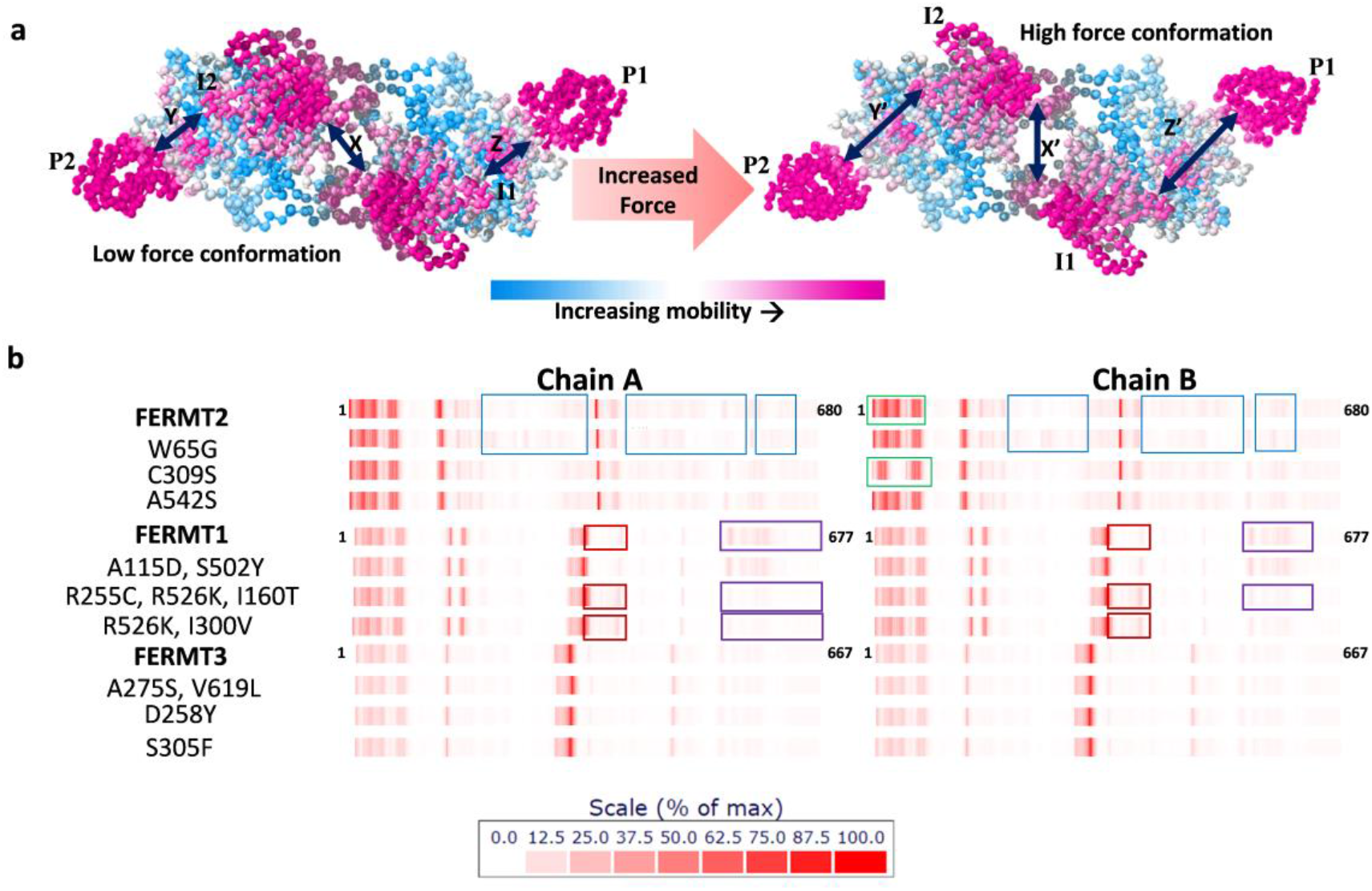
Mutation induced root mean squared fluctuation perturbations in Kindlins. a. Force induced normal mode trajectory of the wildtype kindlin dimers (superposed). Mobility of atoms are defined by a given scale with contrasting colors. I1, integrin binding domain 1; I2, integrin binding domain 2; P1, F0 domain of Chain A; P2, F0 domain of Chain B; The distance between I1 and I2 are given by X (Low force conformation) and X’ (High force conformation). Similarly, distance between I1(Low force)-P1 (Low force), Z; I2(Low force)-P2(Low force), Y; I1(High force)-P1 (High force), Z’; I2(High force)-P2(High force), Y’. b. Residue-wise B-factor analysis of kindlins and their three most-perturbed mutants each. Extent of redness indicates increase in B-factor. Same colored boxes are used to show similar position with altered B-factor with respect to the wildtype. Different colored boxes are used if alteration is different in some mutant(s) or in different positions. Numbers represent the start and end amino acid residue index in each chain.

**Fig 6.**
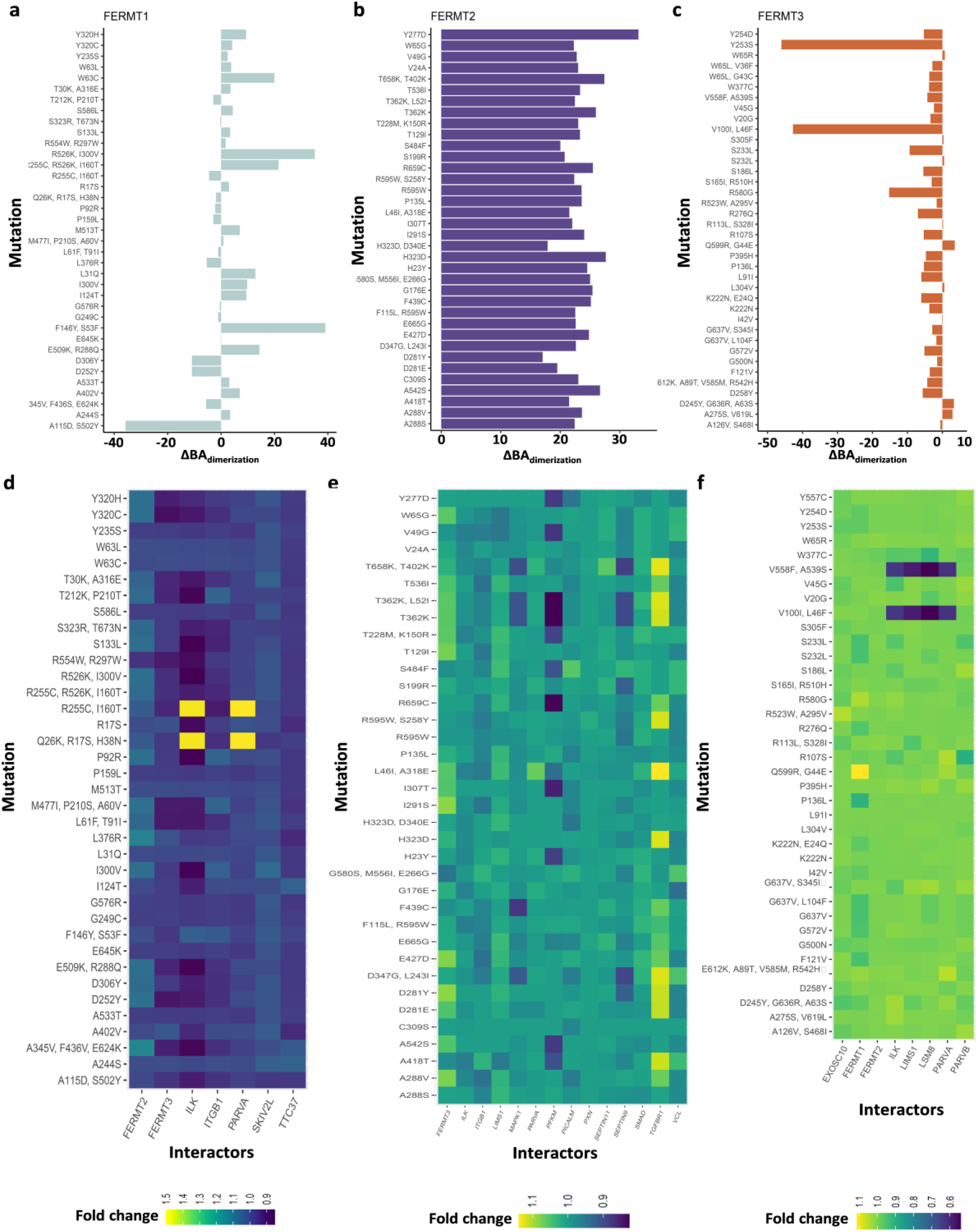
Mutation induced perturbation in binding affinity of Kindlins. Change in dimerization affinity with respect to the wildtype is calculated as ΔBA_dimerization_ (KJ/mol) and plotted for all the significantly perturbing mutants of FERMT1 (a), FERMT2 (b) and, FERMT3 (c). (d-f) represents mutation induced fold change (mutant/WT) in binding affinity of the mutant-direct binder interactions of FERMT1 (d), FERMT2 (e), and FERMT3 (f). Fold change of 0.1 = 24 KJ/mol for FERMT1, 22.7 KJ/mol for FERMT2, 23.1 KJ/mol for FERMT3. Interaction fold change>1, increased interaction; Interaction fold change<1, decreased interaction; Interaction fold change=1, affinity same as the wild-type. (Reference for cellular significance: energy stored within one high-energy molecule (ATP)∼30 KJ/mol.)

We further analyzed the signaling properties of kindlins and their changes in kindlin mutants using the Markov chain model^38^ (Fig. S30-S41). In the case of kindlin1, the receiving signal rate was sporadically perturbed across the sequence for multiple mutations, whereas transmitter signals were significantly unaffected. For single missense mutations, the recipient signaling rate is slightly perturbed within the 200-500 residue, along with trends in the 1-150 range for a tryptophan-removing mutation. However, the broadcasting signal rate is largely undisturbed in kindlin1 mutants. For Kindlin2 containing multiple mutations, there is some degree of interference across all FERM domains with receiver signaling rates. The broadcasting signal rate is not disturbed as much. Both receiving and transmitting signal rates are sporadically and significantly disrupted across all FERM domains for multiple mutations of Kindlin3 (Fig. S38-S41). As for Kindlin3 single mutants, the broadcaster rate appears to be altered in the F0, F1 and parts of the F2 domains, whereas the receiver signal affects the 200-450 amino acid index more significantly.

### Kindlin alterations are associated with global mechanochemical signal perturbations favoring cancer progression

The change in signal-receiving or signal-broadcasting properties of the mutant proteins might affect their ability to convert mechanical signals into biochemical cues, causing altered physical interactions. Our predictions of binding affinity change indicate altered kindlin-partner interactions in cancer-specific mutations. While kindlin1 mutants appear responsible for minor changes in homeostatic interactions, kindlin2 and kindlin3 cause a considerable decrease (Fig. S42). Most of these changes are either mutation-specific or interactor specific, or both. For kindlin1, its interaction with ILK is most affected among the mutants, except for two multiple mutants R255C-I160T and Q26K-R17S-H38N, where the interactions increase (Fig. 6d). Depending on highly-destabilizing kindlin2 mutations, we found a 0.2-fold decrease in affinity for PFKM and an increased affinity of highly stabilizing and destabilizing mutants to TGFBR1 and FERMT3 (0.2-fold 0.1-fold respectively) (Fig. 6e). Kindlin3 interactions are slightly disturbed and show a trend toward decreasing affinity. However, in two Kindlin3 multiple mutants, V558F-A539S and V100I-L46F, their interactions with ILK, LIMS1, LSM8, and PARVA decrease sharply (Fig. 6f).

Kindlins subtypes form huge interacting networks due to their function as adapters (Fig. S43-S45) linking many major biological processes (Fig. S46). Mutations can alter their homeostatic interactions with wild-type interactors, while the interaction partners might also be altered in cancer samples triggering a synergistic effect. Therefore, we checked the co-alterations of the corresponding physical interactors in cancer. In FERMT1 altered samples, TTC37, SKIV2L, and FERMT2 show maximum alterations compared to unaltered samples (Fig. 7a). Similarly, in FERMT2 altered cohort, its partners TGFBR1, SEPTIN9, SEPTIN11, PXN, and PFKM are mostly co-altered (Fig. 7b). FERMT3 co-alters to almost the same extent as all its interactors (Fig. 7c). Notably, this also demonstrates a strong mutual exclusiveness (tendency towards co-occurrence) in alterations of all the kindlins in cancer (p-value <0.001, q-value <0.001, log2 odds ratio: FERMT1-FERMT2 >3, FERMT2-FERMT3 =2.77, FERMT1-FERMT3 =1.93).

**Fig 7.**
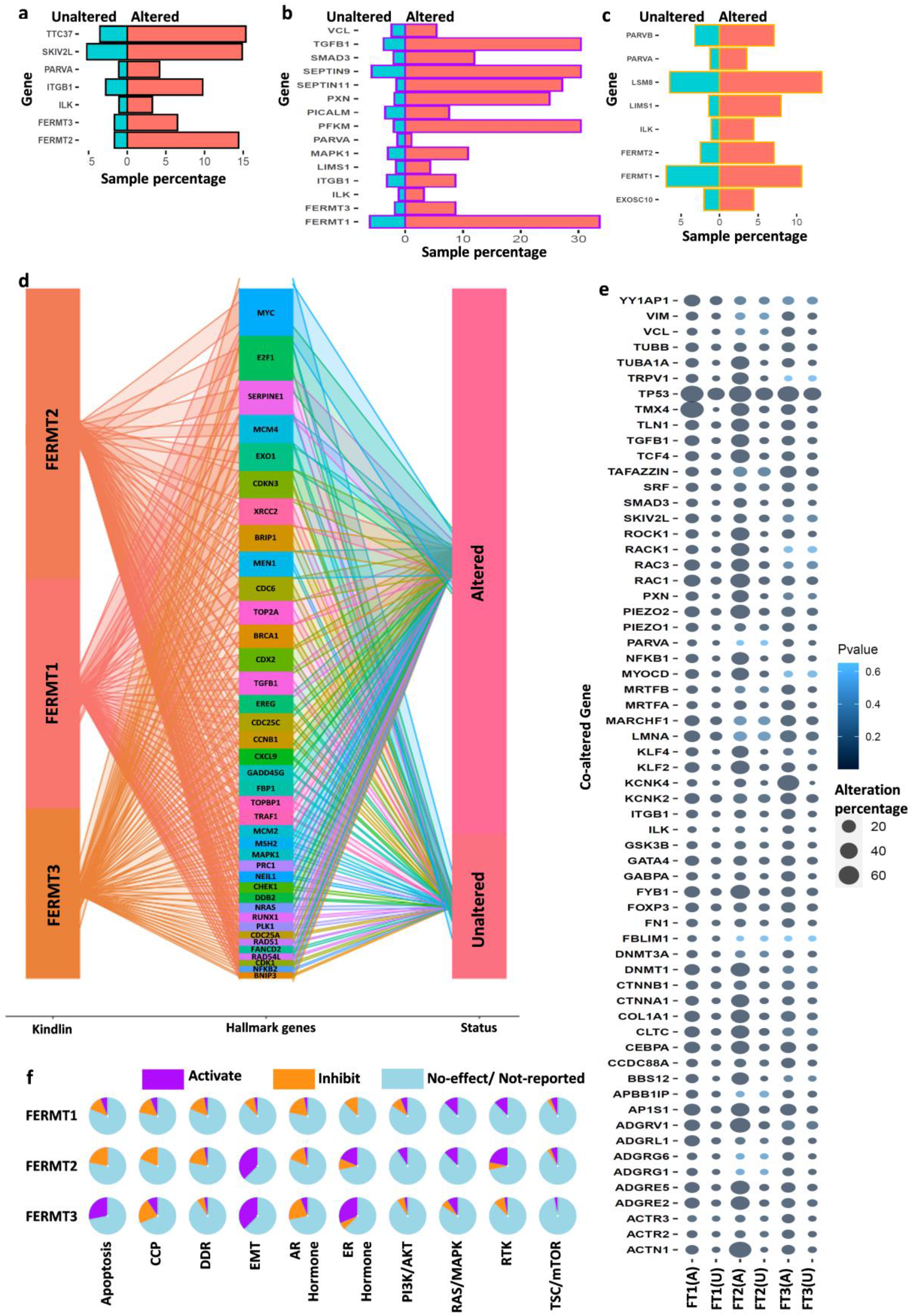
Co-alteration analysis of direct interactors of Kindlins, cancer hallmark genes and Kindlins associated mechanochemical signal proteins and their effect on major cancer-associated pathways. (a-c). The alteration frequencies (in percentage) of direct interactors of corresponding kindlins in respective kindlin-altered and non-altered cancer samples (PCAWG/ICGC/TCGA cohort): FERMT1 (a), FERMT2 (b), and FERMT3 (c). d. The alteration frequencies (in percent) of major cancer hallmark genes with respective kindlin altered and unaltered cancer samples (PCAWG/ICGC/TCGA cohort). Different colored chords connecting hallmark genes and status columns indicate respective hallmark genes. The breadth of the chords is directly proportional to the number of altered or unaltered samples. e. The alteration frequencies (in-percent) of major kindlin associated proteins involved in mechanochemical signalings with respective kindlin altered and unaltered cancer samples (PCAWG/ICGC/TCGA cohort). The size of the circles is indicative of percentage altered/unaltered samples. The colors represent p-values as given aside the image. Significance cut-off p-value<0.05. f. Influence of kindlin (FERMT1, 2, 3) alterations in signature cancer-related pathways in 32 different cancer types. CCP, cell cycle progression; DDR, DNA damage response; EMT, Epithelial-mesenchymal-transition; AR, Androgen receptor; ER, Estrogen receptor; RTK, Receptor tyrosine kinase pathway. A pie represents 100%.

Kindlins alterations accompany significant alterations in major cancer hallmark genes (Fig. 7d). For a better understanding of the synergistic effects via co-alteration analysis, we introduce a term ‘co-alteration dynamics’ of a biological process which denotes the average effect of co-alterations of the components of that particular biological process with respect to one or more components. For example, we calculated co-alterations of cancer-hallmark gene set with respect to kindlins, signifying co-alteration dynamics of cancer-hallmarks with respect to kindlins. Our co-alteration analysis containing 39 hallmark genes shows the most prominent co-alterations with FERMT2 (mean co-alteration dynamics=14.98) followed by FERMT1 (mean co-alteration dynamics=11.08), FERMT3 being the lowest (mean co-alteration dynamics=5.35). Kindlins act as interlinks of major cellular pathways, including other mechanosensitive or mechanochemical proteins directly linked either to kindlins or through kindlin interactors. Our meta-analysis revealed 62 proteins encompassing mechanochemical transcription factors, receptors, ion channels, cytoskeletal proteins, and other various types (Table S10). Almost all of these mechanochemical proteins are co-altered with all the Kindlins (Fig. 7e). Kindlin alterations mostly co-occur with ACTN1, ADGRs, DNMT1, RAC1, TMX4, and TP53. Here also, mechanochemical protein forming genes show the most prominent co-alterations with FERMT2 (mean co-alteration dynamics=19.3) followed by FERMT1 (mean co-alteration dynamics=12.34) and least co-alteration with FERMT3 (mean co-alteration dynamics=9.92). Interestingly, most of these co-altered mechanochemical proteins are either transcription factors or cytoskeletal proteins. To link these kindlin-associated co-alterations to cancer progression, we analyzed the contribution of kindlins in ten major cancer-associated pathways. FERMT1 alterations correlate with inhibitory effects on these pathways, whereas FERMT2 and FERMT3 alterations activate cancer-signature pathways of concern more frequently (Fig. 7f). However, FERMT1 alterations correlate with inhibition of apoptosis, cell cycle progression, DNA damage response, and androgen-receptor pathway like FERMT2 but to a lower extent (Fig. 7f). Interestingly, FERMT3 and FERMT2 alterations are significant in epithelial-mesenchymal transition (EMT). EMT-specific pathway enrichment analysis reveals a strong correlation between EMT-promoting processes like UV response downregulation, TGFβ signaling, angiogenesis, and hedgehog signaling to their alterations in 33 cancer types (Fig. S47). In parallel, EMT-inhibiting pathways like DNA repair, oxidative phosphorylation, and P53 tumor suppression are strongly anti-correlated with kindlin alterations (Fig. S47). FERMT2 alterations are also consistent with EMT-related immune filtrations supporting their significance in EMT (Fig. S48). While, FERMT2 correlates with the inhibition of apoptosis, we find consistent FERMT3 alterations with the activation of apoptotic pathways in cancer samples (Fig. 7f). Kindlins also correlate with other hormonal pathways; PI3K/AKT, mTOR, RTK, and MAPK pathways which are significant for any type of cancer progression.

## Discussion

Clinical advancements in precision oncology currently face two major problems: understanding the tumor heterogeneity and predicting the trend in the intracellular complexity arising from modulated microenvironment. Mechanosensitive adapter proteins play an indispensable role in these events linking extracellular mechano-environment with intracellular events through molecular clutch dynamics^39^. Thus, alterations of these proteins could become the Achilles’ heel for cellular homeostasis, favoring malignancy. In fact, intra- and extracellular mechanical events are found to guide cancer progression, metastasis, and recurrence^40–42^. Therefore, analysing the role of kindlins as one of the mechanosensitive adapters involved in diverse biochemical signaling and other regulatory functions might answer the mechanisms of cancer complexity and heterogeneity.

Previously, in the case of PAAD, it was found experimentally that kindlin downregulation contributes to intra-tumoral heterogeneity^43^. Based on the nature of the kindlin distribution in normal tissues, we observe genomic alterations in cancers originating from different tissues. This leads us to propose a plausible role for changes in the kindlin family genes in contributing to regulating tumor heterogeneity. This heterogeneity corresponds to activation of different cellular properties within tumor cells. We have shown that kindlin-mediated biochemical alterations arise due to combined alterations of kindlins and their networks. Kindlin-mediated cancer-specific upregulation or downregulation of miRNAs can also be important for inducing malignancy and metastasis. Our analysis suggests an interesting feedback-loop mechanism of kindlin and miRNA expression, which has also been shown in breast cancer malignancies^44^. We observed that miRNAs, regulated by kindlin2, target FERMT1 or FERMT3 mRNA. Another interesting observation is the correlation between total mutation and kindlin1, kindlin3 expression levels along with an anti-correlation between increase in genomic mutation and kindlin-2 expression. This advocates in favour of kindlin-mediated regulation of genomic instability, as was found experimentally by Zhao et al. for breast cancer^45^. This suggests that kindlins activities are dependent on a balance of stability and flexibility. When this balance is perturbed, it will lead to disconcerted mechano-chemical signal transmission.

Kindlin dimerization is a key factor for integrin activation and mechanosensing. In the presence of ITG1BP1, kindlin-integrin interaction is competitively hindered and hence, the focal adhesion loses its stability. We found a trend of kindlin-dimerization disruption for mutated kindlin2. Moreover, ITG1BP1-Kindlin2 co-alteration is also anti-correlated (p-value <0.05, q-value <0.05, percent-altered in Kindlin2 altered cohort (n=195) ∼1%, percent-unaltered in Kindlin2 altered cohort (n=2588) (∼8%) which might ease focal adhesion disruption and hence aid cancer cell migration. All the other important focal adhesion proteins like VCL, PXN, ITGB1, and LSM8 are also significantly altered (Fig. 7a-7c) showing a prominent cancer-specific disturbance in the cell’s kindlin-associated mechanochemical hub^46^.

Kindlin interactions are regulated in cancer either by expression or by kindlin mutations. Both mutation specific binding affinity and co-alteration of these direct interactors point to a global perturbation in signaling modules and their specific effects relating to wrong input of mechanosensing via kindlins. We analysed the functional involvements of kindlin-interactors from data available in Genecards (www.genecards.org) to get plausible altered roles of kindlins in cancer. Kindlin-1 direct-interactor TTC37 and SKIV2L are involved in protein folding due to their chaperone activity along with their role in super-killer complex. Both of them are significantly altered in cancer, suggesting a plausible path of kindlin-mediated protein folding and mRNA regulation alteration. PARVA, an important protein in sarcomere organization is co-altered significantly with all the three kindlins. Moreover, mutation induced altered interaction also suggests perturbed kindlin-mediated PARVA action in cancer. ILK and PARVB are known to form a complex to regulate cell motility, epithelial polarity, and the TNF pathway^47^. Kindlins plausibly interacts with either ILK (Kindlin1/2/3) or PARVB (kindlin-3) to regulate this complex. Therefore, the altered interaction would be responsible for significant change in the activity of this complex in cancer cells. Kindlin2 is proven to be a key player in TGBRI-SMAD3 interaction through direct binding^48^, both of which are upregulated in cancer. Interaction analysis shows a higher affinity of kindlin2 mutants for TGFBRI and SMAD3, plausibly promoting TGFβ-signaling in kindlin2-mutant cancer-patients. Another kindlin2 interactor SEPTIN11 is also associated with chronic neutrophilic leukaemia. Due to mutations, a decreased possibility of kindlin2-SEPTIN11 interaction and further co-alteration in pan-cancer cohort has been observed. We see a significant co-alteration of kindlin-2 interactor MAPK, which was previously linked to chromosomal/microsatellite instability through COAD, GBM, and PAAD signaling pathways. Another cancer signature, telomere instability has been recently proved to be dependent on ECM stiffness and mechanosensitivity^49^. Interestingly, kindlin3 interactors EXOSC10 and Kindlin1 interactor SKIV2L are linked to telomere maintenance. Alterations of these two proteins in cancer might therefore associate kindlin-mechanochemical signaling with cancer-specific telomeric instability.

Kindlins are also important for connecting mechanical environment to cellular metabolism. Previous reports suggest involvement of kindlin2 in amino acid biosynthesis and HIF-1α induced tumor growth in hypoxic condition.^16,50^ Our analysis suggests, a role of kindlin2 in GSK3 mediated inhibition of TCA cycle either via AKT1 or via CDC42 (Fig. S49). Predicted kindlin2 interactor PFKM is important in upregulating glycolysis. Although, specific role of this interaction is unknown, we can see a probable decrease in kindlin2-PFKM interactions in cancer. From this, we might hypothesize a PFKM-inhibiting role of kindlin2, which is altered in cancer thereby increasing glycolysis. This downregulation of TCA cycle and higher anaerobic glycolysis is a signature of cancer cell proliferation, growth, and chemo-resistance^51^. In pathway-specific alteration analysis, we see a combined effect of all the kindlins, especially kindlin2 and 3, in activation of EMT accompanied by inhibition of DNA damage response. In addition, we see a role of kindlins in cell-cycle arrest, a common signature during EMT associated to increased ribosome biogenesis^52^. Notably, we observe that kindlin1 alterations are correlated with negative regulation of cell-cycle progression, DNA-damage response, apoptosis as well as EMT. This might be because of patient-specific role rather than an overall scenario. A second possibility indicates its role in maintaining the temporal regulation of drug efflux, invasion, proliferation and stemness in different time points of EMT^53^. Therefore, it becomes important to further elucidate the spatiotemporal role of kindlins in cancer progression and metastasis.

In summary, this analysis represents a pan-cancer scenario of how mechanosensitive adaptor-proteins like kindlins connect environmental mechanical cues and translate them to intracellular chemical events to favour cancer. This alteration seems an effect of altered intrinsic structural properties of kindlins – stability, flexibility, hydrophobicity, signal transmission property, and chemical properties – specific interactions, phosphorylation, and loss of functional domains, among others. A majority of these observations suggests correlations and not causality, hence demand further experimental mechanistic validations using structural, biochemical, and cellular approaches to find the exact causations in mechanistic details. However, it is certain that kindlins act as ‘super adaptors’ rendering many homeostatic and pathophysiological pathways and therefore, can be a good target for therapeutic strategy against cancer.

## Methods

### Dataset Curation

Preliminary genomic data for FERMT1, FERMT2, and FERMT3 was acquired from The Cancer Genome Atlas’ (TCGA, https://www.cancer.gov/tcga), pan-cancer analysis of whole genomes (PCAWG)^23^ study and processed. Cancer-associated somatic mutations were obtained from the Catalogue of Somatic Mutations in Cancer (COSMIC)^54^, and a dataset of multiple mutations per donor was curated from the International Cancer Genome Consortium (ICGC)^55^ database.

### Analysis of Expression Levels

Donor-specific mRNA expression levels were obtained from the ICGC^55^ Data Portal for donors who possessed sequencing-based gene expression and array-based gene expression data. In addition to these donors across the Kindlins family (n=1045), mutation-specific expression data for genomic alterations (n=268; 18 cancer cohorts) in the non-coding regions was also obtained from the ICGC^55^ dataset. For both analyses, initial data was filtered for donors who possessed high and low functional impact mutations only, leaving out donors with mutations of unknown impact.

Furthermore, expression analysis across 33 cancer cohorts matching TCGA normal and GTEx data (https://gtexportal.org/home/) was performed using data from GEPIA2^56^, employing the one-way analysis of variance (ANOVA) test along with a q-value cut-off of 0.01 and a log_2_(TPM + 1) scale (TPM, Transcripts per million). Lastly, mRNA expression levels were analysed as a function of two clinical attributes – Tumor Stage (AJCC) and Metastasis Stage Code (AJCC) – using data from the PCAWG study^23^.

For all genes outlined as part of the direct and indirect physical interactome of FERMT1, FERMT2, and FERMT3 (using BioGRID^4.4^, https://thebiogrid.org/)^57^, we performed co-alteration of expression analysis on the PCAWG^23^ data to characterise their impact and predict their involvement in associated pathways and downstream functions.

For cancer and non-cancer patients, protein expression data was obtained and analysed using the Genomic Data Commons’ (GDC) Clinical Proteomic Tumor Analysis Consortium (CPTAC, NCI/NIH) database that incorporates the TARGET (Therapeutically Applicable Research to Generate Effective Therapies) and TCGA libraries on cProSite (https://cprosite.ccr.cancer.gov/). Consequently, relevant microRNAs and their expression was analysed in the context of cancer via systematic analysis of primary literature and text-mining of high-throughput experimental data on miRDB (http://www.mirdb.org/) and miRCancer (http://mircancer.ecu.edu/).

### Protein expression data analysis

Pan-cancer protein expression (mass-spectrometry) data were obtained from CPTAC dataset for 11 types of cancer. We obtained total 1272 tumor sample data and 808 tumor-adjacent tissue samples. Stomach cancer data was omitted due to no record of Kindlin expression in tumor adjacent samples. Protein expression data were collected as relative protein abundance in TMTlog2 ratio (TMT, Tandem mass tag) Patient-specific protein-phosphorylation data were collected from the same dataset. Phosphorylation analysis was available for 1272 tumor samples and 782 tumor adjacent samples. Phosphorylation alteration was calculated as:

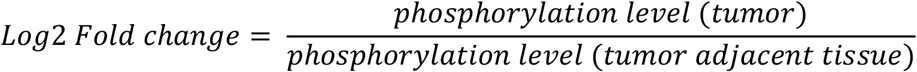

### Survival Analysis

Kaplan Meier (KM) plots for expression-specific overall and disease-free survival were performed using curated data originally taken from TCGA/ICGC cohort using cBioportal (https://www.cbioportal.org/) and GEPIA2 (http://gepia2.cancer-pku.cn/)^56^ across 33 different cancer types. Overlapped samples (n=193) were excluded from the analysis. High mRNA expression data for which we have obtained were: 2405 (FERMT1), 2375 (FERMT2), and 2376 (FERMT3). Similarly low mRNA expression data were available for 2380 (FERMT1), 2376 (FERMT2), and 2376 (FERMT3) samples. Data from the PCAWG^23^ study and TCGA data (n=2583) for which status was available, was employed to curate mutation status-specific overall survival KM plots.

### Mutated variant analysis

COSMIC^54^ data were first collected individually for FERMT1, FERMT2, and FERMT3. Kindlin Point mutated samples were collected from curated dataset containing PCAWG^23^, TCGA, ICGC^55^ data. Overlapped samples were filtered out which obtained a total 981 samples. Based on consensus transcript sequences for FERMT1, FERMT2, and FERMT3, multiple mutations per transcript (having same transcript ID within individual sample ID) were also recorded (n=38). These multiple-nucleotide variants (MNV) were further assessed and filtered manually applying MNV calling criteria by Wang et al.^58^ Functional impact i.e., loss-of-function or neutral, of the single nucleotide mutations were obtained using SIFT^59^.

### Cancer-Specific Mutational Stability Characterisation

The monomeric structures for FERMT1 (Q9BQL6), FERMT2 (Q96AC1), and FERMT3 (Q86UX7) were derived from AlphaFold^60^. Dimers for each of them were prepared by performing homology modelling on trRosetta^61^. Structural homology was further checked using POSA^62^. Cancer-specific substitution mutations and frameshift insertion-deletion mutations were incorporated using mutagenesis tool of PyMOL (v. 2.5.2, The PyMOL Molecular Graphics System, Version 1.2r3pre, Schrödinger, LLC) and trRosetta^61^, respectively.

Perturbations in the dynamics and stability of FERMT1, FERMT2, and FERMT3 dimers as a function of cancer-specific mutations were characterized employing an Elastic Network Contact Model – based normal mode analysis (ENCoM – based coarse-grained NMA)^63^ to compute predicted changes in folding free energy (ΔΔG) and change in vibrational entropy (ΔΔS). The ENCoM – based NMA approach takes into consideration the distinct nature and consequent effects of specific amino acids on the dynamics of the structure^63^. Moreover, the involvement of vibrational normal modes and entropic analysis within the NMA method represents a novel approach to characterising protein structure dynamics and the effect of mutations^63^.

### Preparation of Insertion-deletion Mutant Structures

FASTA protein sequences for FERMT1, FERMT2, and FERMT3 were obtained from NCBI following the correct amino acid sequence length within the *<u>Homo sapiens</u>* collection of transcripts. Each of the sequences were queried against translated nucleotide databases using the TBLASTN tool’s BLOSUM62 matrix adjusted for conditional compositional score, and existence:11 extension:1 gap cost. Resulting sequences producing significant alignments were sorted by highest percentage identity with their respective query sequences. With the three best matches against FERMT1, FERMT2, and FERMT3, insertion-deletion mutations obtained from COSMIC^54^ were incorporated within each nucleic acid sequence. These mutated sequences were translated to their respective peptide sequences using EMBOSS Transeq. For each nucleic acid sequence, we employed frame 1, and standard code parameters. The protein sequences obtained were screened and submitted on the trRosetta server^61^ to predict insertion-deletion mutant structures. The structures with the highest homology modelling score were selected for downstream analysis.

### Structural Analysis of Cancer-specific Variants

The characteristic deviation of stability (ΔΔG) and flexibility (ΔΔS) of cancer-specific mutant variants with reference to the wild type structures was performed using the ENCoM – based NMA method^63^, allowing for a more accurate representation of intramolecular interactions and the prediction of consequent effects on conformational stability and molecular flexibility^18^.

In our analysis ΔΔG was represented as:

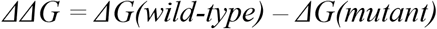

And thus, ΔΔG = (+) ve means stabilization and ΔΔG = (-) ve means destabilization; ΔG, free energy change.

Similarly, ΔΔS was calculated as:

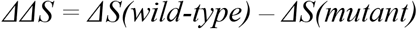

Here, ΔΔS = (+) ve means increased flexibility and ΔΔS = (-) ve means decreased flexibility. ΔS, change in vibrational entropy.

Preliminary data collected from all cancer-associated somatic mutations via this method was then filtered to screen for mutations that corresponded to a ΔΔG_ENCoM_ value of ≥ +1.24 and ≤ -1.24 for further analysis (as very highly stabilizing and destabilizing mutants).

Furthermore, the B-factor (generally termed as the Debye-Waller Factor) was computed using the following formula to predict the flexibility of each amino acid residue within the respective chains, where ΔR^2^ indicates the oscillation of every residue from its equilibrium position^64^.

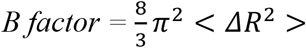

Following NMA, the deformation energies and eigenvalues of multiple low-frequency normal modes were computed using the below-mentioned formula for all wild type and mutant variants to measure the degree of collectivity for each mode, and consequently predict the extent of displacement of large regions of the protein structures^64^ as:

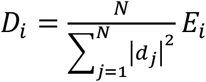

Here, E_i_ is the energy contribution of atom *i*, *d_j_* is the displacement vector of atom *j*, and N is the number of atoms in the molecule.

The alterations of signal reception and transmission properties in Kindlin mutants were computed using Markov’s chain model in DynOmics^65^. We used Phos3D^66^ to examine the effects on phosphorylation of experimental validated kindlin phosphorylation sites using 3D pdb coordinates.

Hydrophobic (Isoleucine, Leucine, Valine) clusters predicted to confer stability to proteins were visualized based on the Contacts of Structural Units (CSU) algorithm^67^, creating a Fibonacci-styled residue matrix and a subsequent graph plotting hydrophobic clusters for all mutant structures. Electrostatic interactions, which conferring stability and structural compactness to proteins, were computed by predicting the extent of charge segregation (κ) and the fraction of charged residues in a sequence (FCR) for all mutant variants^67^.

### Gene Ontology and Pathway Enrichment Analysis

To locate associated cellular compartments, pinpoint related molecular functions, and outline linked cellular processes, we performed gene ontology using BioGRID^4.457^ to profile the high-throughput physical interactome of FERMT1, FERMT2, and FERMT3 respectively. The interactome components were restricted to proteins found in *<u>Homo sapiens</u>* only. Using STRING (v. 11.5)^68^, the full STRING network type with a medium confidence, minimum interaction score of 0.400 was employed to indicate both functional and *direct* physical protein associations.

Ensembl ID sets for FERMT1, FERMT2, and FERMT3 were used to predict and perform pathway enrichment analysis. Gene ontology (GO) molecular functions were computed as a function of fold enrichment. The Kindlin-associated mechanochemical proteins were identified by a meta-analysis from text-mined articles. Hallmark genes were identified from MsigDB (http://www.gsea-msigdb.org/gsea/msigdb/index.jsp). The co-alteration analysis was performed for these mechanosensitive proteins and hallmark genes and the extent of co-alterations were represented using the term mean co-alteration dynamics which we have defined as:

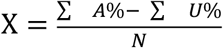

Here, X= mean co-alteration dynamics of a gene set; A% = percentage of samples altered; U% = percentage of samples unaltered; N= number of genes in set.

Pathway alterations were applying gain-of-function and loss-of-function of pathway associated genes when Kindlin is altered for checking activation and inhibition respectively. Global percentage of pathway activity for a particular pathway and for a particular Kindlin was calculated as:

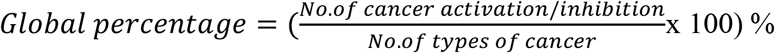

The detailed methodology EMT-specific signalling modules and biological pathways were analysed using EMTome^69^.

### Partner-Specific Interaction Analysis

Direct physical protein associations filtered from STRING (v. 11.5)^68^ were docked with their respective partner Kindlin employing a hybrid algorithm of template-based and *ab initio* free modelling and docking^28^. The computed docking scores, and subsequent curation of fold change in docking energy relative to the wild type enabled the prediction of the ligands’ and the receptor’s binding affinity for all cancer mutation variants. The effects of cancer-associated substitution mutations on the monomeric Kindlin’s ability to form a dimer structure were also predicted via symmetric C2 docking^70^ of the monomer using the same algorithm. The binding affinity of dimerization was calculated as:

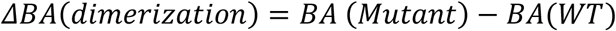

BA = binding affinity, WT= wild type. Binding affinities are represented in Kcal/mol unit. If ΔBA_dimerization_ = (+) ve, it suggests a destabilization and unfavourable while (-) ve ΔBA_dimerization_ suggests a stabilization and is favourable.

The fold change in interactor specific interaction affinity was calculated as:

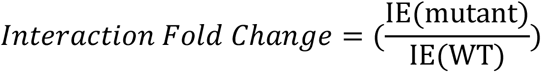

WT=wild type; IE=mutant; Fold change of 0.1 = 24 KJ/mol for FERMT1, 22.7 KJ/mol for FERMT2, 23.1 KJ/mol for FERMT3. Interaction fold change>1, increased interaction; Interaction fold change<1, decreased interaction; Interaction fold change=1, affinity same as the wild-type.

Further details of datasets used, copy number variation analysis, methylation analysis, classification of stabilizing and destabilizing mutants, meta-analysis of Kindlin-associated mechanochemical signalings, calculation of signal transduction properties and cancer-specific pathway analysis are given in Supplementary methodology.

### Statistical Analysis

Statistical analyses were performed using R version-4.2.1, R version-4.0.3 (http://cran.r-project.org/) and OriginPro (https://www.originlab.com/). Routine statistical tests were employed to validate statistical significance. This includes the Log rank p-test and Cox proportional-hazards model for KM plots with 95% confidence intervals, Kruskal-Wallis for comparing the survival corresponding to mutations in different Kindlins. While comparing two groups, we used unpaired two-tailed t-tests. One-Way ANOVA and Bonferroni post-hoc test was employed to calculate statistical significance within cancer subtypes of the same cancer type (cut-off ≤ 0.05). Significance cut-off for p-value was taken <0.05. Z-scores to study gene expression alterations were considered for samples with a cut-off of ±1.96 corresponding to p-value <0.05. For significant data of fold enrichment in –Log_10_ (FDR) scale, cut-off for FDR was taken at less than 0.05. Spearman’s Coefficient correlation was used to validate correlation between two non-parametric variables, where +1 indicates highest correlation, -1 indicates anti-correlation, and 0 indicates no correlation. The nature of data distribution was verified and represented in terms of the Shapiro-Wilk normality test and skewness. Bars, wherever used, indicate standard error of mean.

## Data availability

Cancer patients genomics data is available in the following sites: PCAWG data (https://dcc.icgc.org/pcawg), ICGC data (https://dcc.icgc.org/), TCGA (https://www.cancer.gov/about-nci/organization/ccg/research/structural-genomics/tcga). Cancer patient-specific proteomics data are available from CPTAC (https://proteomics.cancer.gov/data-portal, https://proteomic.datacommons.cancer.gov/pdc/). Mutation data is available from COSMIC (https://cancer.sanger.ac.uk/cosmic). Normal tissue gene expression data is available at GTEx (https://gtexportal.org/home/). miRNA data is available at miRDB (http://www.mirdb.org/) and their cancer-specific alteration data (TCGA set) is available at miRCancer (http://mircancer.ecu.edu/). Mutation-specific structural data, phosphorylation data and Mechano-chemical protein meta-analysis generated data are available in the supplementary information file. Other data generated during this study are available from the corresponding authors upon reasonable request.

## Code availability

Codes generated during this study are available from the corresponding authors upon reasonable request.

## Author Contributions

D.C and S.H designed the project. D.C handled the COSMIC dataset. A.M handled the TCGA dataset. R.B and S.W handled the PCAWG dataset. D.C, A.M, R.B, and S.W analysed the data. D.C, A.M, R.B, and S.W performed the experiments. D.C, A.M, R.B, and S.W analysed the experimental data. D.C, R.B and A.M built the codes. D.C, A.M, R.B, and S.W performed the meta-analyses. D.C, A.M, R.B, and S.C drafted the manuscript with input from all the authors. D.C, A.M, R.B and S.W participated in figures preparation. S.H and S.C reviewed the manuscript. All authors have read and approved the final version of the manuscript.

## Acknowledgement

The results published here are based upon data generated by the TCGA, ICGC, PCAWG, COSMIC and ENCODE project. We thank Ashoka University and the Mphasis foundation for support and funding. S.H. thanks DBT Ramalingaswami Fellowship and DST SERB Core Research Grant for funding. We thank Dr. Sougata Roy (Ashoka University), for sharing valuable suggestions.

## Supplementary Information

### Cancer Name Abbreviations

ACC: Adrenocortical Cancer
BLCA: Bladder Urothelial Carcinoma
BRCA: Breast Invasive Carcinoma
CESC: Cervical Squamous Cell Carcinoma and Endocervical Adenocarcinoma
CHOL: Cholangio Carcinoma
COAD: Colon Adenocarcinoma
DLBC: Lymphoid Neoplasm Diffuse Large B-cell Lymphoma
ESCA: Esophageal Carcinoma
GBM: Glioblastoma Multiforme
HNSC: Head and Neck Squamous Cell Carcinoma
KICH: Kidney Chromophobe
KIRC: Kidney Renal Clear Cell Carcinoma
KIRP: Kidney Renal Papillary Cell Carcinoma
LAML: Acute Myeloid Leukemia
LGG: Brain Lower Grade Glioma
LIHC: Liver Hepatocellular Carcinoma
LUAD: Lung Adenocarcinoma
LUSC: Lung Squamous Cell Carcinoma
MESO: Mesothelioma
MSI: Microsatellite Instability
MSS: Microsatellite Stable
OV: Ovarian Serous Cystadenocarcinoma
PAAD: Pancreatic Adenocarcinoma
PCPG: Pheochromocytoma and Paraganglioma
PRAD: Prostate Adenocarcinoma
READ: Rectum Adenocarcinoma
SARC: Sarcoma
SKCM: Skin Cutaneous Melanoma
STAD: Stomach Adenocarcinoma
TGCT: Testicular Germ Cell Tumours
THCA: Thyroid Carcinoma
THYM: Thymoma
UCEC: Uterine Corpus Endometrial Carcinoma
UCS: Uterine Carcinosarcoma
UVM: Uveal Melanoma

### Gene Descriptions (Direct Interactors of Kindlins)

**FERMT1,** Fermitin family homolog 1 or Kindlin-1 protein is encoded by this gene, which plays a role in cell adhesion and integrin activation.

**FERMT2,** encodes a protein called Fermitin family homolog 2 or Kindlin-2 that enhances integrin activation mediated by talins and are required for the assembly of focal adhesions.

**FERMT3,** gene coding for a protein called Fermitin family homolog 3 or Kindlin-3, which plays a crucial role in cell adhesion in hematopoietic cells.

**SkiV2L**, Superkiller Viralacidic Activity-2 homolog is a protein-coding gene that codes for an enzyme in humans called Helicase SKI2W, which is a helicase that has ATPase activity and is part of the SKI complex.

**ITGB1**, gene encoding for a protein called Integrin beta-1, which is involved in the motility of the endothelial cell. It also acts as a receptor for various other proteins such as fibronectin and collagen.

**PARVA**, gene coding for a protein called Alpha-parvin, which is important in sarcomere organisation and smooth muscle cell contraction and angiogenesis.

**TTC37,** gene encoding for a protein called Tetratricopeptide Repeat Protein 37 involved in the SKI complex of proteins to form an exosome that is thought to break down abnormal RNA molecules in the cytosol.

**ILK**, codes for a protein called Integrin-linked Protein Kinase that interacts with integrins and regulates signal transduction.

**PARVB**, protein-coding gene for Beta-parvin, a protein that binds to ILK and plays a role in integrin signalling and cytoskeletal reorganisation.

**LIMS1**, gene coding for a protein called LIM and Senescent Cell Antigen-like-containing Domain Protein 1, which is responsible for linking beta-integrins to the actin cytoskeleton.

**LSM8**, encodes a protein called U6snRNA-associated Sm-like protein LSm8, which is part of a complex, involved in spliceosome assembly.

**EXOSC10,** protein-coding gene that encodes a protein called Exosome component 10, which is a part of the RNA exosome complex.

**PFKM**, gene encoding a protein called ATP -dependent 6-phosphofructokinase, muscle type, which is a protein that plays a catalytic role in a step-in glycolysis.

**SEPTIN9**, encodes a protein named Septin-9, which is speculated to play a role in cytokinesis.

**SEPTIN11**, codes for a protein called Septin-11, which may potentially have a role to play in the process of cytokinesis.

**PICALM**, gene coding for Phosphatidylinositol-binding Clathrin Assembly Protein, which is essential in clathrin-mediated endocytosis.

**VCL**, encodes Vinculin, a protein that is an F-actin binding protein and is therefore involved in cell-matrix and cell-cell adhesions. Vinculin is also involved in the mechanosensitive ability of the E-cadherin complex.

**PXN**, gene coding for Paxillin, a mechanosensitive cytoskeletal protein playing a crucial role in focal adhesion sites.

**MAPK1**, encoding a protein called Mitogen-activated Protein Kinase 1, which is an important component of the MAP kinase signal transduction pathway.

**SMAD3,** encodes for a protein called Mothers against decapentaplegic homolog 3, which is an intercellular signal transducer, and transcriptional modulated that is regulated by other proteins and kinases.

**TGF**β**R1**, protein-coding gene that forms a heteromeric complex with type II TGFβ receptors when bound to TGFβ, facilitating the TGFβ signalling transduction from the cell surface to the cytoplasm.

**HIF1A,** encodes the alpha subunit of the transcription factor hypoxia-inducible factor-1, a heterodimer composed of an alpha and beta subunit that plays an essential regulatory role in multiple cellular processes.

**EGFR**, protein-coding gene that is widely recognized for its role in cancer, epidermal growth factor receptor is a transmembrane glycoprotein in the protein kinase superfamily that serves as a receptor for the epidermal growth factor family members.

### Glossary Terms

**FERM Domain**, the FERM domain (4.1 protein, Ezrin, Radixin, and Moesin) is approximately 300 amino acids long and present in a large number of proteins involved with actin cytoskeleton and membrane interactions and plasma membrane localisation.

**Mechanochemical Signalling**, a process involving the transduction of mechanical cues into biochemical signals because of force-induced conformational modulation.

**B-factor**, also known as Debye-Waller factor, B-factor analysis of protein structures give information about their atomic displacement parameters, or the disorder of atoms from their equilibrium or average (root mean square) position or temperature. Higher B-factor indicates higher flexibility, as in greater displacements from the mean, which could also be indicative of a lower electron density. The formula can be used to predict the flexibility/rigidity of individual atoms, specific or entire regions, and even side chains of a protein. The model used in this paper to predict B-factor of mutated protein structures used the following equation:

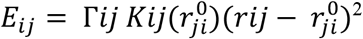

Where:

*r_ij_* → distance between residues *i* and *j*
*r_ij_*^0^ → equilibrium value
Γ*_ij_* → Kirchhoff connectivity matrix
*K_ij_*(*r_ij_*^0^)→ stiffness force constant

**Copy Number Variation (CNV),** a circumstance wherein there is a variation in the number of copies of specific segment(s) of DNA within an individual.

**Kappa (κ) Value**, a parameter to compute the sequence-specific degree of charge segregation, which is dependent on the fraction of charged residues (FCR) in a sequence. This indicates the role of salt-bridge interactions in protein compactness.

**Fig.S1:**
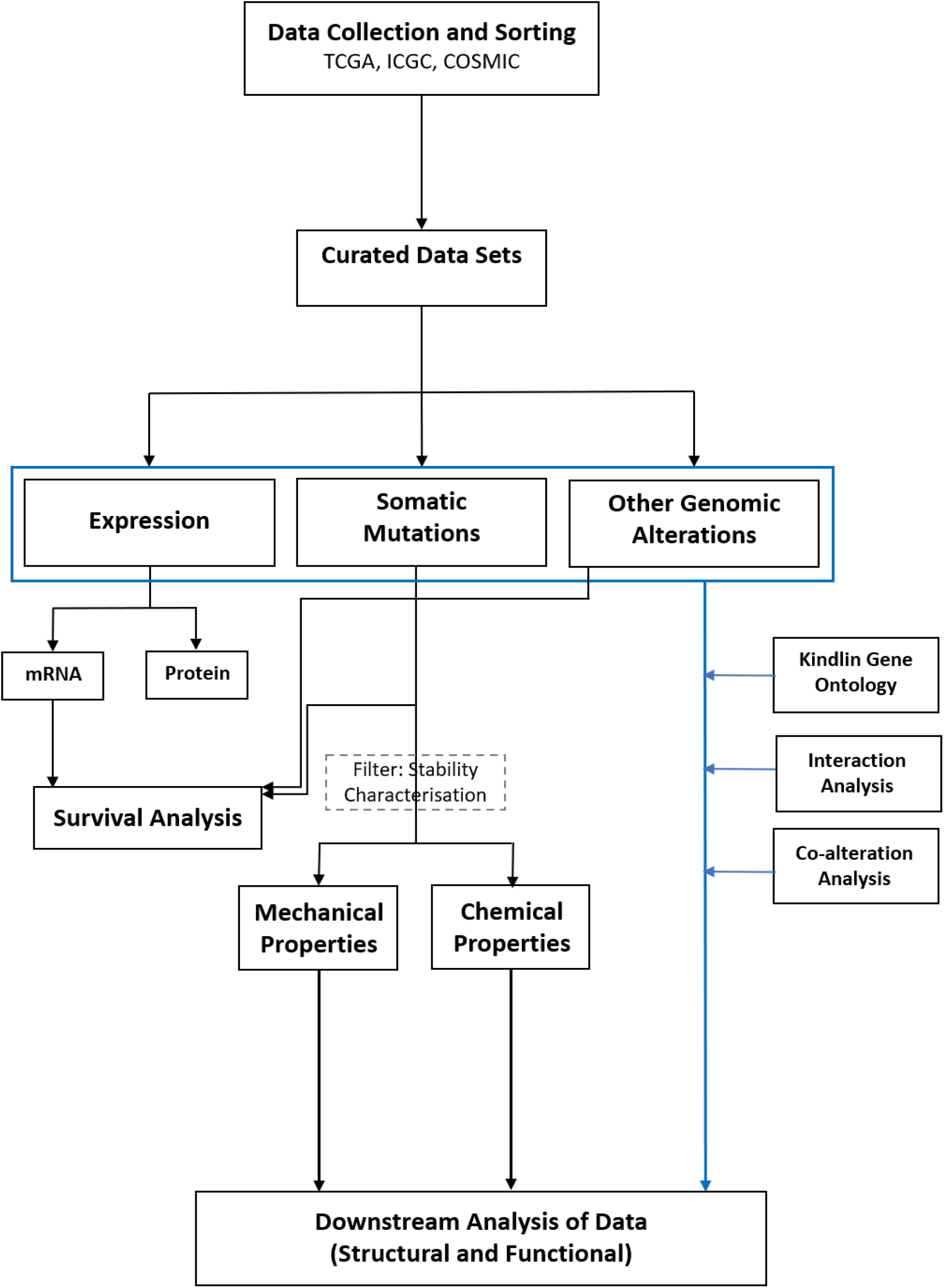
Dataset Curation Pipeline Facilitating Downstream Analysis. Multi-omics data for all kindlin family proteins was collected from respective databases and curated in an assay-specific manner – expression, somatic mutations, and genomic alterations – to perform relevant assays that form the constituent components and basis of subsequent structural and functional downstream analyses.

**Fig.S2:**
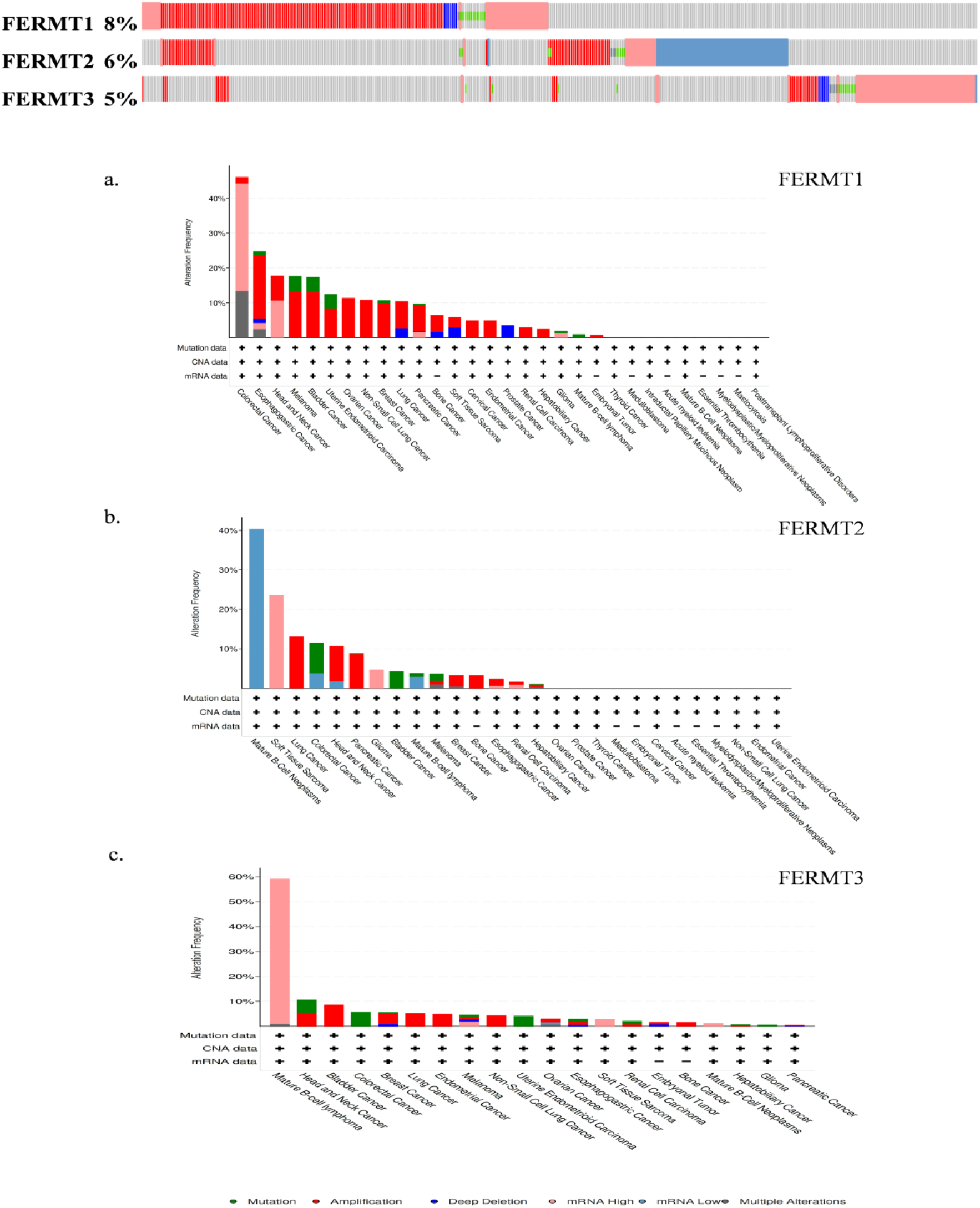
Pan-cancer Alterations of All Kindlin Family Proteins. a. Percentage genetic alterations in all three kindlin family proteins across the PCAWG cohort. Red boxes, amplification; red-bordered grey boxes, high mRNA expression; blue-bordered grey boxes, low mRNA expression; blue boxes, deep deletion; green boxes, mutation; olive boxes, truncating mutations; grey boxes, no alterations. (b-d). Cancer-type-specific prevalence of genomic alterations in FERMT1, FERMT2, and FERMT3, respectively, along with specifications regarding the nature of data profiled for each cancer cohort. Green, mutation; red, amplification; blue, deep deletion; sky blue, mRNA low; peach, mRNA high; grey, multiple alterations; (+) sign, profiled for; (-) sign, not profiled for.

**Fig.S3:**
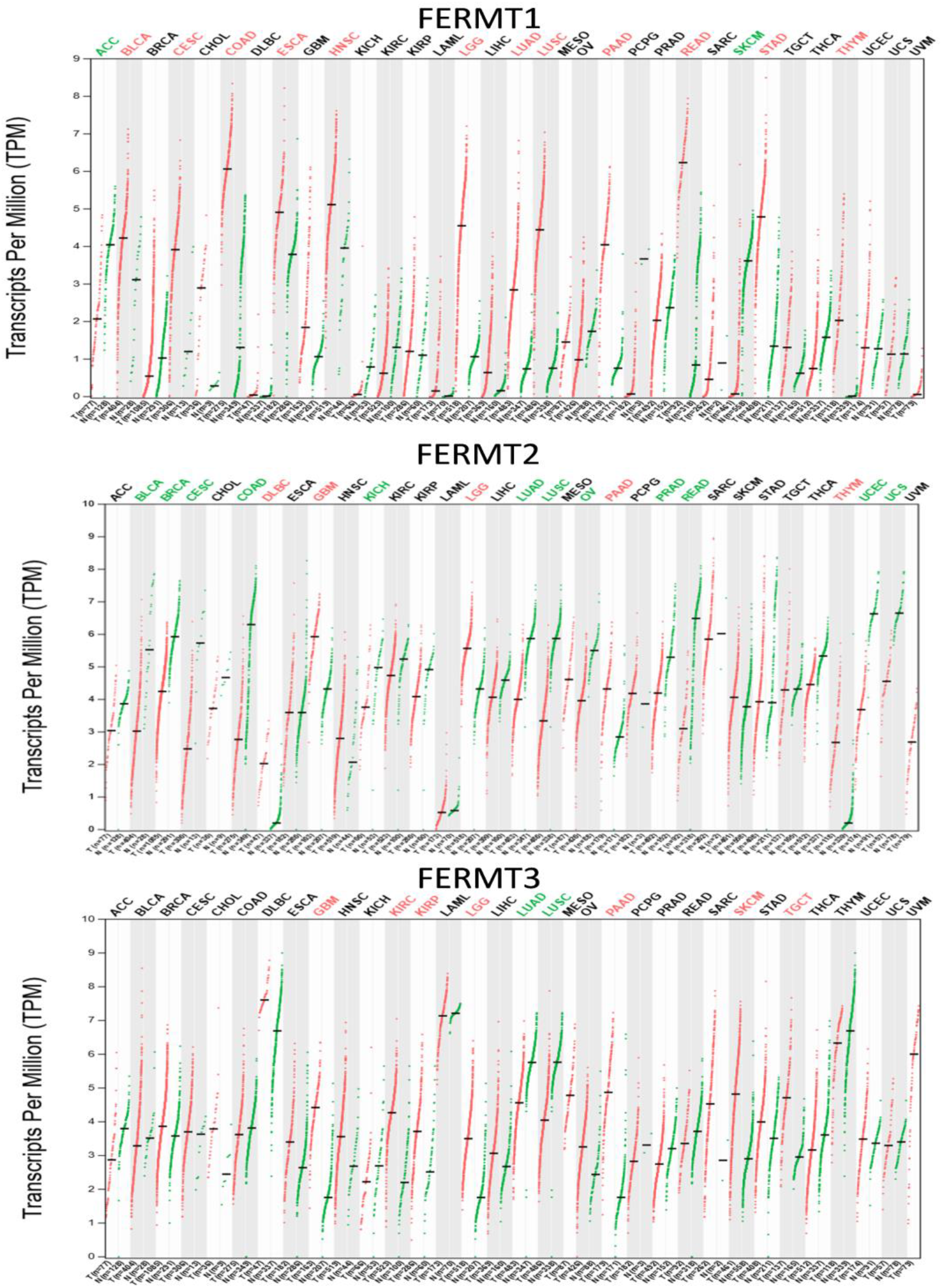
Gene Expression Profiles of All Kindlin Family Proteins Across Tumor and Normal Tissue Samples in Multiple Cancer Cohorts as Dot Plots. The x-axis represents the number of tumor samples in red (T) and the number of non-tumor samples in green (N), while the y-axis represents transcripts per million (TPM) in the Log_2_(TPM +1) scale. The upper horizontal margin indicates abbreviated forms of every profiled tumor cohort; red and green represent statistical significance in favour of the tumor and normal sample group, respectively. Black shows statistically non-significant data, and the black dash on each vertical plot represents the mean value for that range.

**Fig.S4:**
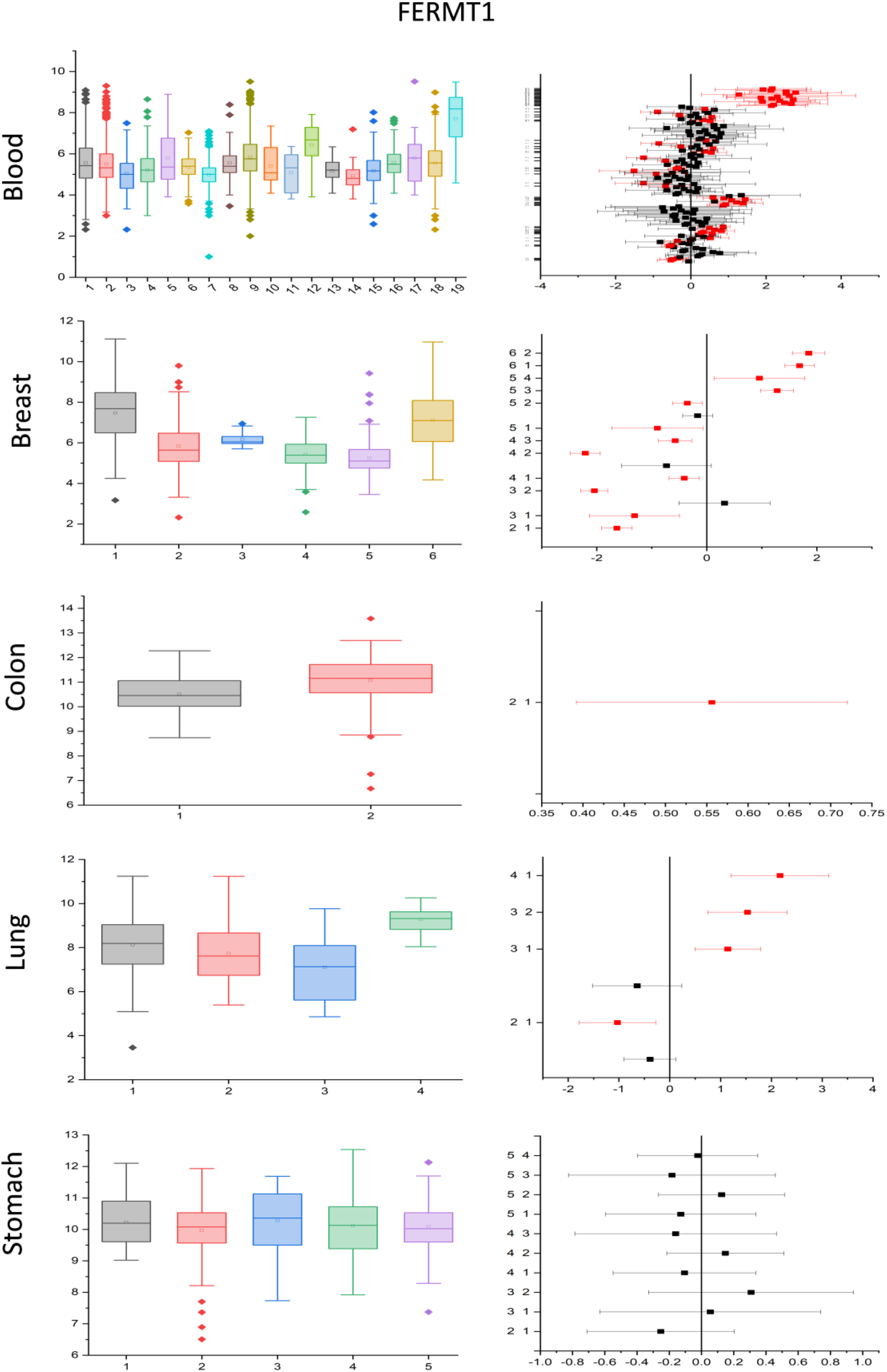
Subtype-Specific Gene Expression Patterns of FERMT1 across Five Tumor Types. For each tumor type, the boxplots depict subtype-specific gene expression, with different colours representing each subtype. The box indicates the region where majority of the values reside. The range of data is depicted by error bars for each cancer subtype. The points represent outliers for each dataset. For blood cancer – 1, acute lymphoblastic leukaemia; 2, acute myeloid leukaemia; 3, B-cell acute lymphoblastic leukaemia; 4, B-cell lymphoma; 5, Burkitt lymphoma; 6, chronic Hodgkin lymphoma; 7, chronic lymphocytic leukaemia; 8, chronic myeloid leukaemia; 9, diffuse large B-cell lymphoma; 10, follicular lymphoma; 11, hairy cell leukemia; 12, Hodgkin lymphoma; 13, Mantle cell lymphoma; 14, mixed lineage leukaemia; 15, multiple myeloma; 16, myelodysplastic syndrome; 17, splenic marginal zone lymphoma; 18, T-cell acute lymphoblastic leukaemia; 19, T-cell lymphoma. For breast cancer – 1, basal; 2, HER2; 3, luminal; 4, luminal A; 5, luminal B; 6, TNBC. For colon cancer – 1, MSI; 2, MSS. For lung cancer – 1, non-small cell lung cancer; 2, non-small cell lung cancer adenocarcinoma; 3, non-small cell lung cancer large-cell carcinoma; 4, non-small cell lung cancer small-cell carcinoma. For stomach cancer – 1, EMT; 2, MSI; 3, MSS; 4, MSS/TP53-; 5, MSS/TP53+. For subtypes within each cancer, one-way ANOVA analysis was performed to verify the statistical significance, and their mean differences corrected via Bonferroni post-hoc test at p ≤ 0.05. In all graphs to the right, the red markers correspond to subtypes with statistically significant difference in mean value, whereas the black markers indicate statistical non-significance for each corresponding cancer subtypes assigned as box-whisker plot.

**Fig.S5:**
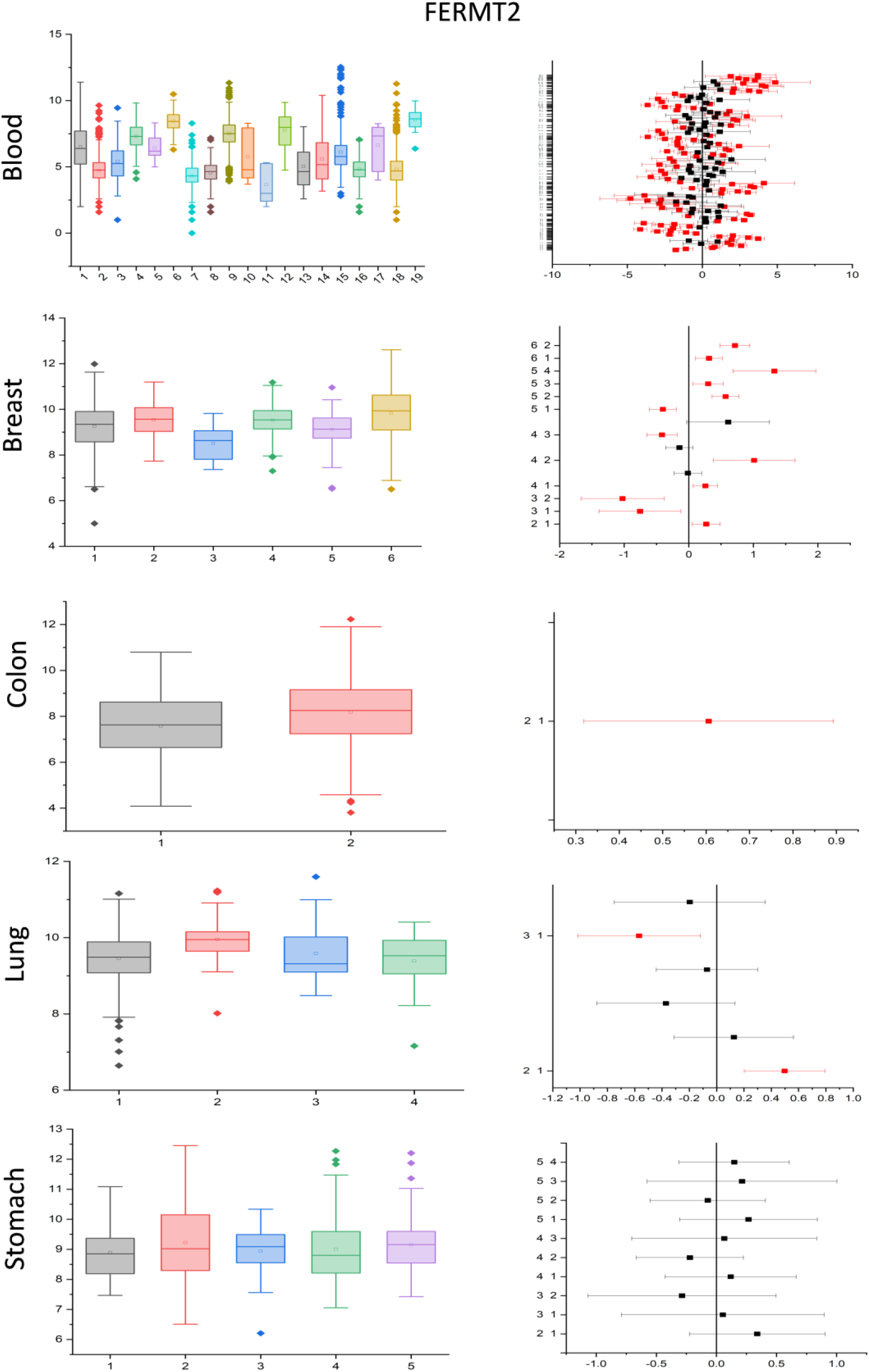
Subtype-Specific Gene Expression Patterns of FERMT2 across Five Tumor Types. For each tumor type, the boxplots depict subtype-specific gene expression, with different colours representing each subtype. The box indicates the region where majority of the values reside. The range of data is depicted by error bars for each cancer subtype. The points represent outliers for each dataset. For blood cancer – 1, acute lymphoblastic leukaemia; 2, acute myeloid leukaemia; 3, B-cell acute lymphoblastic leukaemia; 4, B-cell lymphoma; 5, Burkitt lymphoma; 6, chronic Hodgkin lymphoma; 7, chronic lymphocytic leukaemia; 8, chronic myeloid leukaemia; 9, diffuse large B-cell lymphoma; 10, follicular lymphoma; 11, hairy cell leukemia; 12, Hodgkin lymphoma; 13, Mantle cell lymphoma; 14, mixed lineage leukaemia; 15, multiple myeloma; 16, myelodysplastic syndrome; 17, splenic marginal zone lymphoma; 18, T-cell acute lymphoblastic leukaemia; 19, T-cell lymphoma. For breast cancer – 1, basal; 2, HER2; 3, luminal; 4, luminal A; 5, luminal B; 6, TNBC. For colon cancer – 1, MSI; 2, MSS. For lung cancer – 1, non-small cell lung cancer; 2, non-small cell lung cancer adenocarcinoma; 3, non-small cell lung cancer large-cell carcinoma; 4, non-small cell lung cancer small-cell carcinoma. For stomach cancer – 1, EMT; 2, MSI; 3, MSS; 4, MSS/TP53-; 5, MSS/TP53+. For subtypes within each cancer, one-way ANOVA analysis was performed to verify the statistical significance, and their mean differences corrected via Bonferroni post-hoc test at p ≤ 0.05. In all graphs to the right, the red markers correspond to subtypes with statistically significant difference in mean value, whereas the black markers indicate statistical non-significance for each corresponding cancer subtypes assigned as box-whisker plot.

**Fig.S6:**
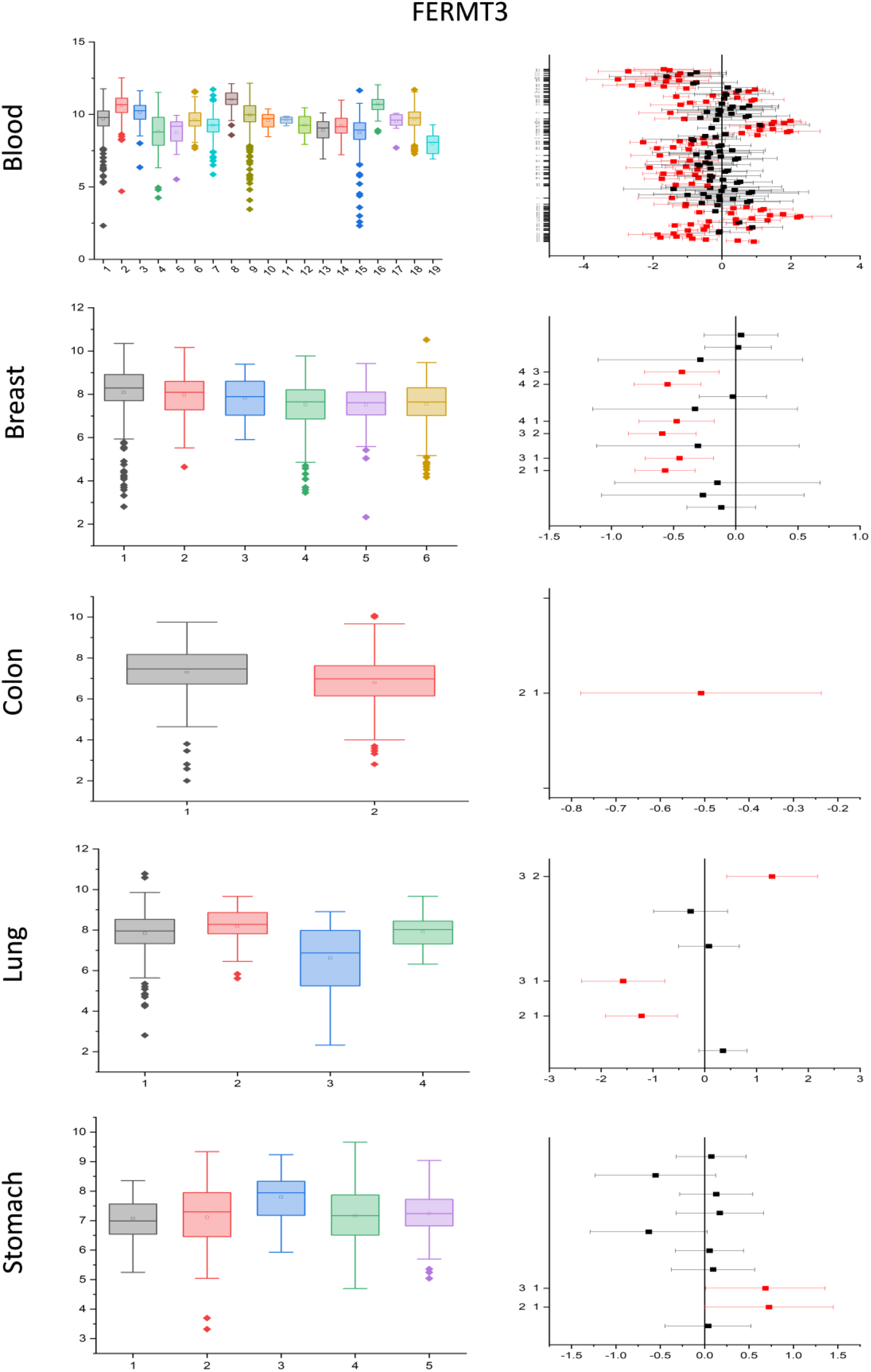
Subtype-Specific Gene Expression Patterns of FERMT3 across Five Tumor Types. For each tumor type, the boxplots depict subtype-specific gene expression, with different colours representing each subtype. The box indicates the region where majority of the values reside. The range of data is depicted by error bars for each cancer subtype. The points represent outliers for each dataset. For blood cancer – 1, acute lymphoblastic leukaemia; 2, acute myeloid leukaemia; 3, B-cell acute lymphoblastic leukaemia; 4, B-cell lymphoma; 5, Burkitt lymphoma; 6, chronic Hodgkin lymphoma; 7, chronic lymphocytic leukaemia; 8, chronic myeloid leukaemia; 9, diffuse large B-cell lymphoma; 10, follicular lymphoma; 11, hairy cell leukemia; 12, Hodgkin lymphoma; 13, Mantle cell lymphoma; 14, mixed lineage leukaemia; 15, multiple myeloma; 16, myelodysplastic syndrome; 17, splenic marginal zone lymphoma; 18, T-cell acute lymphoblastic leukaemia; 19, T-cell lymphoma. For breast cancer – 1, basal; 2, HER2; 3, luminal; 4, luminal A; 5, luminal B; 6, TNBC. For colon cancer – 1, MSI; 2, MSS. For lung cancer – 1, non-small cell lung cancer; 2, non-small cell lung cancer adenocarcinoma; 3, non-small cell lung cancer large-cell carcinoma; 4, non-small cell lung cancer small-cell carcinoma. For stomach cancer – 1, EMT; 2, MSI; 3, MSS; 4, MSS/TP53-; 5, MSS/TP53+. For subtypes within each cancer, one-way ANOVA analysis was performed to verify the statistical significance, and their mean differences corrected via Bonferroni post-hoc test at p ≤ 0.05. In all graphs to the right, the red markers correspond to subtypes with statistically significant difference in mean value, whereas the black markers indicate statistical non-significance for each corresponding cancer subtypes assigned as box-whisker plot.

**Fig.S7:**
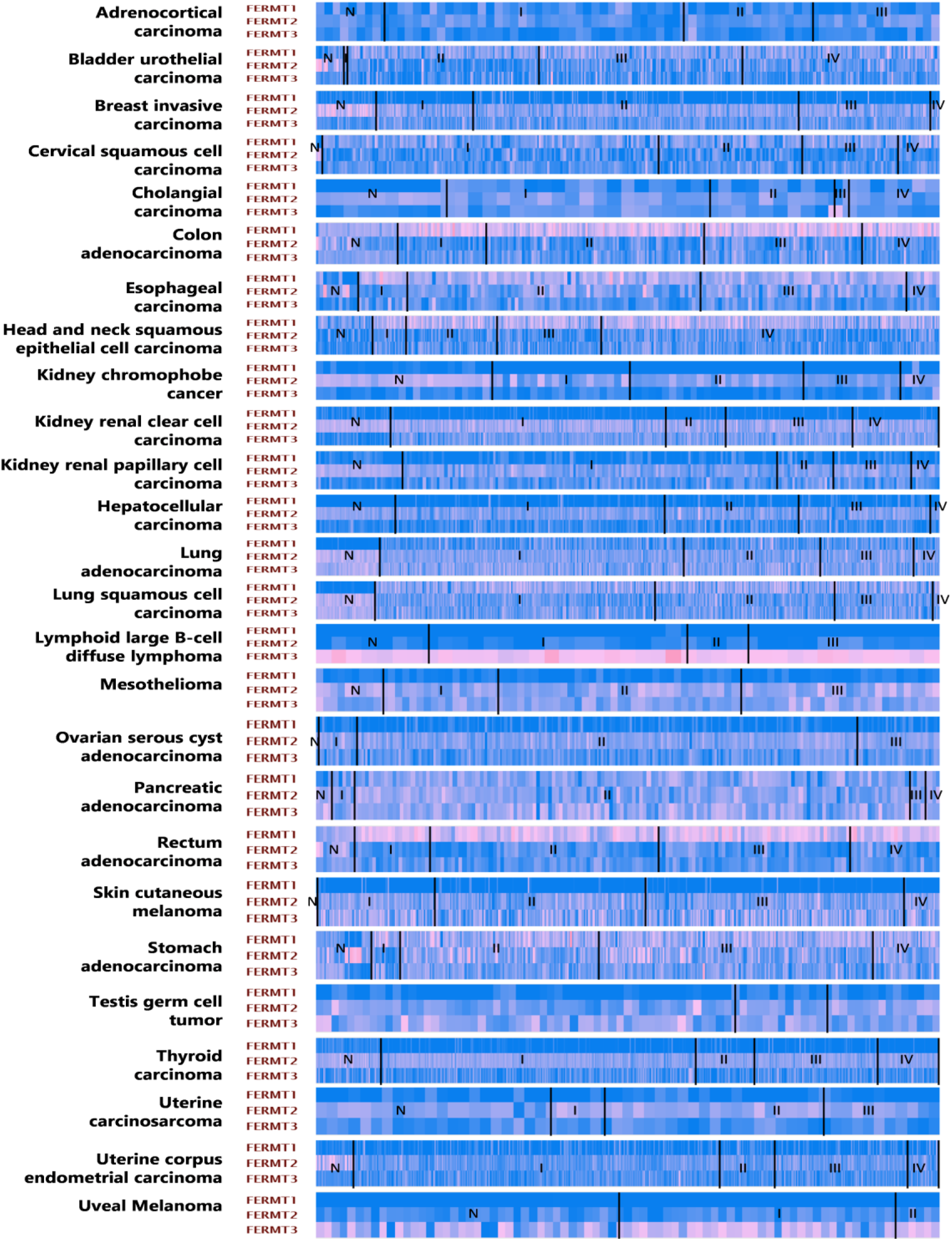
Gene Expression Patterns of all Kindlin Family Proteins across Multiple Tumor Subtypes. Each box across the horizontal heatmap depicts individual samples, and the vertical index enlists multiple cancer types and subtypes for all three kindlin proteins. Red represents the higher levels of expression, whereas blue depicts lower expression. The heatmap is further divided into normal tissue and tumor stages to depict stage-specific expression. N, normal sample; I, tumor stage 1; II, tumor stage 2; III, tumor stage 3; IV, tumor stage 4. All of these stages contain data for corresponding sub-stages and are not divided further for ease of understanding.

**Fig.S8:**
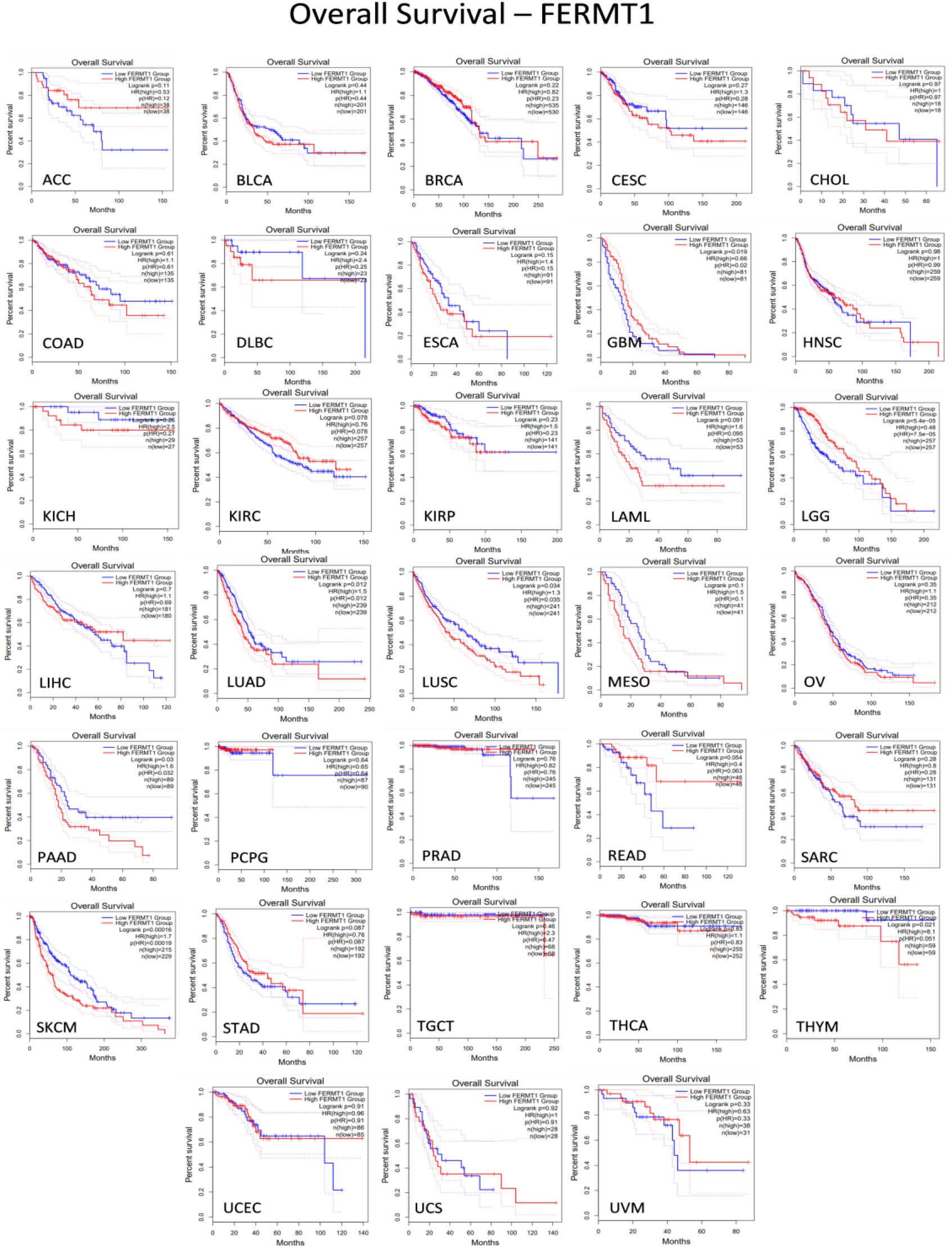
FERMT1 Expression-Specific Kaplan-Meier Plots Depicting Overall Survival (OS) across 33 Cancer Types. The x-axis depicts the number of months that have passed since diagnosis while the y-axis illustrates the survival probability. Each plot has a red curve depicting high FERMT1 transcripts per million, and a blue trend showing low FERMT1 expression levels. This survival analysis data used a median cut-off and 95% confidence intervals, which can be followed by the dotted lines on the graphs. Statistical data is given on the top right corner of each plot, containing log-rank p value, hazard ratio, p-value of hazard ratio obtained by Cox regression test, and the number of patients in the high and low expressing cohorts. Corresponding cancer (sub)-types are written in corresponding images.

**Fig.S9:**
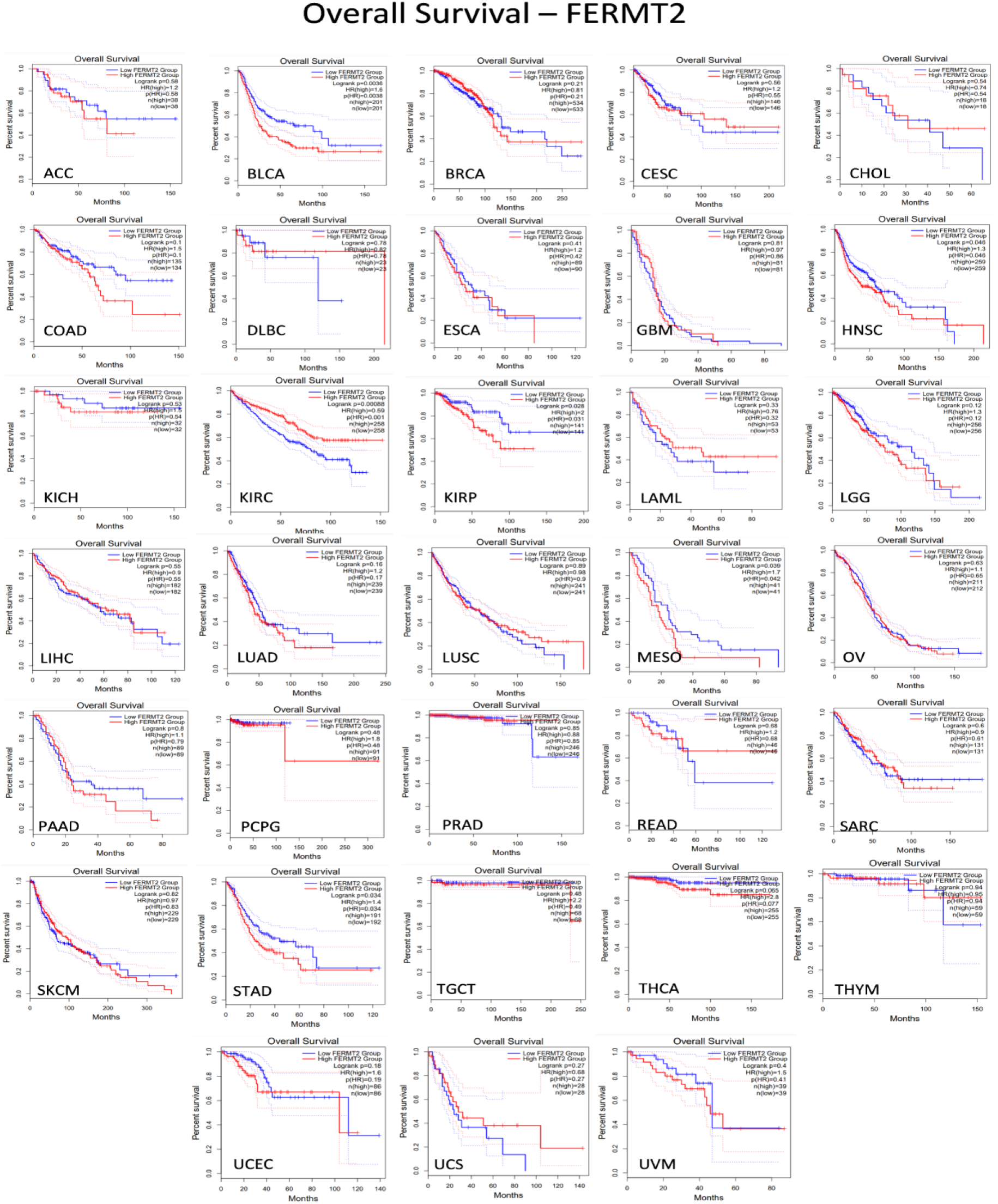
FERMT2 Expression-Specific Kaplan-Meier Plots Depicting Overall Survival (OS) across 33 Cancer Types. The x-axis depicts the number of months that have passed since diagnosis while the y-axis illustrates the survival probability. Each plot has a red curve depicting high FERMT2 transcripts per million, and a blue trend showing low FERMT2 expression levels. This survival analysis data used a median cut-off and 95% confidence intervals, which can be followed by the dotted lines on the graphs. Statistical data is given on the top right corner of each plot, containing log-rank p value, hazard ratio, p-value of hazard ratio obtained by Cox regression test, and the number of patients in the high and low expressing cohorts. Corresponding cancer (sub)-types are written in corresponding images.

**Fig.S10:**
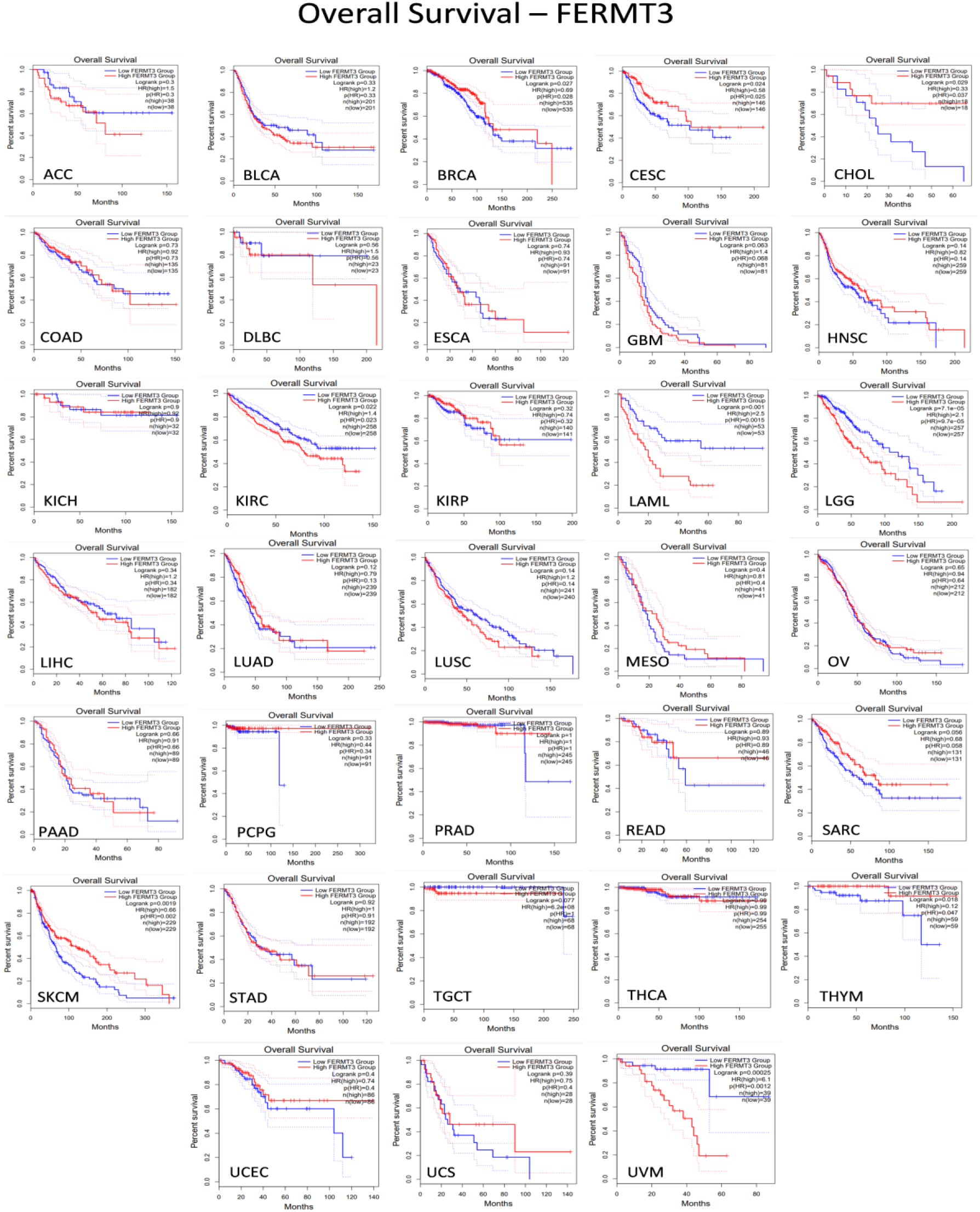
FERMT3 Expression-Specific Kaplan-Meier Plots Depicting Overall Survival (OS) across 33 Cancer Types. The x-axis depicts the number of months that have passed since diagnosis while the y-axis illustrates the survival probability. Each plot has a red curve depicting high FERMT3 transcripts per million, and a blue trend showing low FERMT3 expression levels. This survival analysis data used a median cut-off and 95% confidence intervals, which can be followed by the dotted lines on the graphs. Statistical data is given on the top right corner of each plot, containing log-rank p value, hazard ratio, p-value of hazard ratio obtained by Cox regression test, and the number of patients in the high and low expressing cohorts. Corresponding cancer (sub)-types are written in corresponding images.

**Fig.S11:**
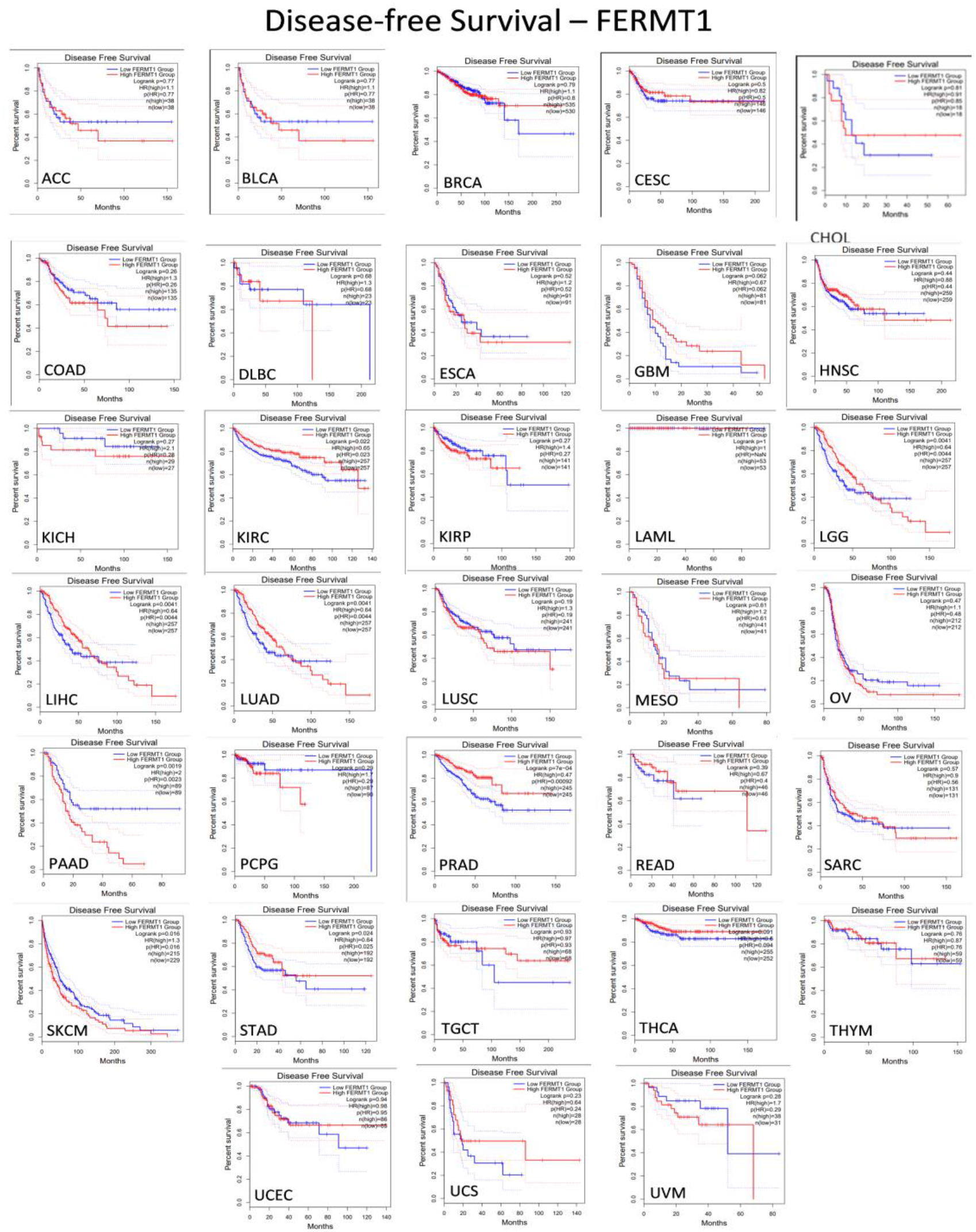
FERMT1 Expression-Specific Kaplan-Meier Plots Depicting Disease-Free Survival (DFS) across 33 Cancer Types. The number of months passed after treatment can be found on the x-axis, while the survival probability can be found on the y-axis. Each plot has a red curve depicting high FERMT1 transcripts per million, and a blue trend showing low FERMT1 expression levels. This survival analysis data used a median cut-off and 95% confidence intervals, which can be followed by the dotted lines on the graphs. Statistical data is given on the top right corner of each plot, containing log-rank p value, hazard ratio, p-value of hazard ratio obtained by Cox regression test, and the number of patients in the high and low expressing cohorts. Corresponding cancer (sub)-types are written in corresponding images.

**Fig.S12:**
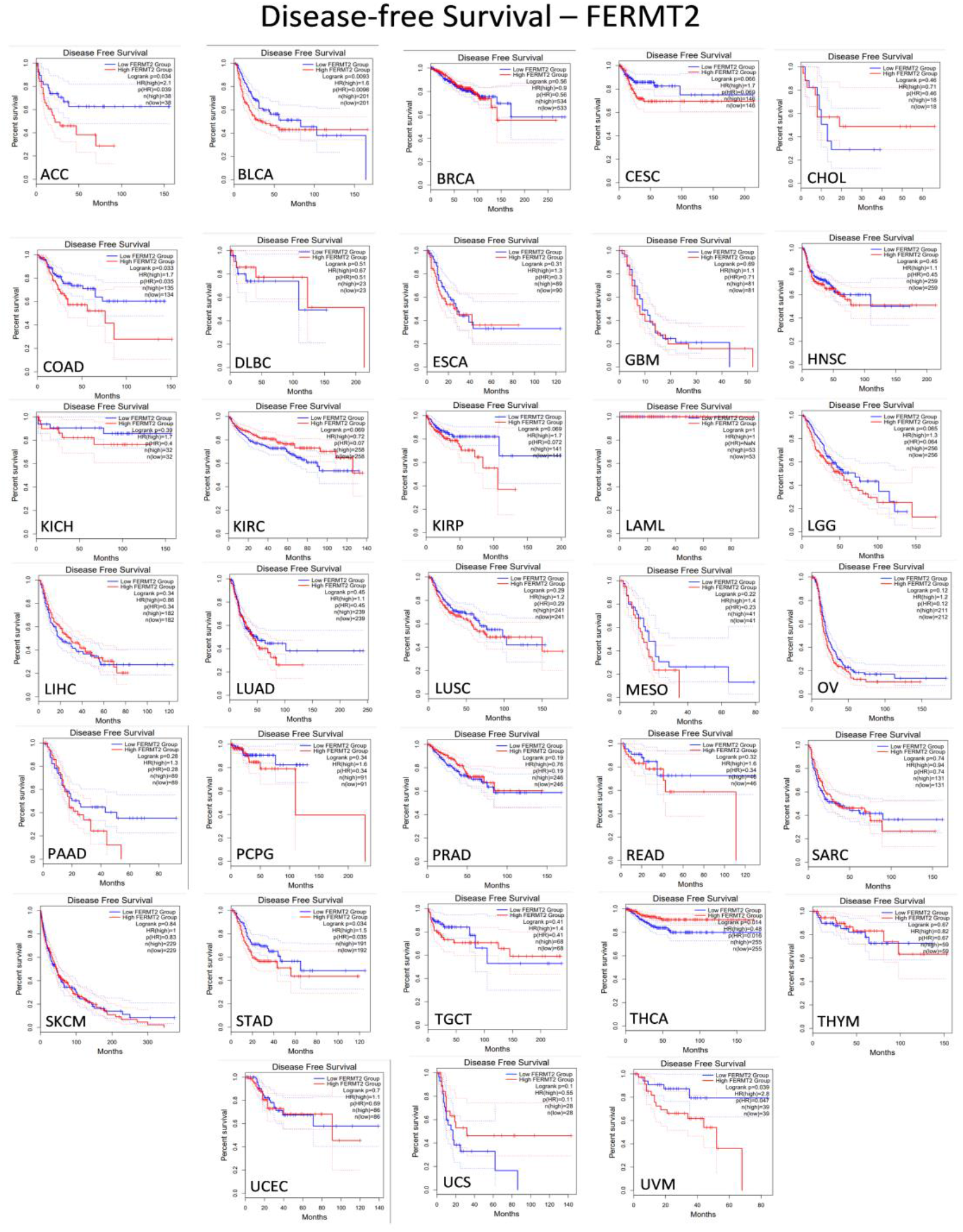
FERMT2 Expression-Specific Kaplan-Meier Plots Depicting Disease-Free Survival (DFS) across 33 Cancer Types. The number of months passed after treatment can be found on the x-axis, while the survival probability can be found on the y-axis. Each plot has a red curve depicting high FERMT2 transcripts per million, and a blue trend showing low FERMT2 expression levels. This survival analysis data used a median cut-off and 95% confidence intervals, which can be followed by the dotted lines on the graphs. Statistical data is given on the top right corner of each plot, containing log-rank p value, hazard ratio, p-value of hazard ratio obtained by Cox regression test, and the number of patients in the high and low expressing cohorts. Corresponding cancer (sub)-types are written in corresponding images.

**Fig.S13:**
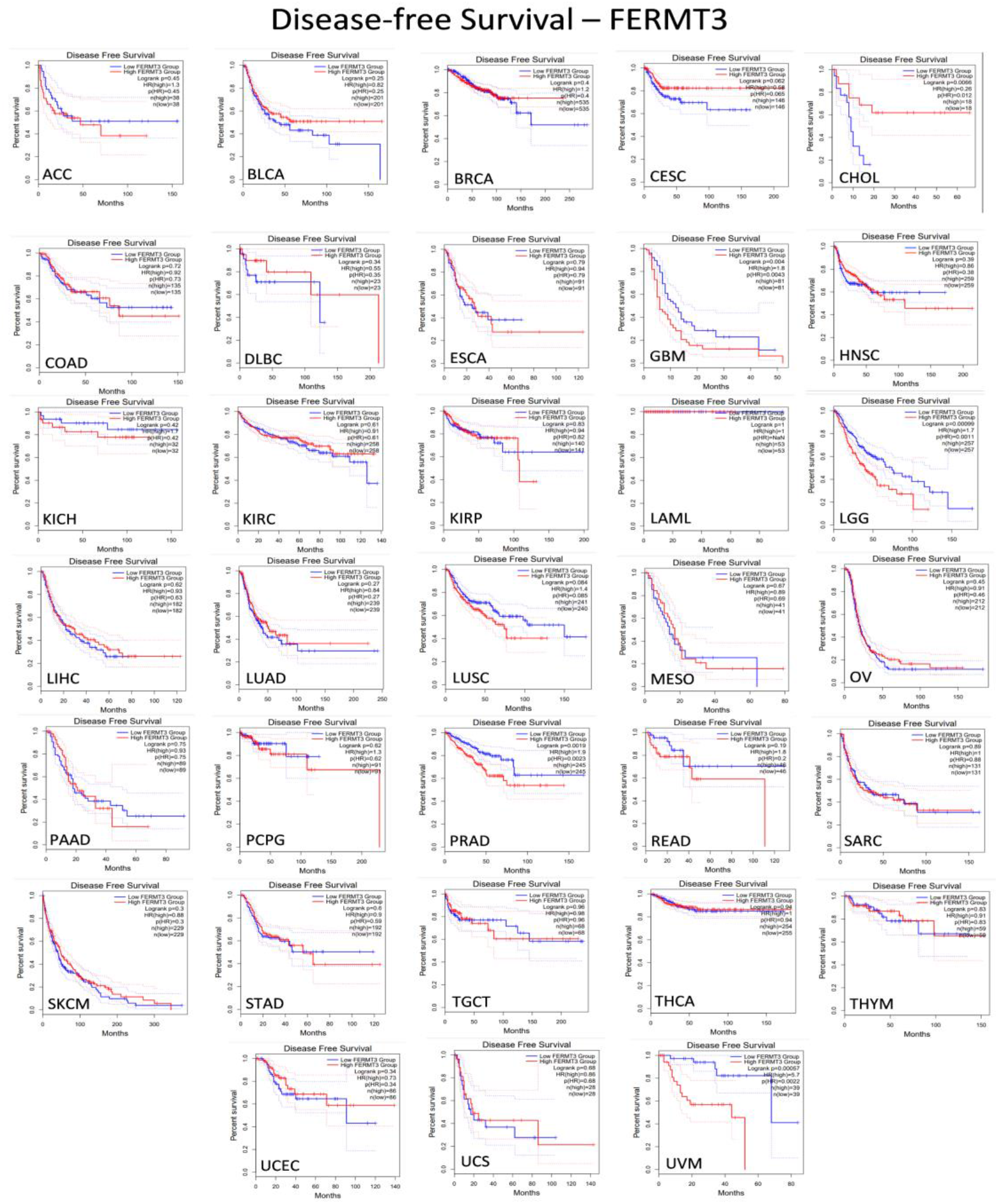
FERMT3 Expression-Specific Kaplan-Meier Plots Depicting Disease-Free Survival (DFS) across 33 Cancer Types. The number of months passed after treatment can be found on the x-axis, while the survival probability can be found on the y-axis. Each plot has a red curve depicting high FERMT3 transcripts per million, and a blue trend showing low FERMT3 expression levels. This survival analysis data used a median cut-off and 95% confidence intervals, which can be followed by the dotted lines on the graphs. Statistical data is given on the top right corner of each plot, containing log-rank p value, hazard ratio, p-value of hazard ratio obtained by Cox regression test, and the number of patients in the high and low expressing cohorts. Corresponding cancer (sub)-types are written in corresponding images.

**Fig.S14:**
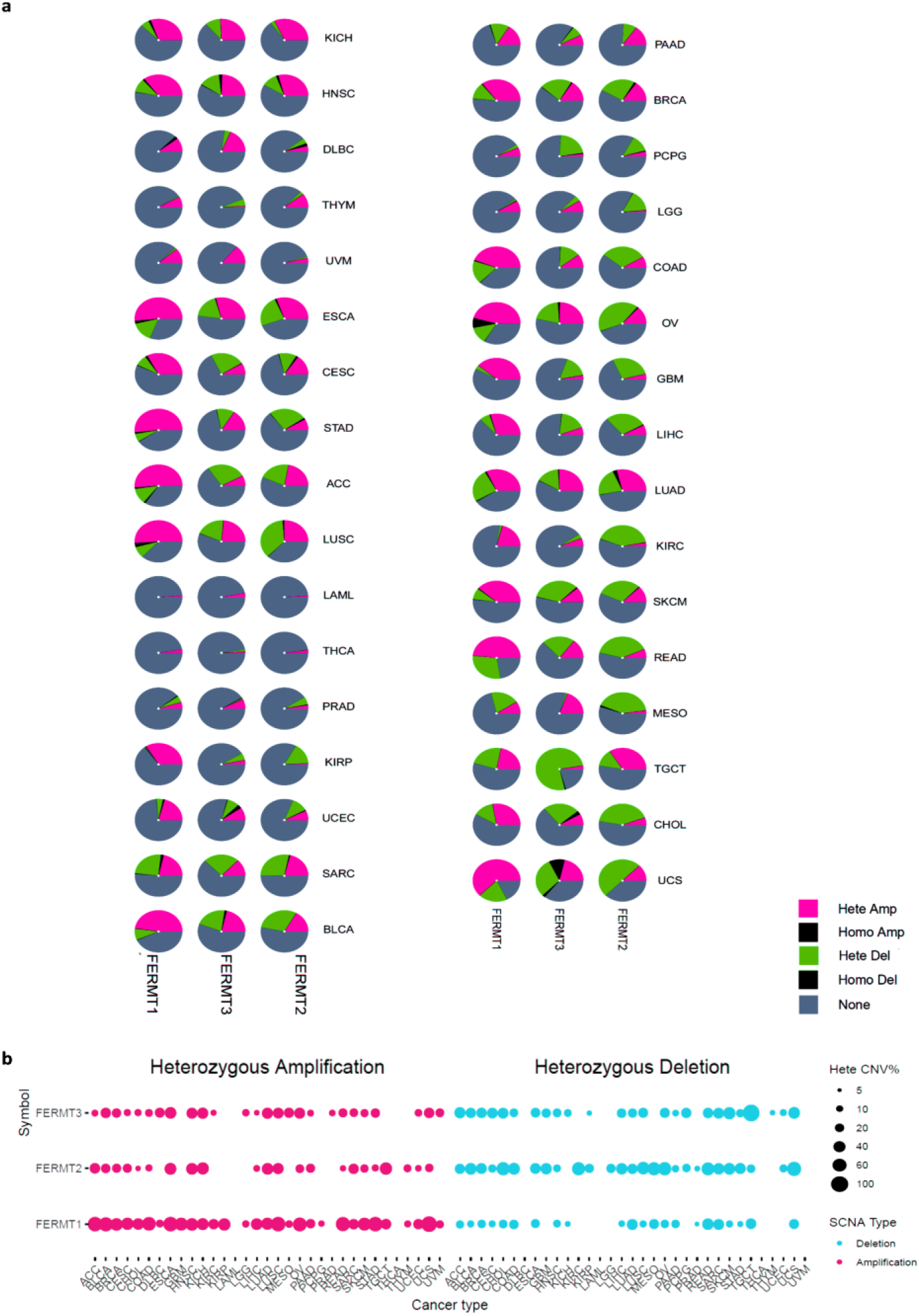
Copy Number Variation (CNV) Analysis in 33 Cancer Types. a. CNV pie plot represents global CNV profiling, showing the make-up of Heterozygous/Homozygous CNV of each kindlin gene in various cancer types. A pie represents the proportion of different types of CNV of one gene in one cancer, and different color represent different types of CNV. The proportion of each color represents the composition of that particular CNV in that cancer. b. Heterozygous CNV profile depicts the percentage of heterozygous CNV, including amplification and deletion of each gene in each cancer. A cut-off of >5% CNV in samples were considered. Hete Amp, heterozygous amplification; Hete Del, heterozygous deletion; Homo Amp, homozygous amplification; Homo Del, homozygous deletion; None, no CNV. The size of the points represents statistical significance so that bigger size implies more significance.

**Fig.S15:**
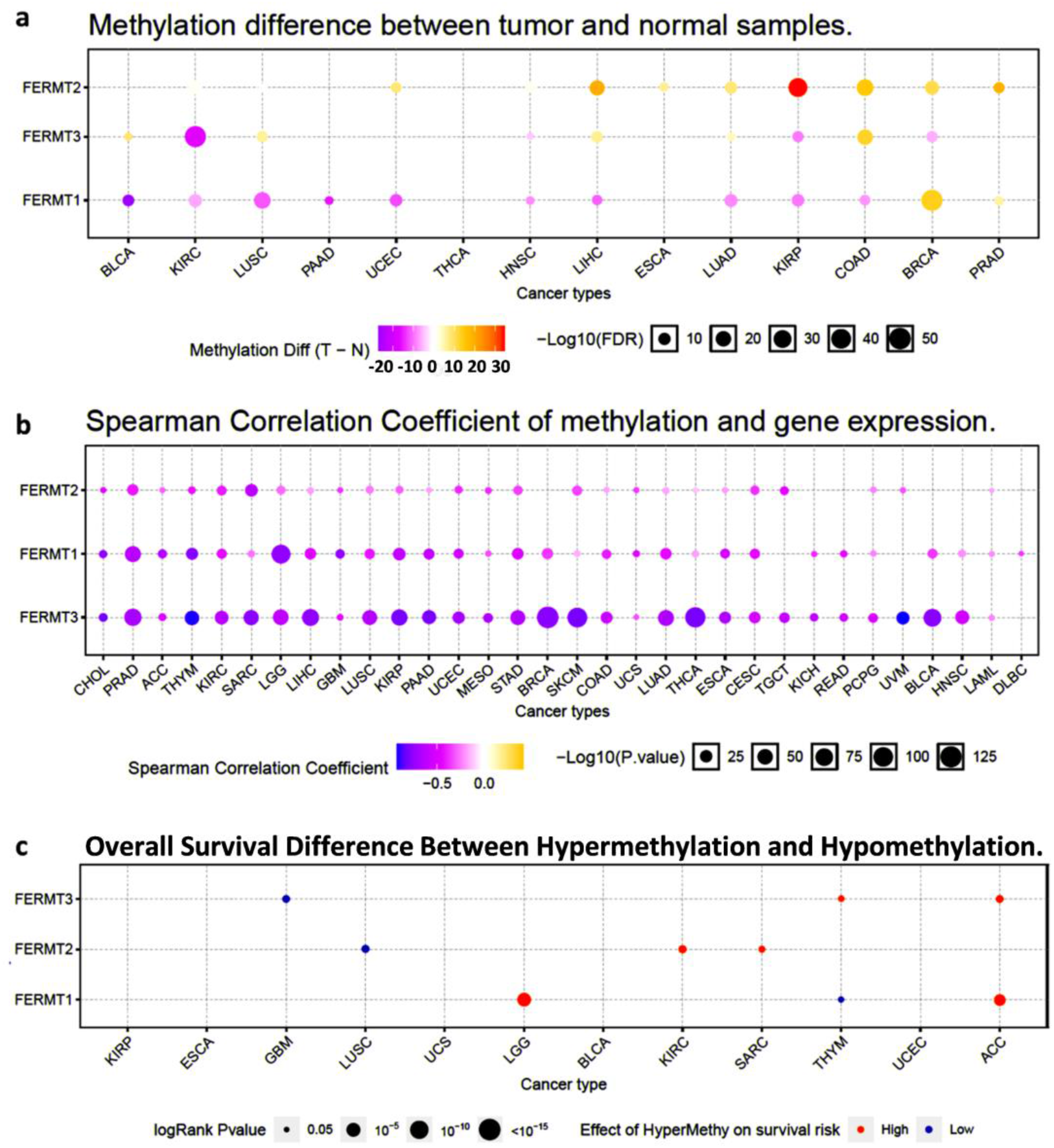
Gene Methylation Pattern Analysis for All Kindlin Genes across 14 Cancer Types. a. Differential gene methylation pattern between tumor and normal samples in different cancer types. Purple points indicate hypomethylation, whereas orange points indicate hypermethylation, both in case of tumor. The size of the bubbles is represented as –Log10 (FDR) scale, representing more statistical significance with increasing bubble size. The darkness of colors is representative of higher value of (tumor – normal). b. Genetic correlation between gene methylation and mRNA expression for FERMT genes derived as Spearman’s correlation coefficient. More blue coloration indicates a negative correlation whereas redder represents positive correlation. The darkness of colors is representative of higher magnitude of correlation. The size of the bubbles is represented as –Log10 (FDR) scale, representing more statistical significance with increasing bubble size. c. Survival difference between samples with FERMT genes’ hyper- and hypo-methylation. Statistical significance of survivability difference has been represented using log-rank p-value (≤ 0.05). Red points represent hyper-methylation with worse survivability, whereas blue points indicate hypo-methylation with better survivability. Bigger size of bubble is indicative of better significance.

**Fig.S16:**
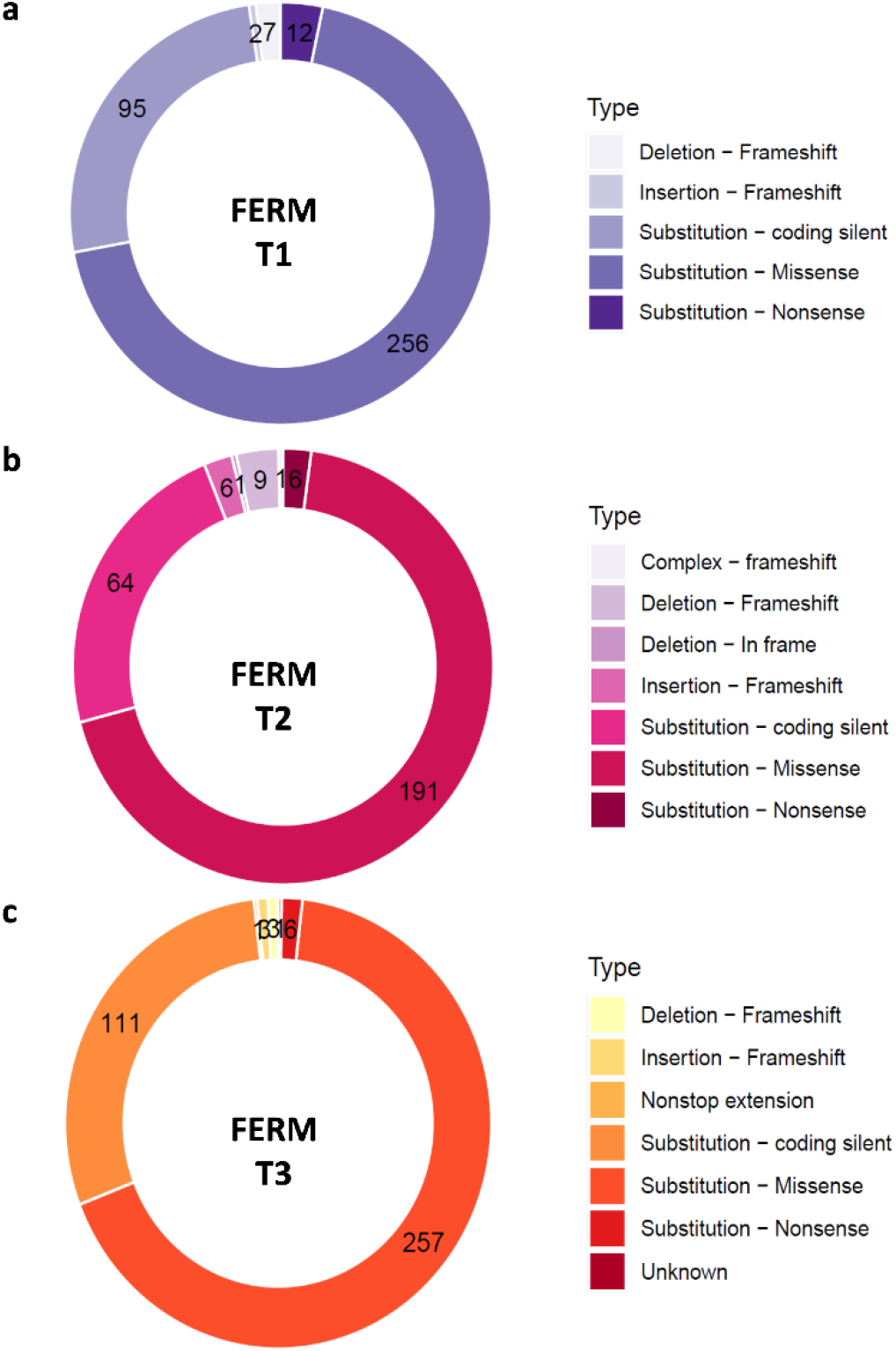
Distribution of Point Mutation Subtypes across All the Kindlin Genes across 33 Different Types of Cancer. The size of doughnut fragments indicates percentage of mutation type contributing to cancer for FERMT1 (a), FERMT2 (b), and FERMT3 (c). The numbers on each fragment show the actual number of mutations. The color codes for each mutation type are depicted adjacent to the corresponding image.

**Fig.S17:**
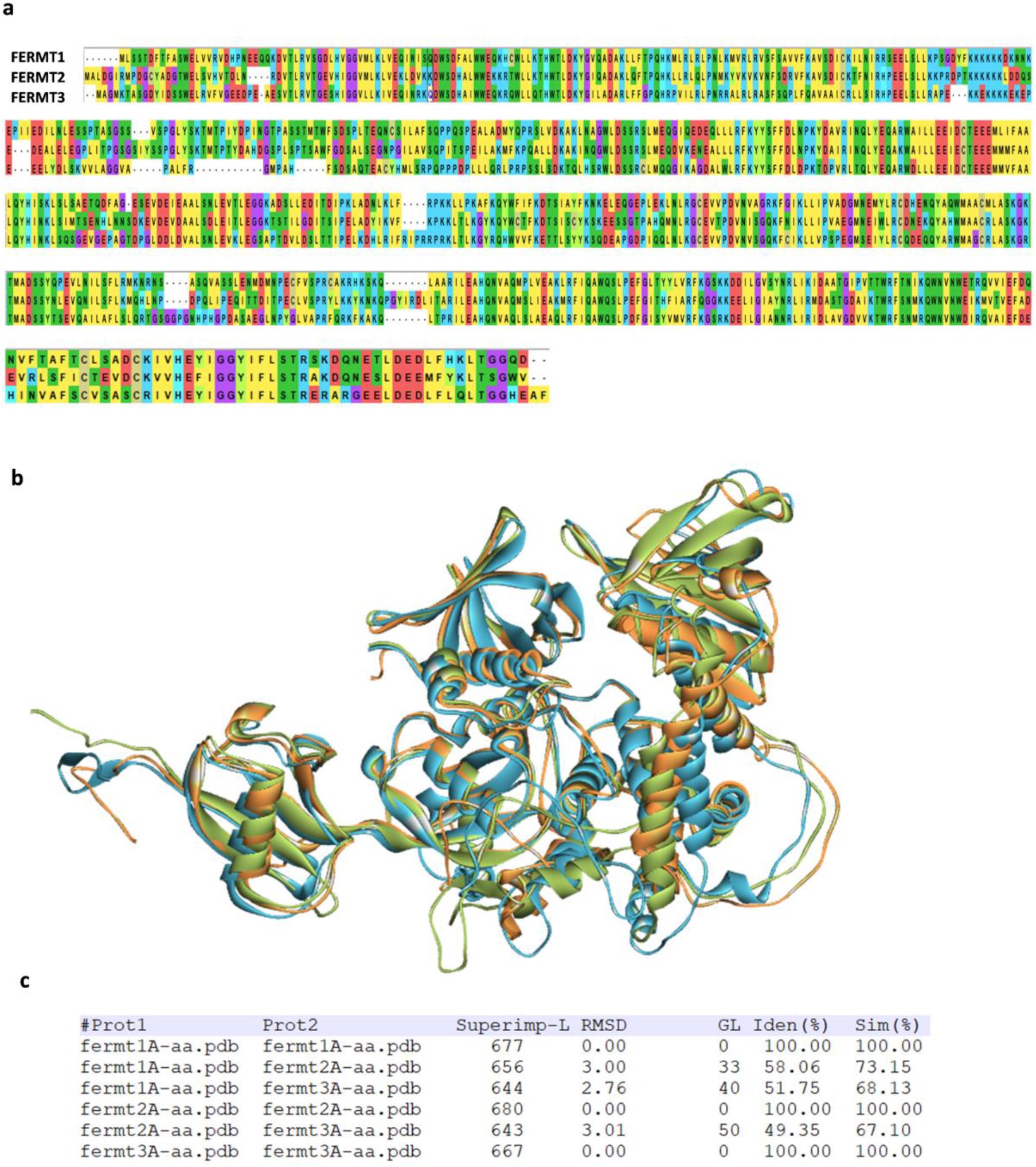
Sequence Alignment of All Kindlin Family Proteins Along with Structural Superimposition. a. Multiple sequence alignment of FERMT1, FERMT2, and FERMT3 amino acid sequences. The dots represent gaps with respect to conservativeness. Different colors indicate types of amino acids according to their similarity. Red, acidic; sky blue, basic; yellow, hydrophobic; light green, aromatic; deep green, polar; deep yellow, sulphur-containing; purple, glycine. b. Structural superimposition of kindlin1, kindlin2, and kindlin3 full-length protein monomers. Green, kindlin1; sky blue, kindlin2; orange, kindlin3. c. Protein sequence similarity and identity between different kindlins. Iden (%), percentage identity; Sim (%), percentage similarity; RMSD, root mean-squared deviation; GL, gap length; Superimp-L, superimposed sequence length.

**Fig.S18:**
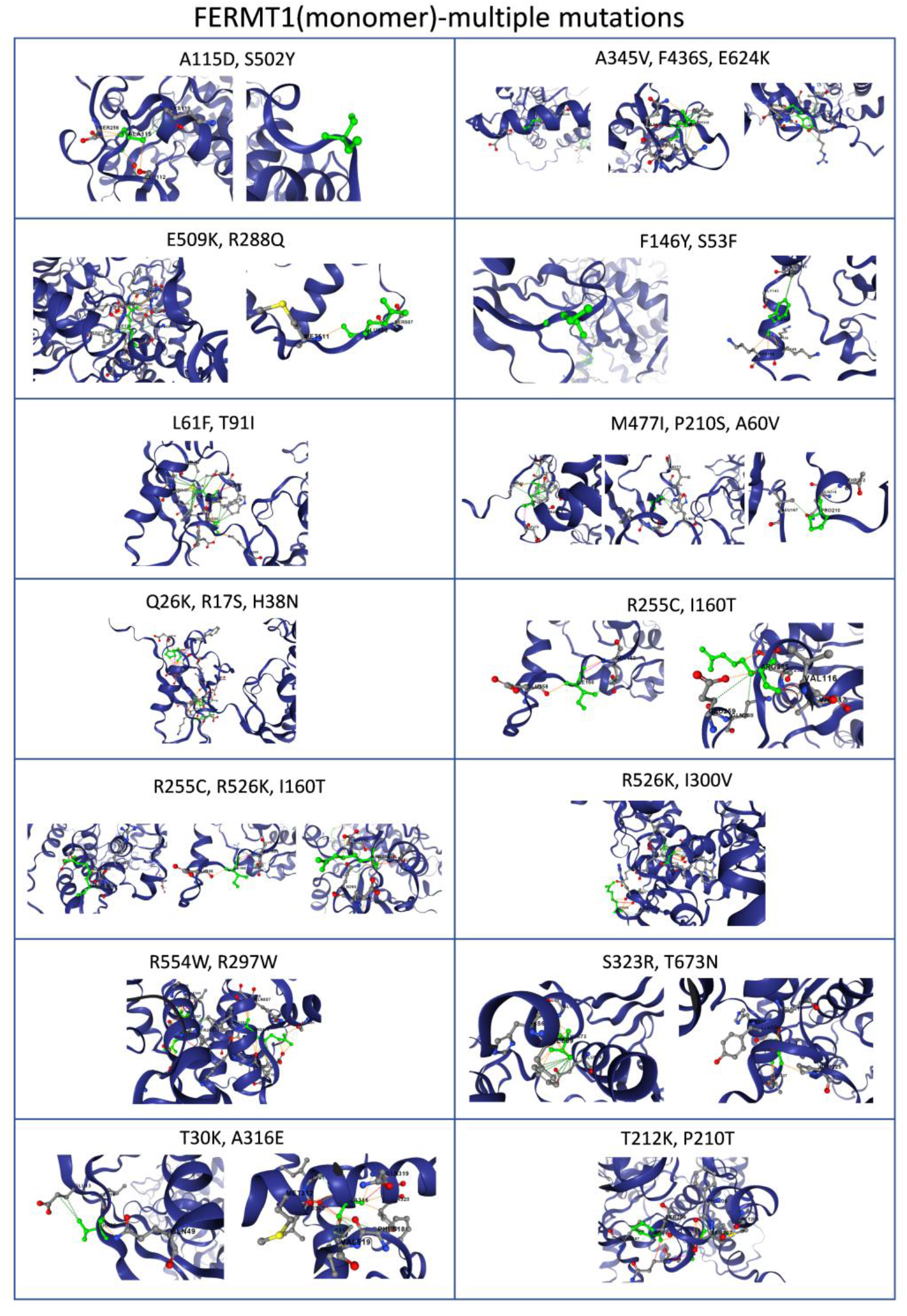
Visualisation of Non-Covalent Interactions Caused by the Mutated Residues on FERMT1 Monomer Structures with Multiple Mutations. Cartoon representation is utilised, and interactions are coloured according to different types: dark green for hydrophobicity, red for hydrogen bonding, dark blue for carbonyl interaction, yellow for ionic interactions, orange for polarity, light blue for Van der Waals interactions, light green for aromatic interaction, and pink for clashes. The mutated amino acid residue and the residue with which its interaction has modified because of mutations are indicated using three-letter amino acid code with insertion ID (residue number). The mutated residue is shown in green, whereas interacting atoms are shown in red. The protein chain is shown in dark blue.

**Fig.S19:**
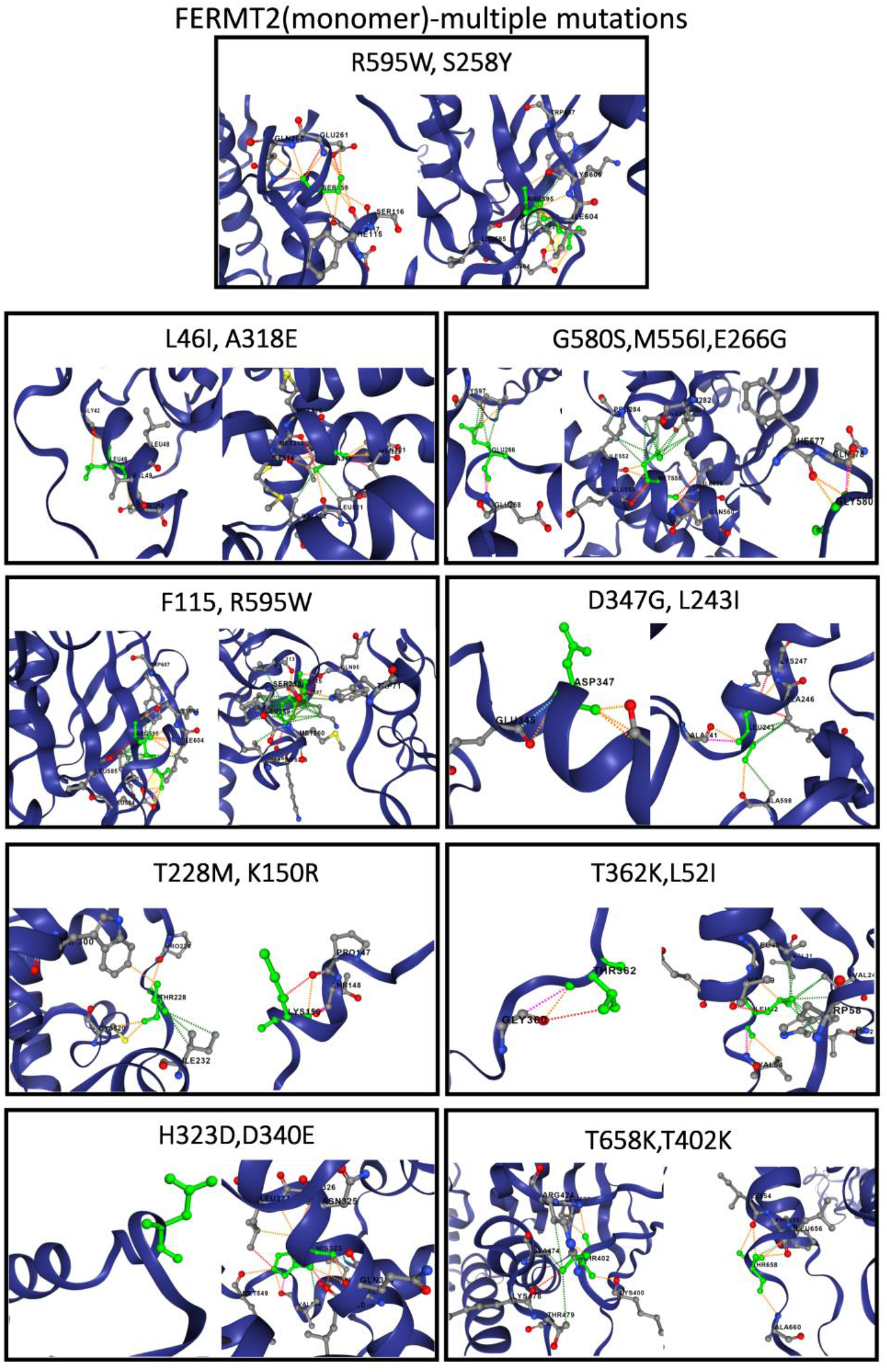
Visualisation of Non-Covalent Interactions Caused by the Mutated Residues on FERMT2 Monomer Structures with Multiple Mutations. Cartoon representation is utilised, and interactions are coloured according to different types: dark green for hydrophobicity, red for hydrogen bonding, dark blue for carbonyl interaction, yellow for ionic interactions, orange for polarity, light blue for Van der Waals interactions, light green for aromatic interaction, and pink for clashes. The mutated amino acid residue and the residue with which its interaction has modified because of mutations are indicated using three-letter amino acid code with insertion ID (residue number). The mutated residue is shown in green, whereas interacting atoms are shown in red. The protein chain is shown in dark blue.

**Fig.S20:**
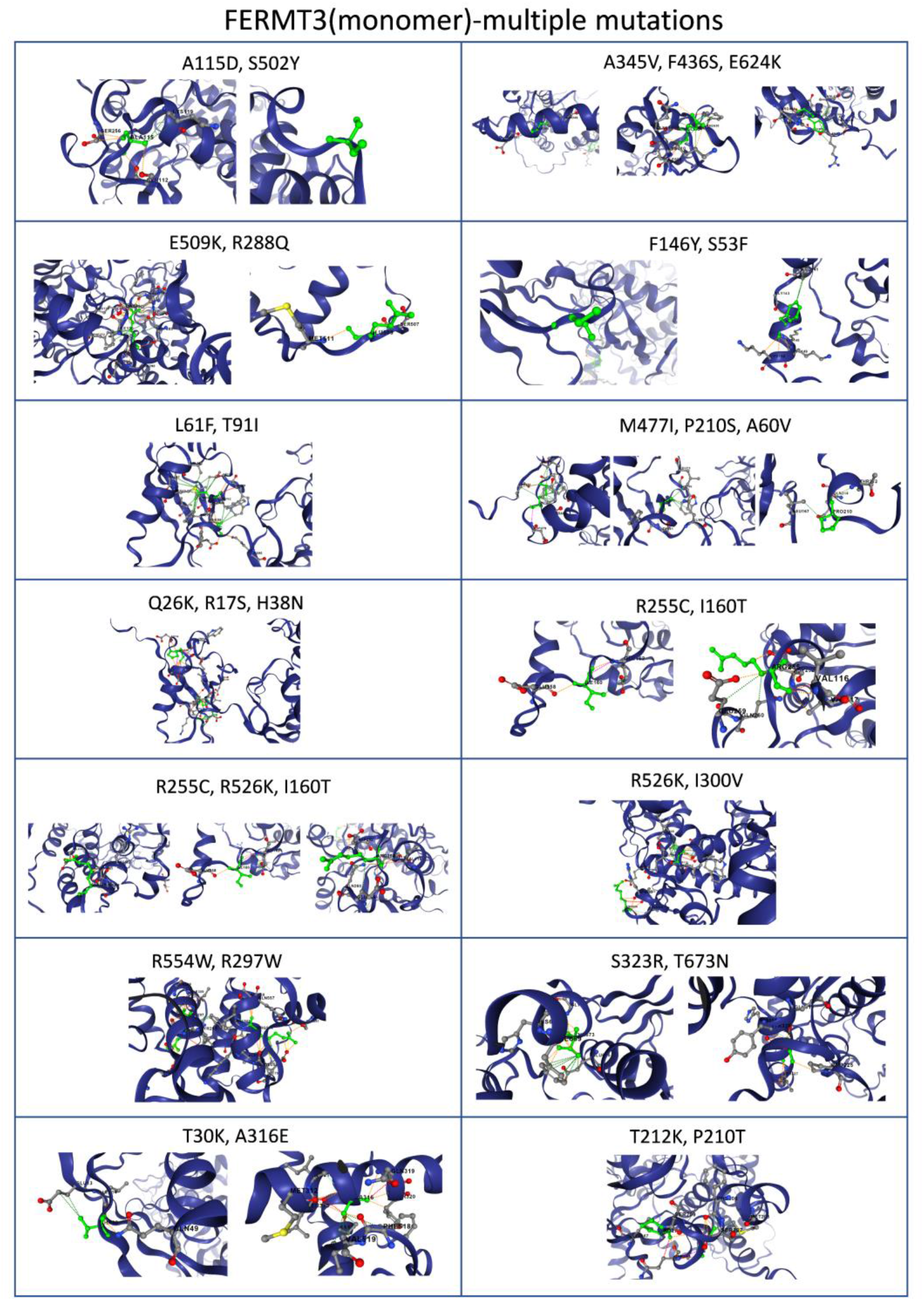
Visualisation of Non-Covalent Interactions Caused by the Mutated Residues on FERMT3 Monomer Structures with Multiple Mutations. Cartoon representation is utilised, and interactions are coloured according to different types: dark green for hydrophobicity, red for hydrogen bonding, dark blue for carbonyl interaction, yellow for ionic interactions, orange for polarity, light blue for Van der Waals interactions, light green for aromatic interaction, and pink for clashes. The mutated amino acid residue and the residue with which its interaction has modified because of mutations are indicated using three-letter amino acid code with insertion ID (residue number). The mutated residue is shown in green, whereas interacting atoms are shown in red. The protein chain is shown in dark blue.

**Fig.S21:**
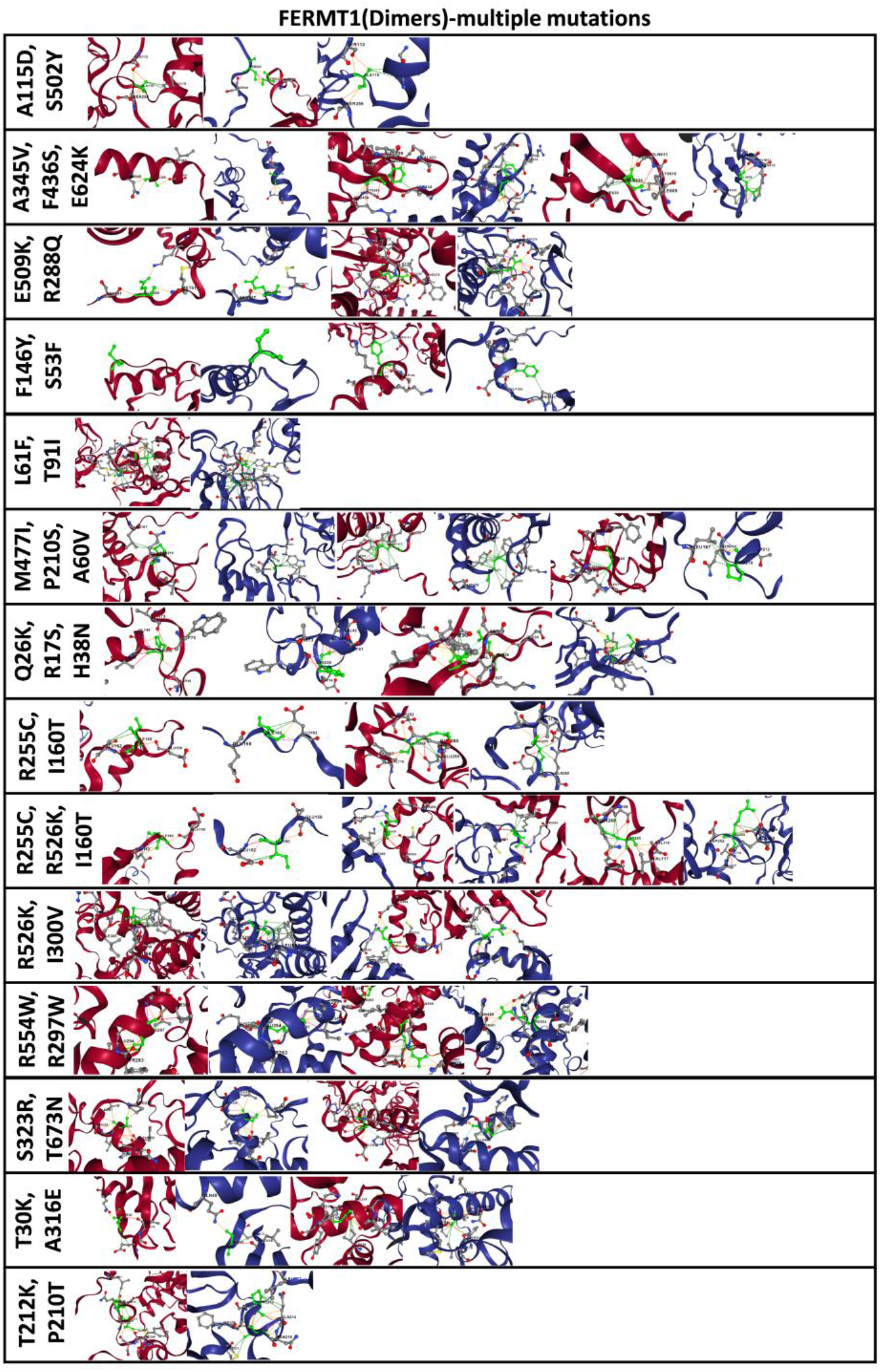
The Impact of Multiple Mutations on the molecular interactions of Dimeric FERMT1. Non-covalent interactions are visualised and can be distinguished by colour: dark green for hydrophobicity; red for hydrogen bonding; dark blue for carbonyl interaction; yellow for ionic interactions; orange for polarity; light blue for Van der Waals interactions; light green for aromatic interaction; and pink for clashes. The mutated amino acid residue and the residue with which its interaction has modified because of mutations are indicated using three-letter amino acid code with insertion ID (residue number). The mutated residue is shown in green, whereas interacting atoms are shown in red. The protein chains are shown in dark blue and dark red, respectively for Chain B and Chain A. If there is a total breakage in interaction, then no bonding or interactions are shown.

**Fig.S22:**
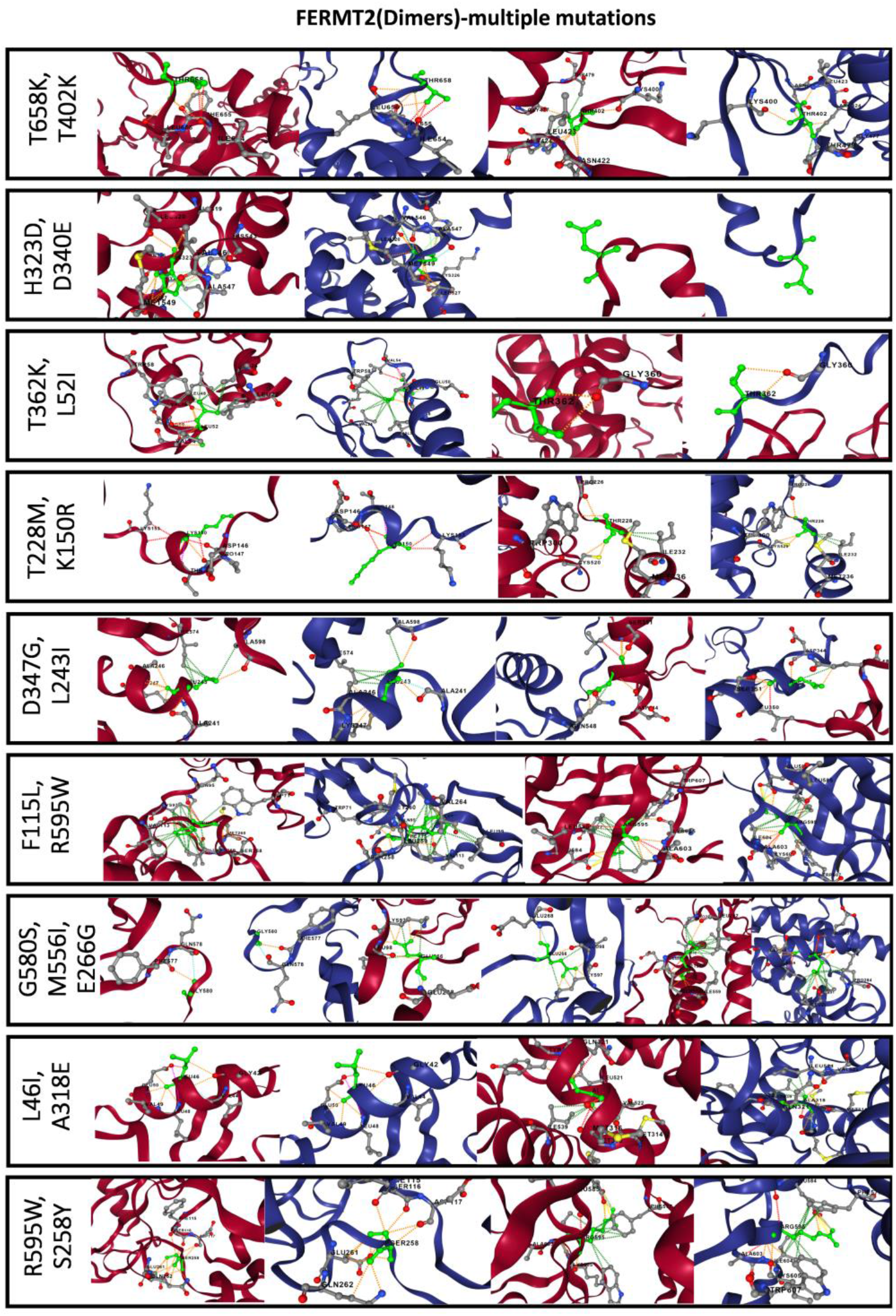
The Impact of Multiple Mutations on the molecular interactions of Dimeric FERMT2. Non-covalent interactions are visualised and can be distinguished by colour: dark

**Fig.S23:**
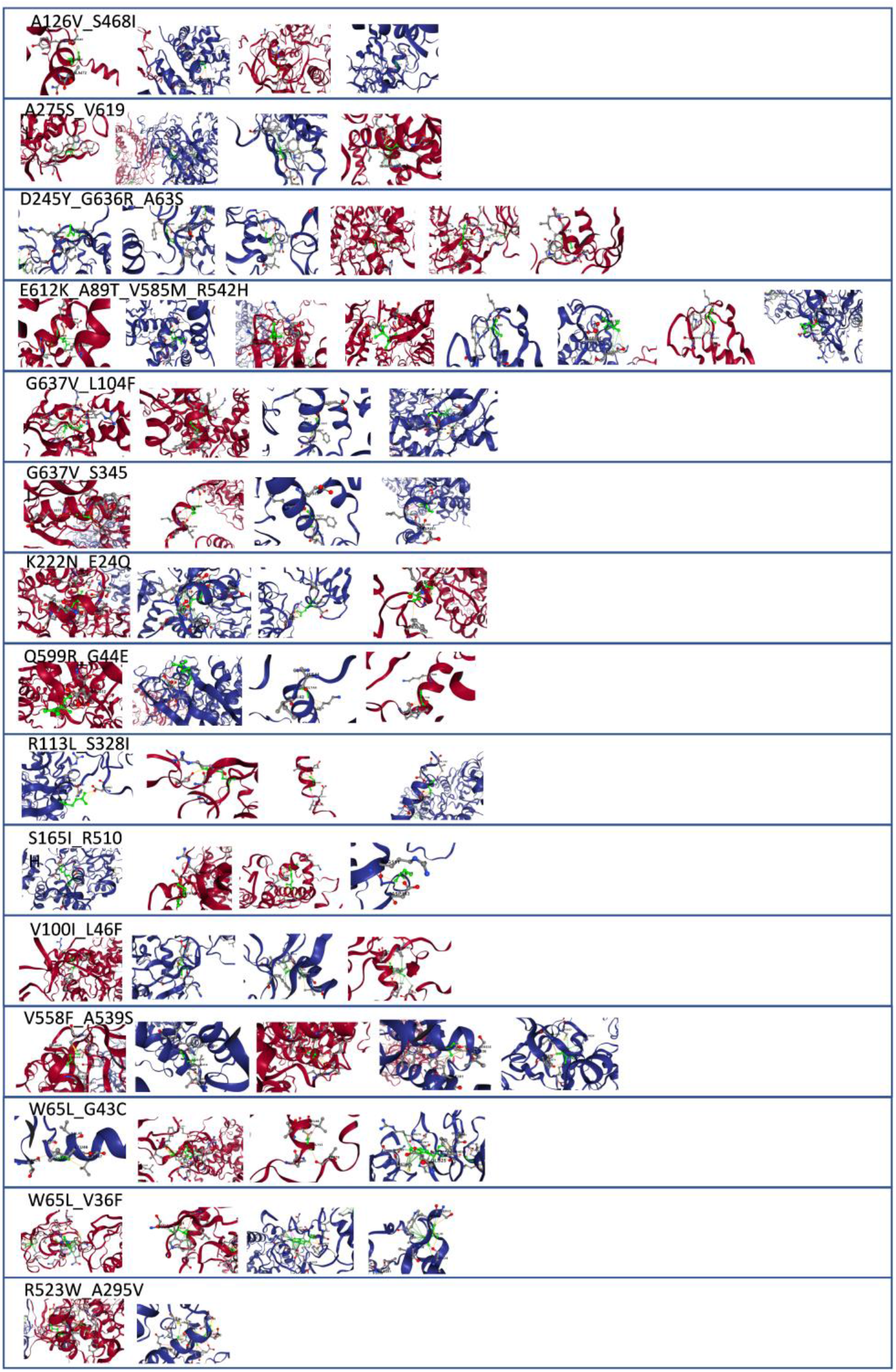
The Impact of Multiple Mutations on the molecular interactions of Dimeric FERMT3. Non-covalent interactions are visualised and can be distinguished by colour: dark

**Fig.S24:**
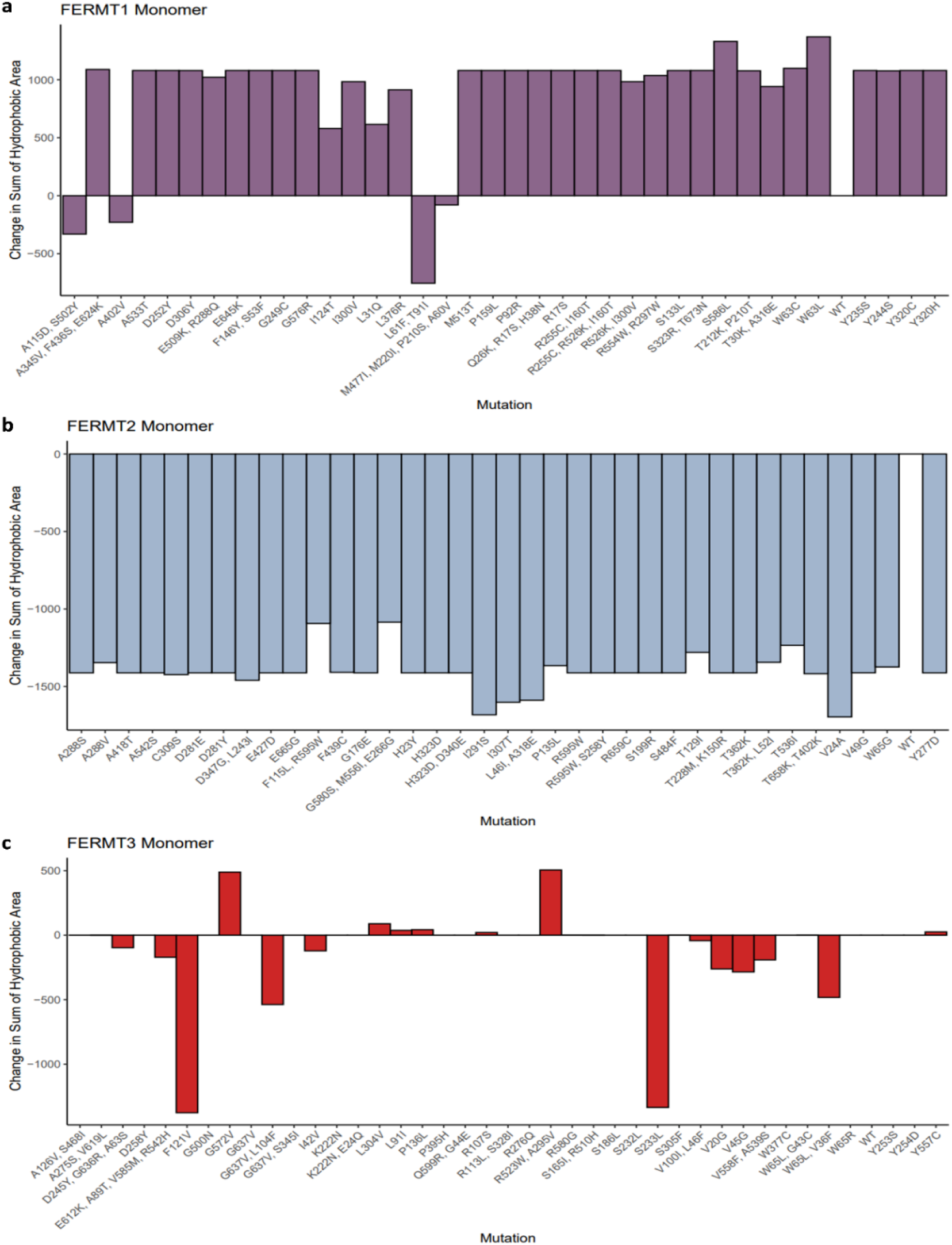
Change in sum of hydrophobic area in the mutated monomeric structures of FERMT1, FERMT2, and FERMT3. The x-axis lists all the mutations, and the y-axis shows the change in the sum of hydrophobic area after being mutated calculated by finding the absolute difference between the sum of hydrophobic area in the wild type vs. the mutated structure.

**Fig.S25:**
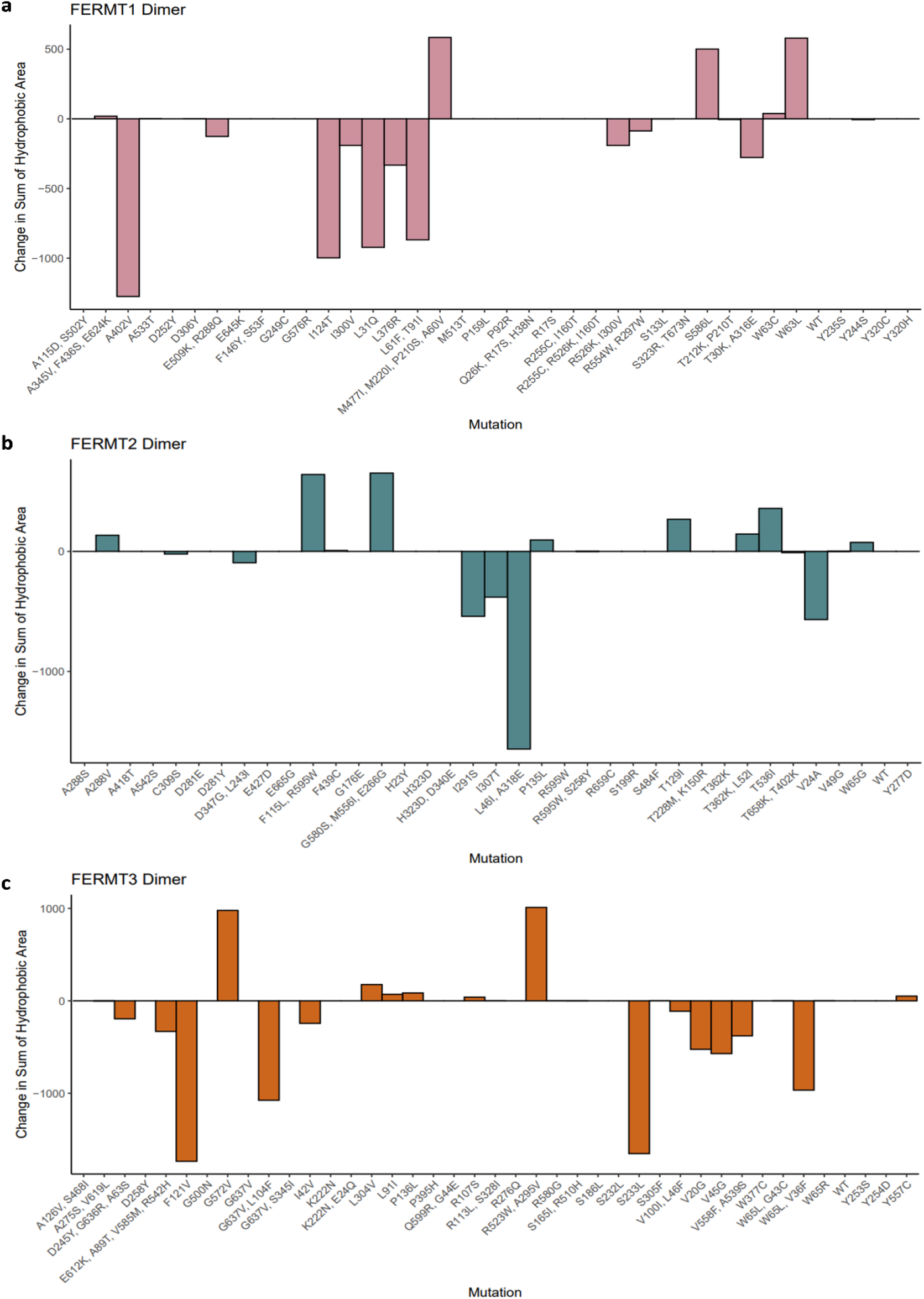
Change in sum of hydrophobic area for each mutated dimeric structure of FERMT1, FERMT2, and FERMT3. All the mutations are listed on the x-axis, while the change in the sum of areas of hydrophobic clusters are represented by the y-axis.

**Fig.S26:**
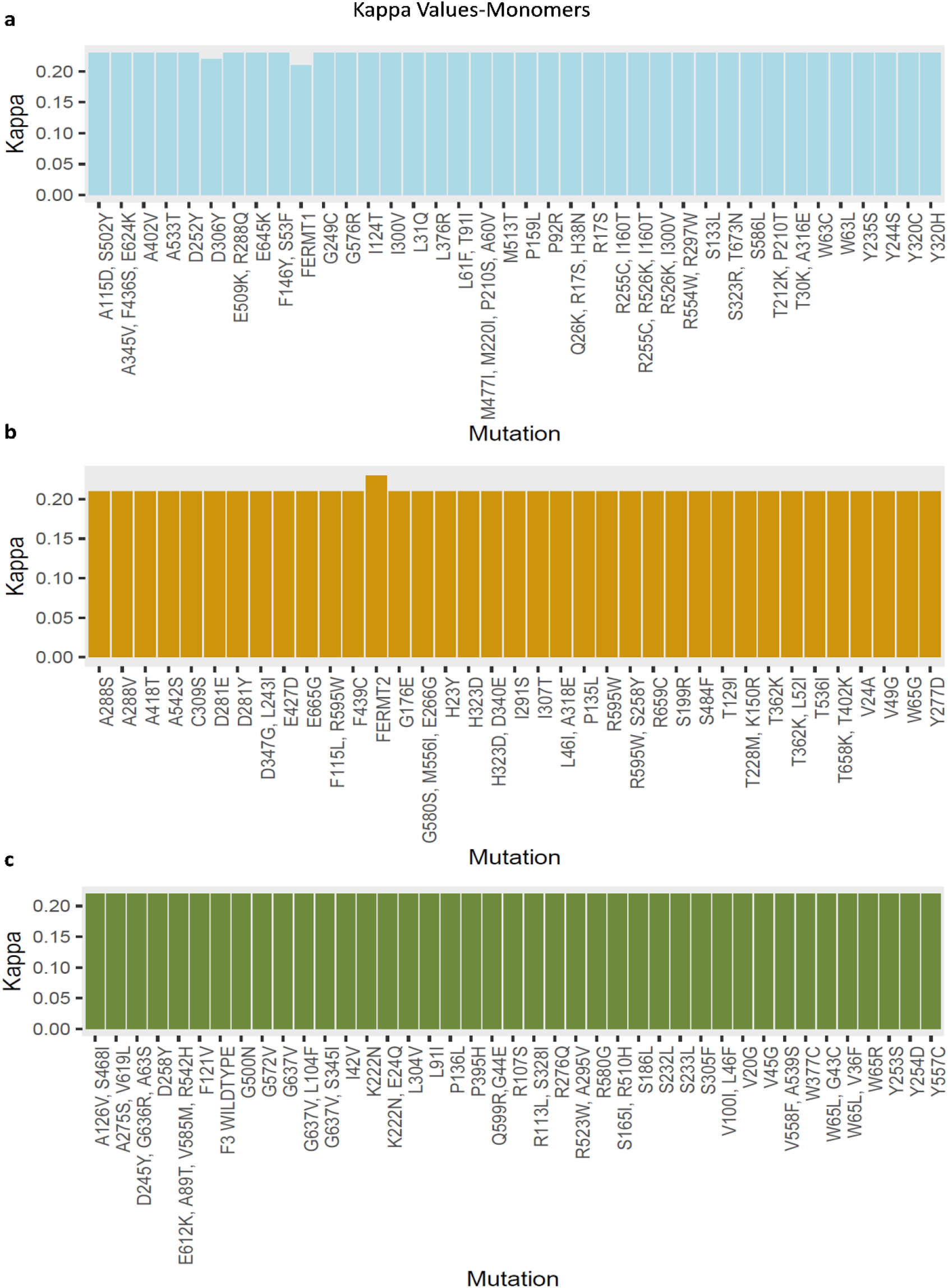
Salt-Bridge Characterization for Each Monomeric Kindlin Family Protein. Kappa (κ) values of each mutated monomeric structure of FERMT1, FERMT2, and FERMT3 represent the salt-bridges present in the structure. The horizontal axis lists the mutations present in the structures, and the vertical axis shows the kappa (κ) values.

**Fig.S27:**
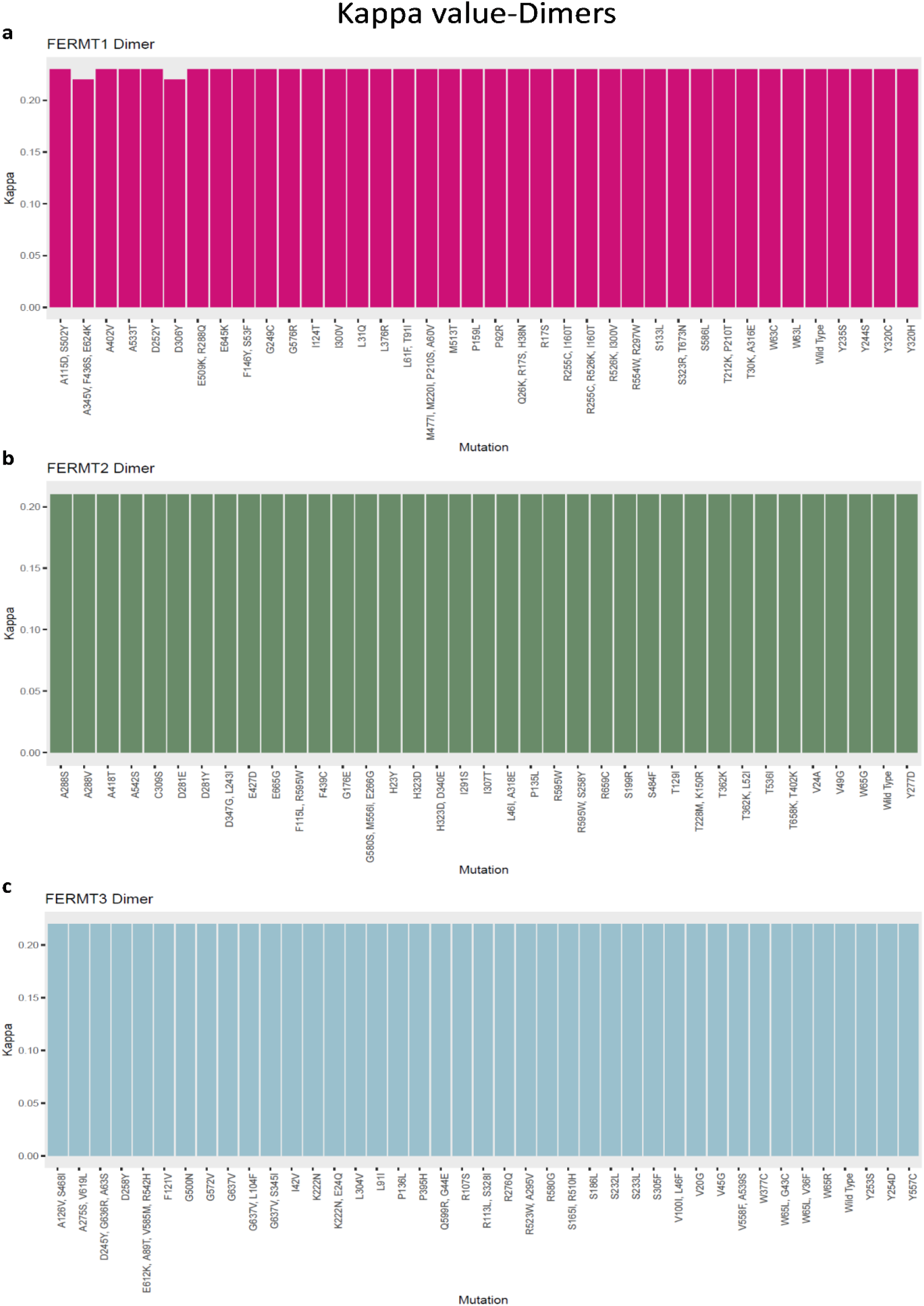
Salt-Bridge Characterization for Each Dimeric Kindlin Family Protein. Kappa (κ) values of each mutated dimeric structure of FERMT1, FERMT2, and FERMT3 represent the salt-bridges present in the structure. The horizontal axis lists the mutations present in the structures, and the vertical axis shows the kappa values.

**Fig.S28.**
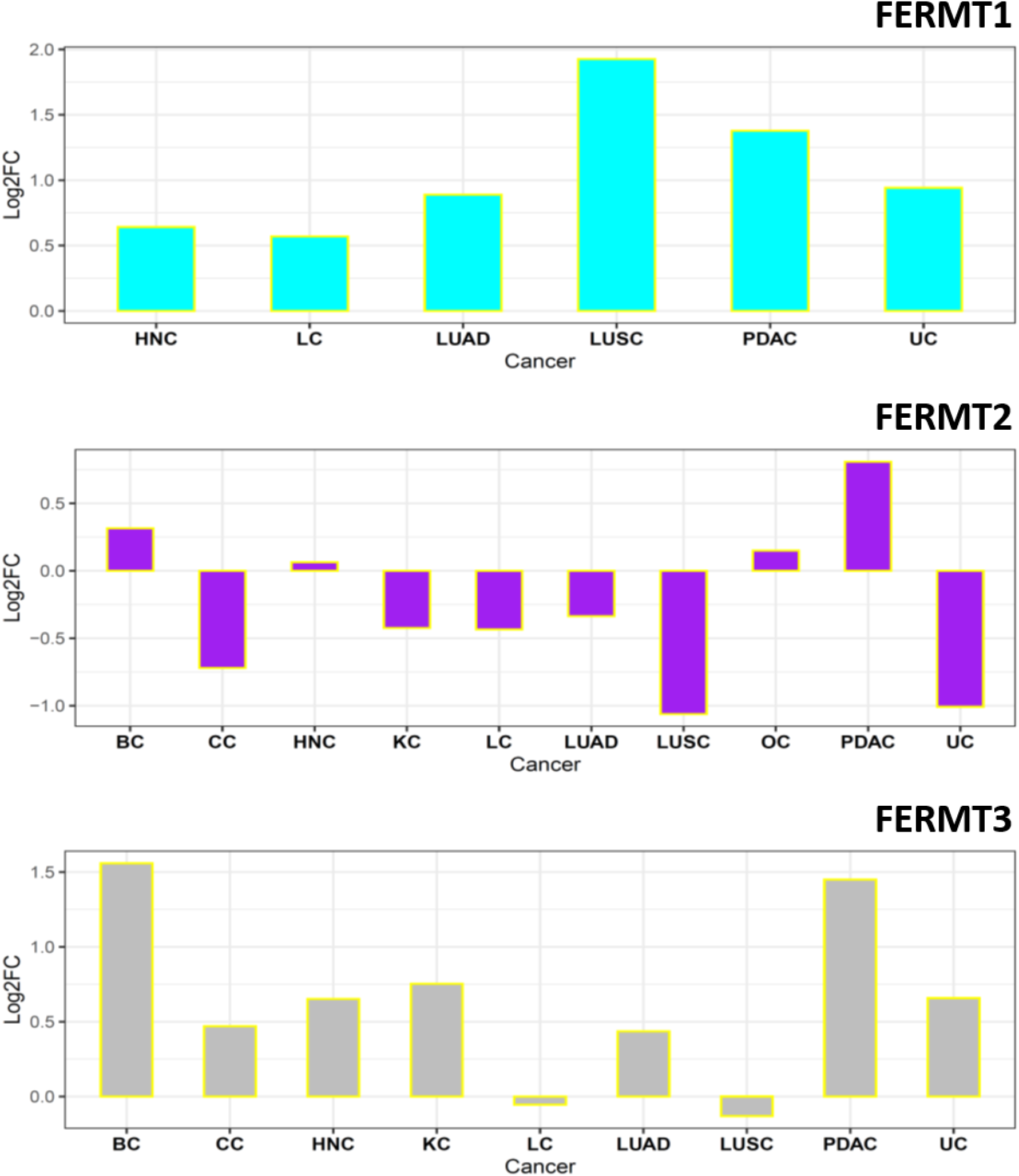
Comparative kindlin phosphorylation analysis of tumor and adjacent tissues. Log2 fold change of tumor (n=1272) phosphorylation levels (TMTlog2 ratio) with that of tumor-adjacent tissues (n=782) for all kindlin family proteins from phosphoproteomic data of CPTAC dataset.

**Fig.S29:**
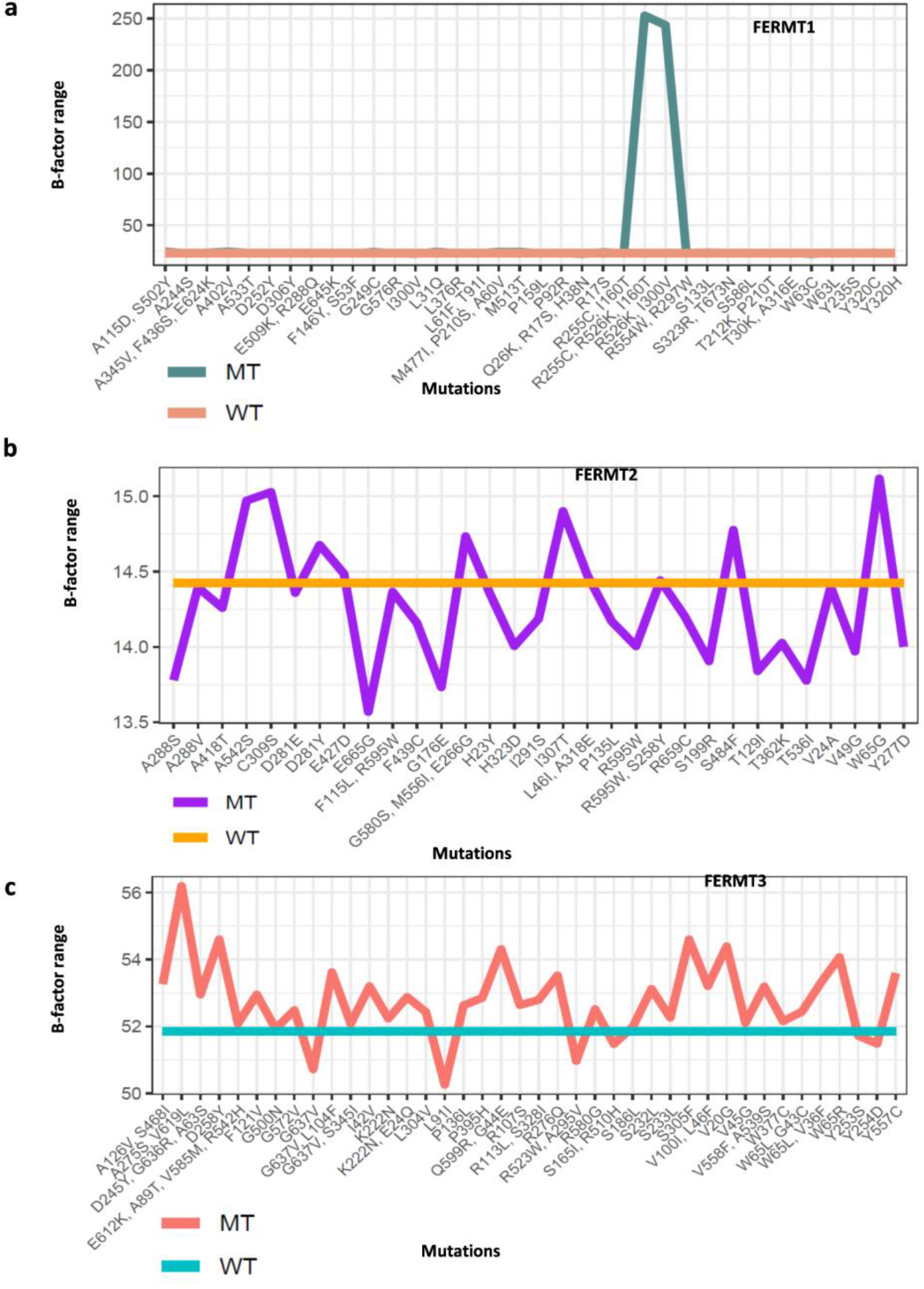
B-Factor Range for Mutations with Respect to Wild Type. a. FERMT1; green, mutated kindlin1; orange, wild type. b. FERMT2; peach, mutated kindlin2; cyan, wild type. c. FERMT3; purple, mutated kindlin3; orange, wild type.

**Fig.S30:**
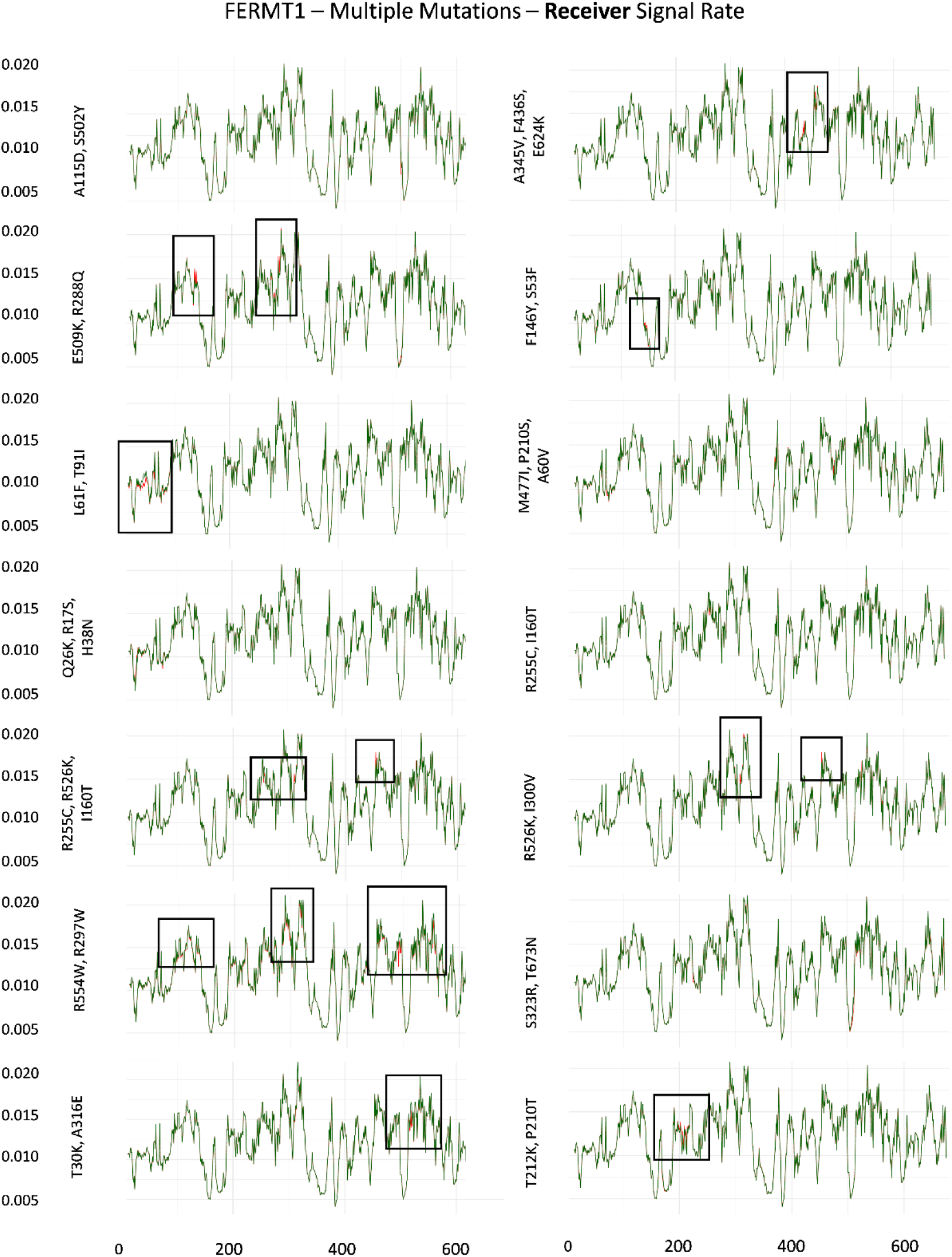
Perturbation in Signal Reception Ability in Multiple Mutated Kindlin1 with Respect to its Wild Type form. The receiver signal rate for every amino acid residue of both, the wild type kindlin1 and the mutated variants is plotted as a line plot. The x-axis represents the amino acid residue index, whereas the y-axis depicts the receiver signal rate, as calculated by Markov’s chain model (distance/hitting time). The mutations are enlisted on the adjacent left of each plot. Green, line plot for wild type kindlin1; orange, line plot for corresponding mutated kindlin1. The boxes within specific line plots highlight a region depicting a visually perturbed trend in signal reception.

**Fig.S31:**
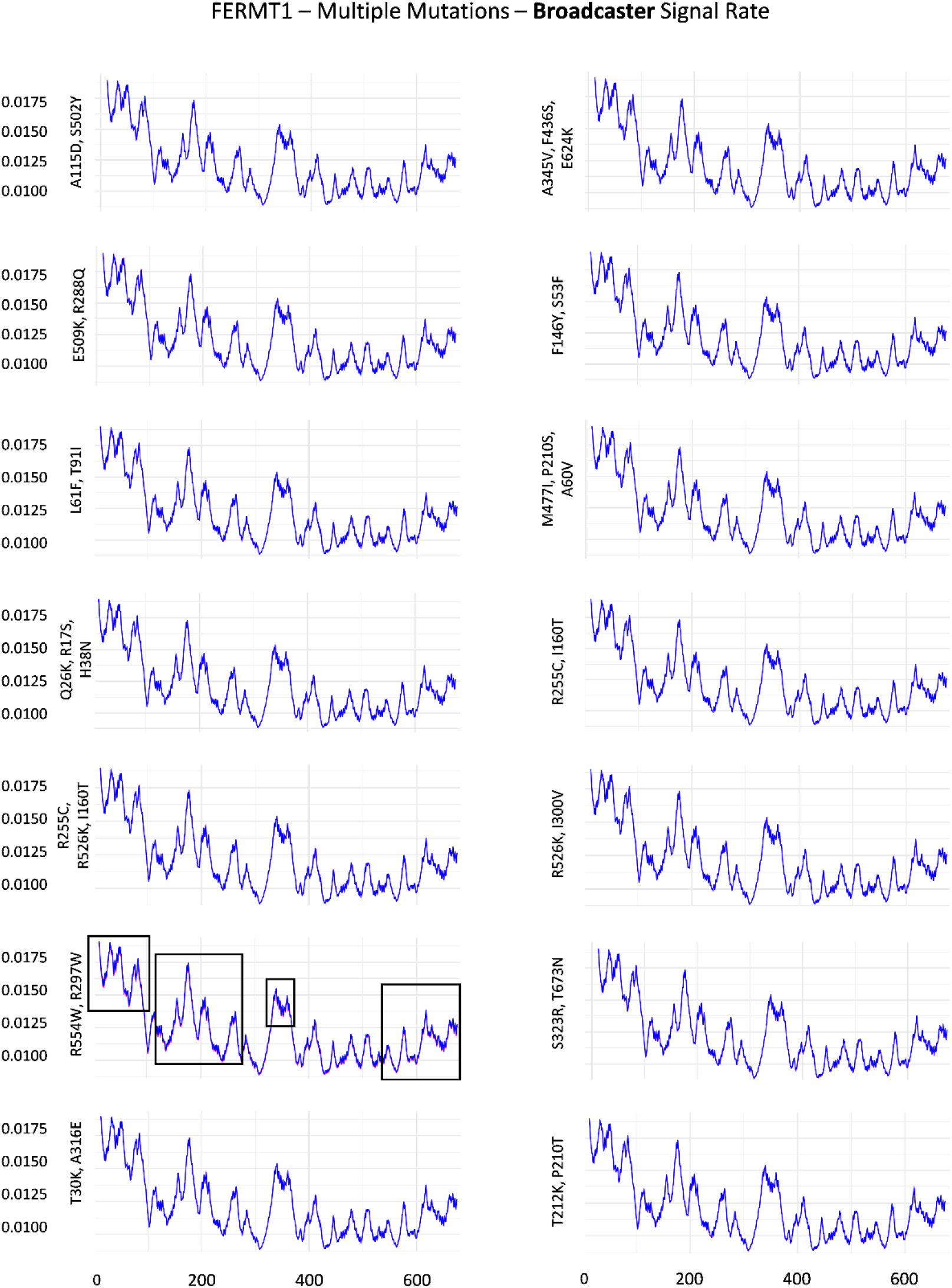
Perturbation in Signal Broadcasting Ability in Multiple Mutated Kindlin1 with Respect to its Wild Type form. The broadcasting signal rate for every amino acid residue of both, the wild type kindlin1 and the mutated variants is plotted as a line plot. The x-axis represents the amino acid residue index, whereas the y-axis depicts the broadcasting signal rate (distance/hitting time), as calculated by Markov’s chain model. The mutations are enlisted on the adjacent left of each plot. Blue, line plot for wild type kindlin1; red, line plot for corresponding mutated kindlin1. The boxes within specific line plots highlight a region depicting a visually perturbed trend in signal reception.

**Fig.S32:**
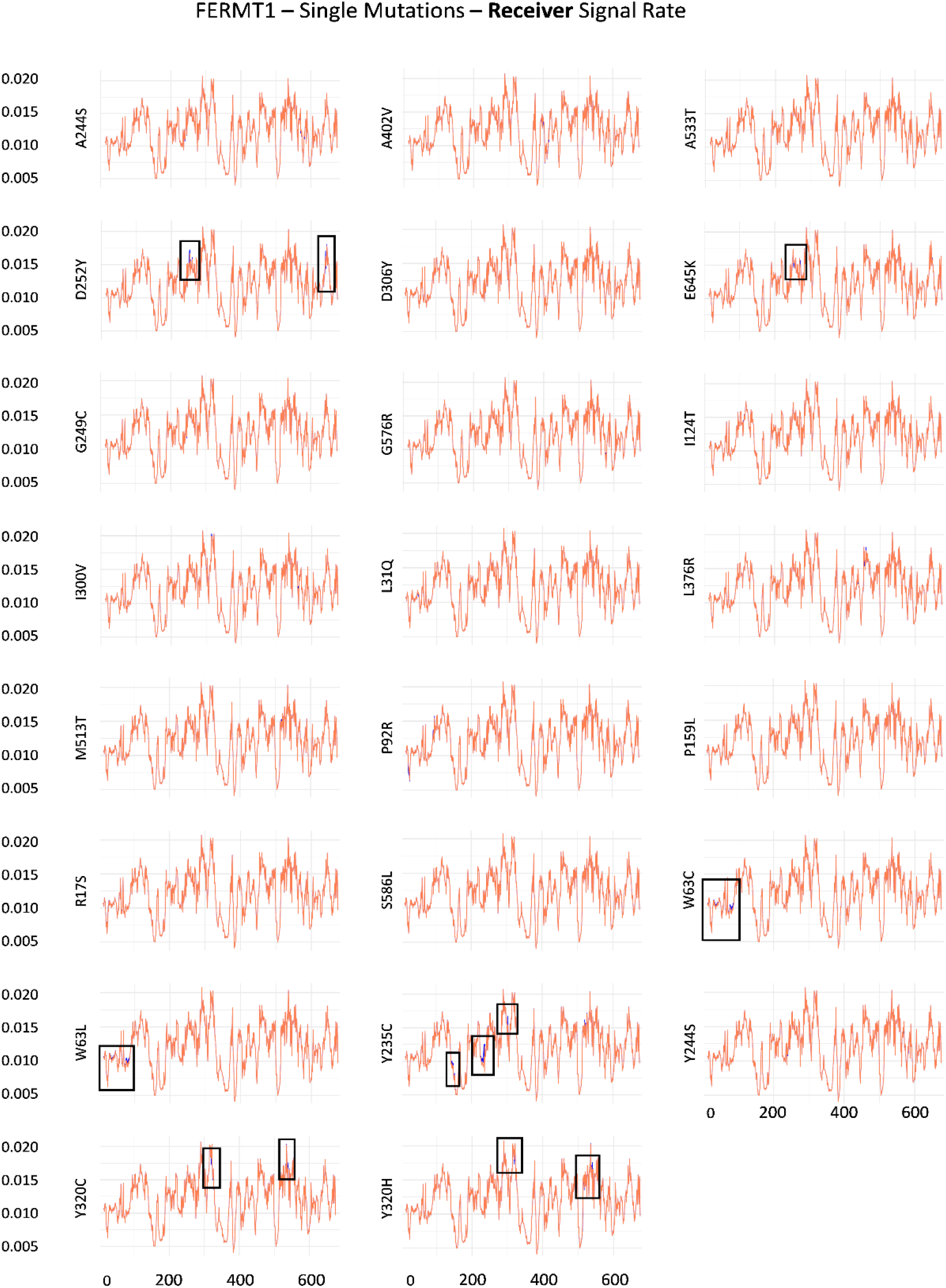
Perturbation in Signal Reception Ability in Single Mutated Kindlin1 with Respect to its Wild Type form. The receiver signal rate for every amino acid residue of both, the wild type kindlin1 and the mutated variants is plotted as a line plot. The x-axis represents the amino acid residue index, whereas the y-axis depicts the receiver signal rate (distance/hitting time), as calculated by Markov’s chain model. The mutations are enlisted on the adjacent left of each plot. Orange, line plot for wild type kindlin1; blue, line plot for corresponding mutated kindlin1. The boxes within specific line plots highlight a region depicting a visually perturbed trend in signal reception.

**Fig.S33:**
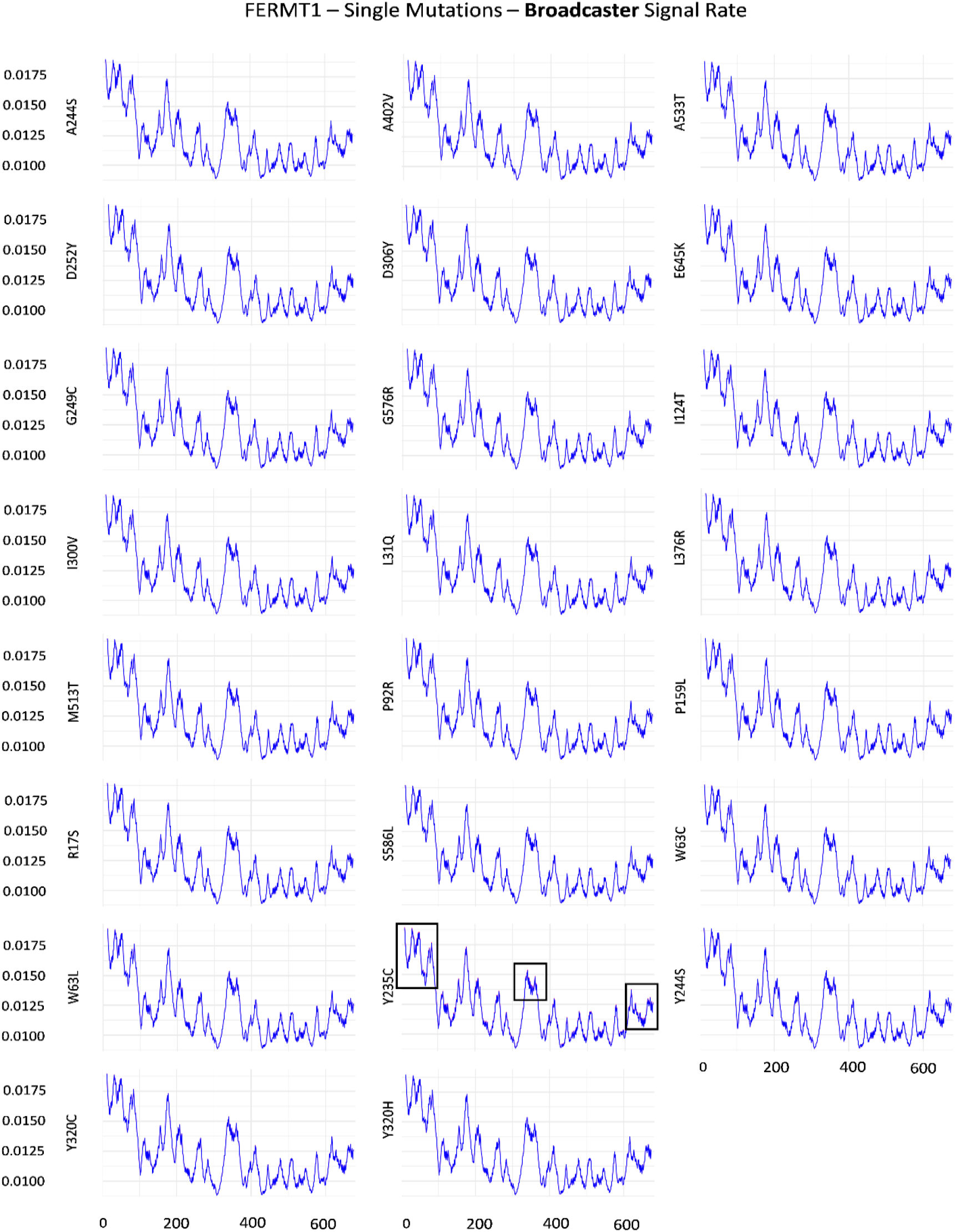
Perturbation in Signal Broadcasting Ability in Single Mutated Kindlin1 with Respect to its Wild Type form. The broadcasting signal rate for every amino acid residue of both, the wild type kindlin1 and the mutated variants is plotted as a line plot. The x-axis represents the amino acid residue index, whereas the y-axis depicts the broadcasting signal rate (distance/hitting time), as calculated by Markov’s chain model. The mutations are enlisted on the adjacent left of each plot. Blue, line plot for wild type kindlin1; red, line plot for corresponding mutated kindlin1. The boxes within specific line plots highlight a region depicting a visually perturbed trend in signal reception.

**Fig.S34:**
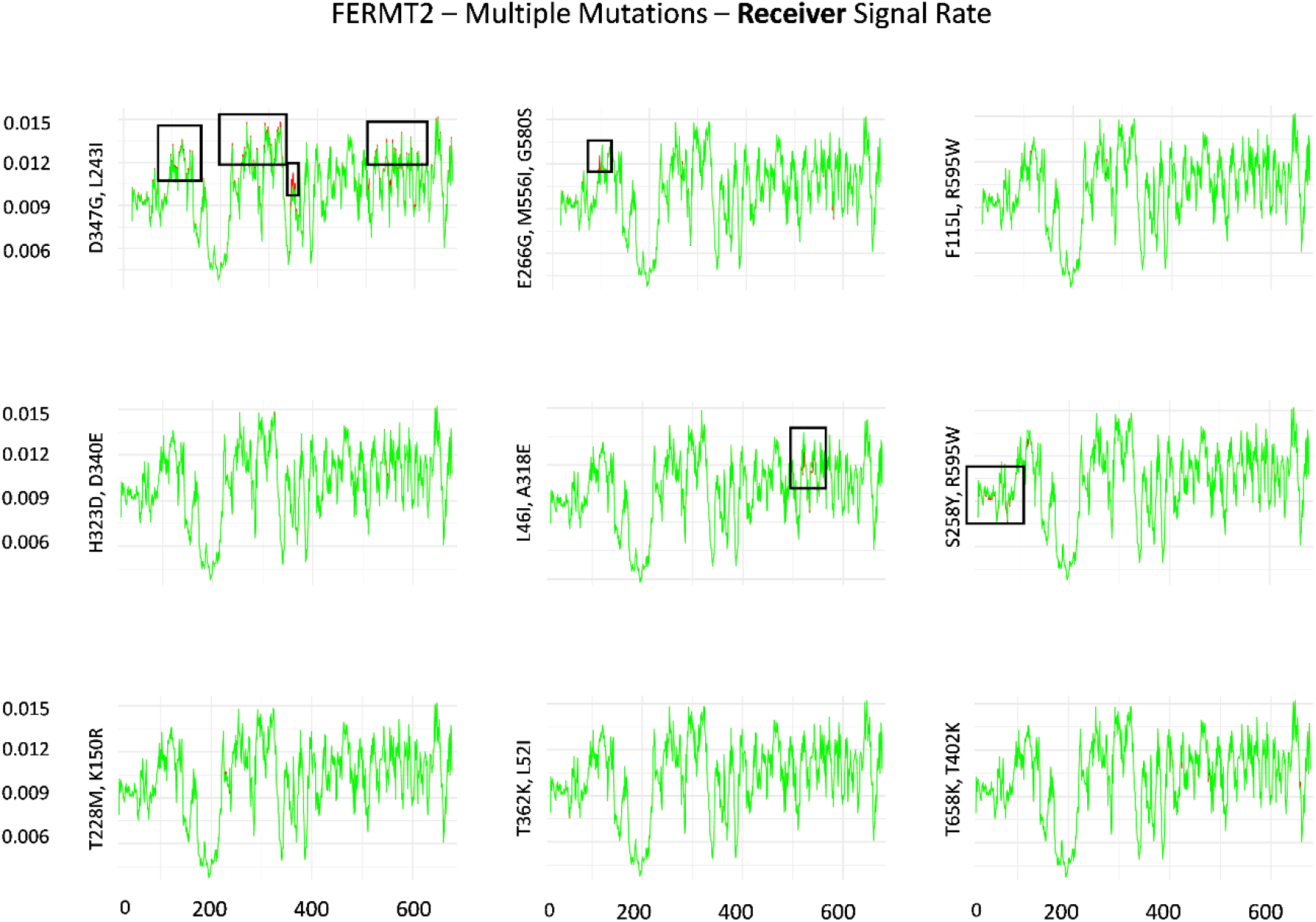
Perturbation in Signal Reception Ability in Multiple Mutated Kindlin2 with Respect to its Wild Type form. The receiver signal rate for every amino acid residue of both, the wild type kindlin2 and the mutated variants is plotted as a line plot. The x-axis represents the amino acid residue index, whereas the y-axis depicts the receiver signal rate (distance/hitting time), as calculated by Markov’s chain model. The mutations are enlisted on the adjacent left of each plot. Green, line plot for wild type kindlin2; red, line plot for corresponding mutated kindlin2. The boxes within specific line plots highlight a region depicting a visually perturbed trend in signal reception.

**Fig.S35:**
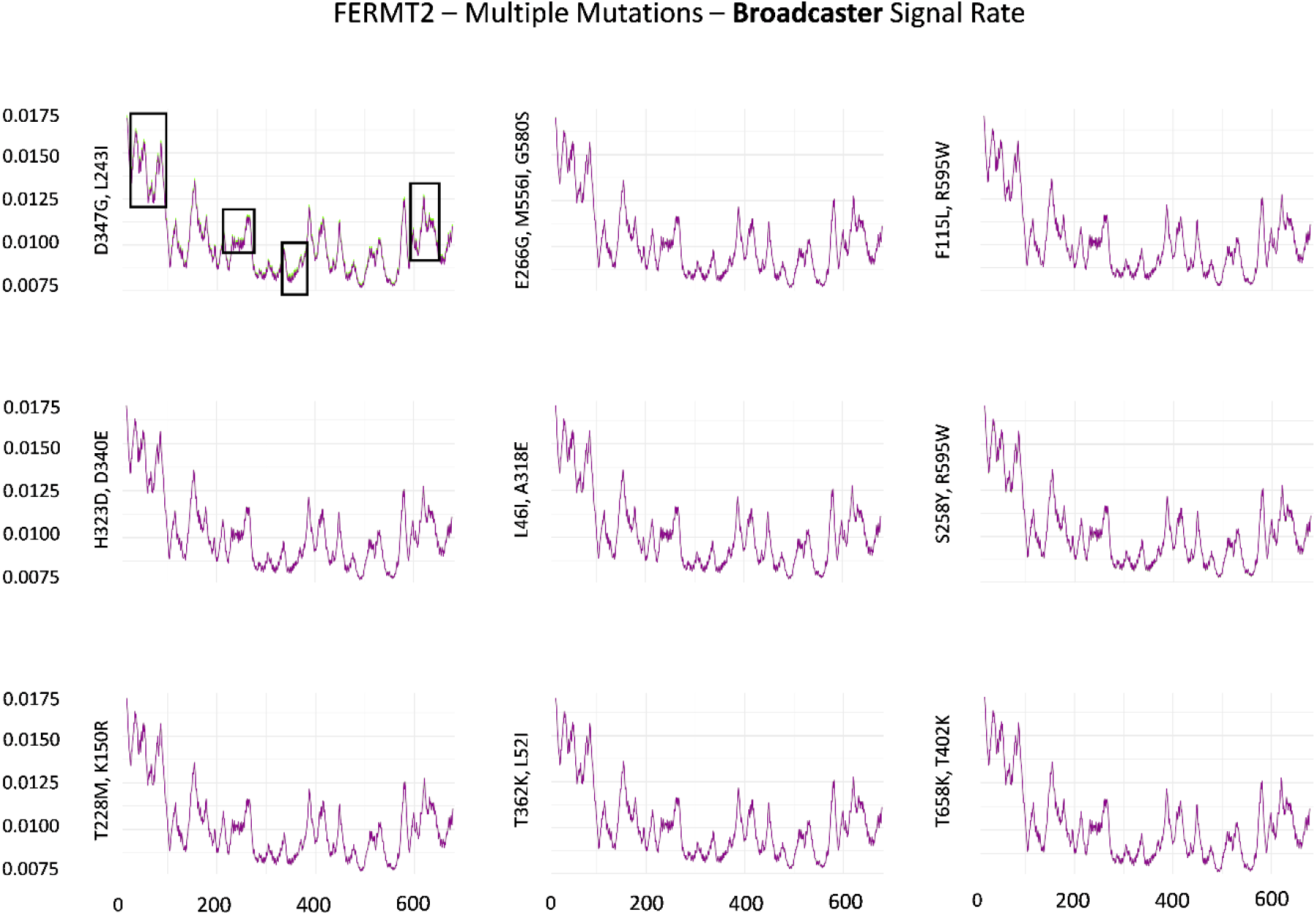
Perturbation in Signal Broadcasting Ability in Multiple Mutated Kindlin2 with Respect to its Wild Type form. The broadcasting signal rate for every amino acid residue of both, the wild type kindlin2 and the mutated variants is plotted as a line plot. The x-axis represents the amino acid residue index, whereas the y-axis depicts the broadcasting signal rate (distance/hitting time), as calculated by Markov’s chain model. The mutations are enlisted on the adjacent left of each plot. Purple, line plot for wild type kindlin2; green, line plot for corresponding mutated kindlin2. The boxes within specific line plots highlight a region depicting a visually perturbed trend in signal reception.

**Fig.S36:**
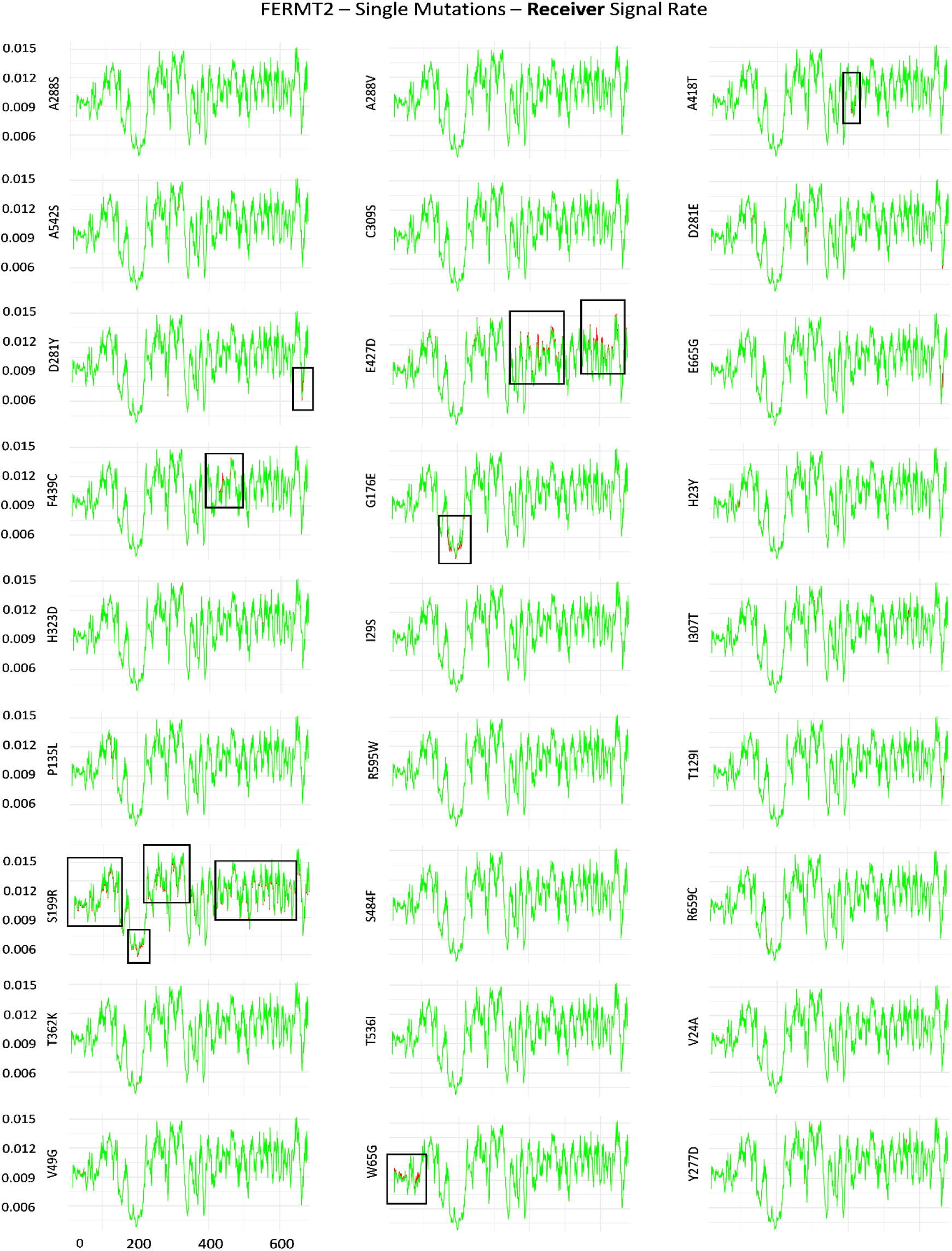
Perturbation in Signal Reception Ability in Single Mutated Kindlin2 with Respect to its Wild Type form. The receiver signal rate for every amino acid residue of both, the wild type kindlin2 and the mutated variants is plotted as a line plot. The x-axis represents the amino acid residue index, whereas the y-axis depicts the receiver signal rate (distance/hitting time), as calculated by Markov’s chain model. The mutations are enlisted on the adjacent left of each plot. Green, line plot for wild type kindlin2; red, line plot for corresponding mutated kindlin2. The boxes within specific line plots highlight a region depicting a visually perturbed trend in signal reception.

**Fig.S37:**
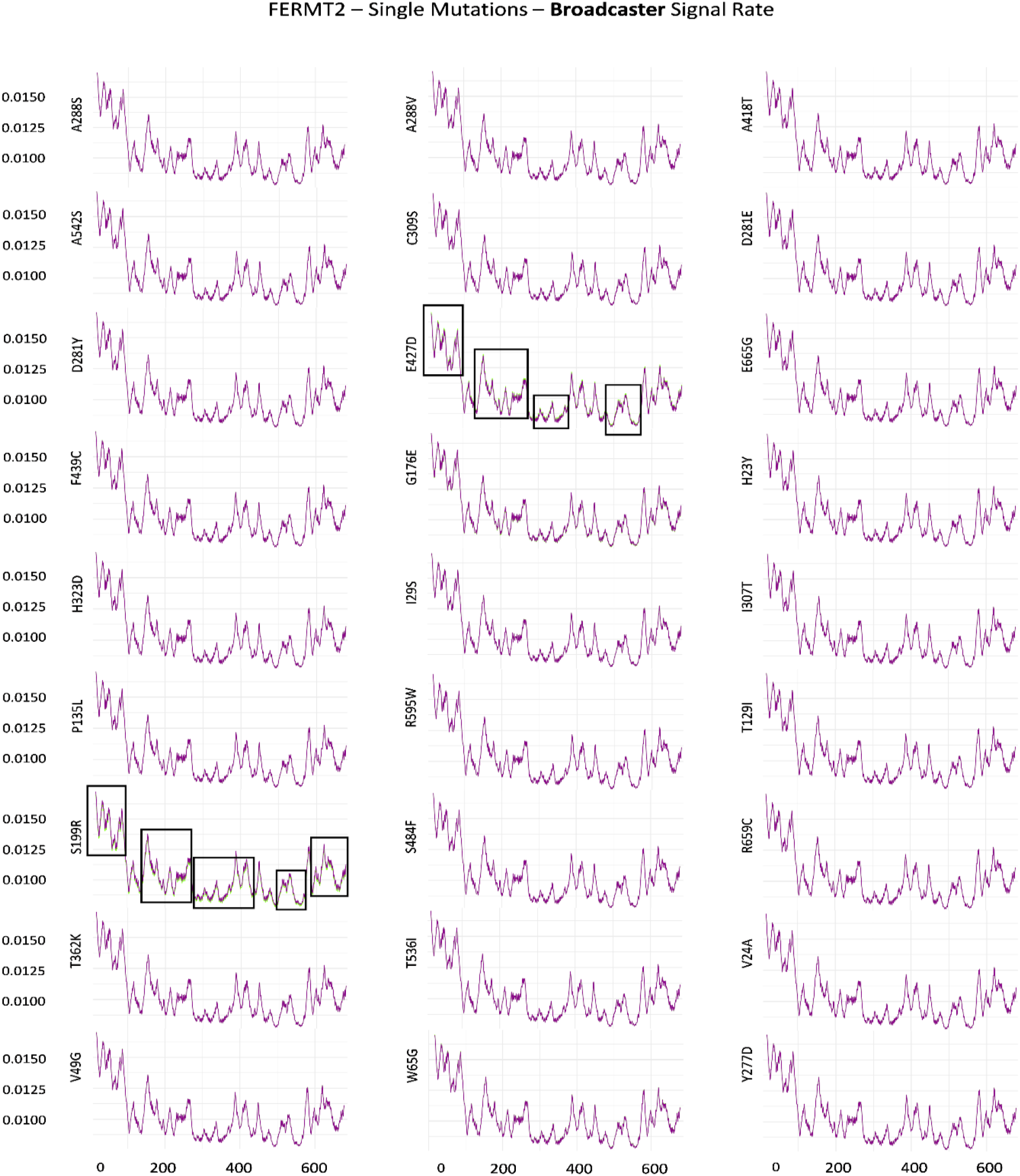
Perturbation in Signal Broadcasting Ability in Single Mutated Kindlin2 with Respect to its Wild Type form. The broadcasting signal rate for every amino acid residue of both, the wild type kindlin2 and the mutated variants is plotted as a line plot. The x-axis represents the amino acid residue index, whereas the y-axis depicts the broadcasting signal rate (distance/hitting time), as calculated by Markov’s chain model. The mutations are enlisted on the adjacent left of each plot. Purple, line plot for wild type kindlin2; green, line plot for corresponding mutated kindlin2. The boxes within specific line plots highlight a region depicting a visually perturbed trend in signal reception.

**Fig.S38:**
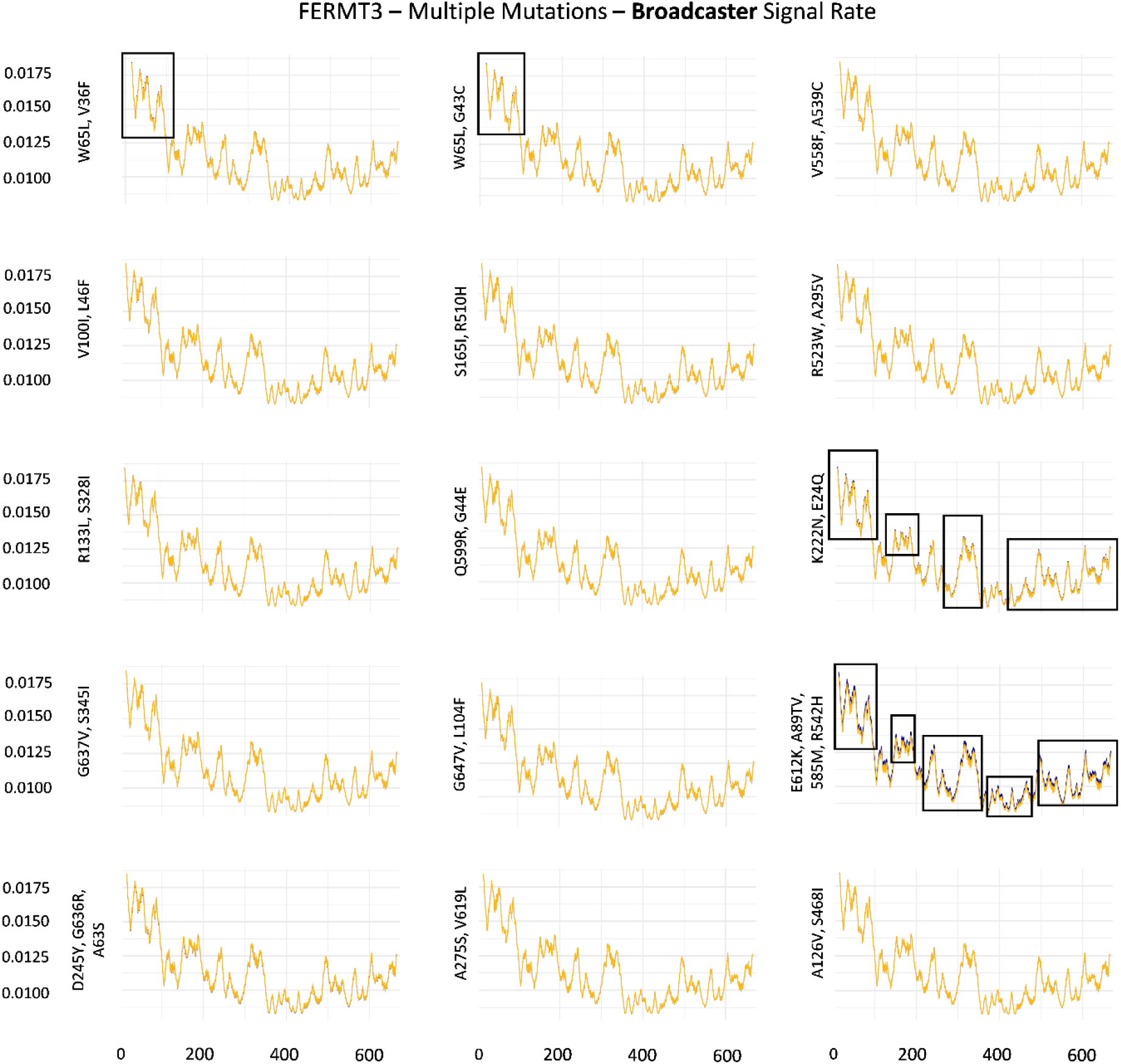
Perturbation in Signal Broadcasting Ability in Multiple Mutated Kindlin3 with Respect to its Wild Type form. The broadcasting signal rate for every amino acid residue of both, the wild type kindlin3 and the mutated variants is plotted as a line plot. The x-axis represents the amino acid residue index, whereas the y-axis depicts the broadcasting signal rate (distance/hitting time), as calculated by Markov’s chain model. The mutations are enlisted on the adjacent left of each plot. Yellow, line plot for wild type kindlin3; blue, line plot for corresponding mutated kindlin3. The boxes within specific line plots highlight a region depicting a visually perturbed trend in signal reception.

**Fig.S39:**
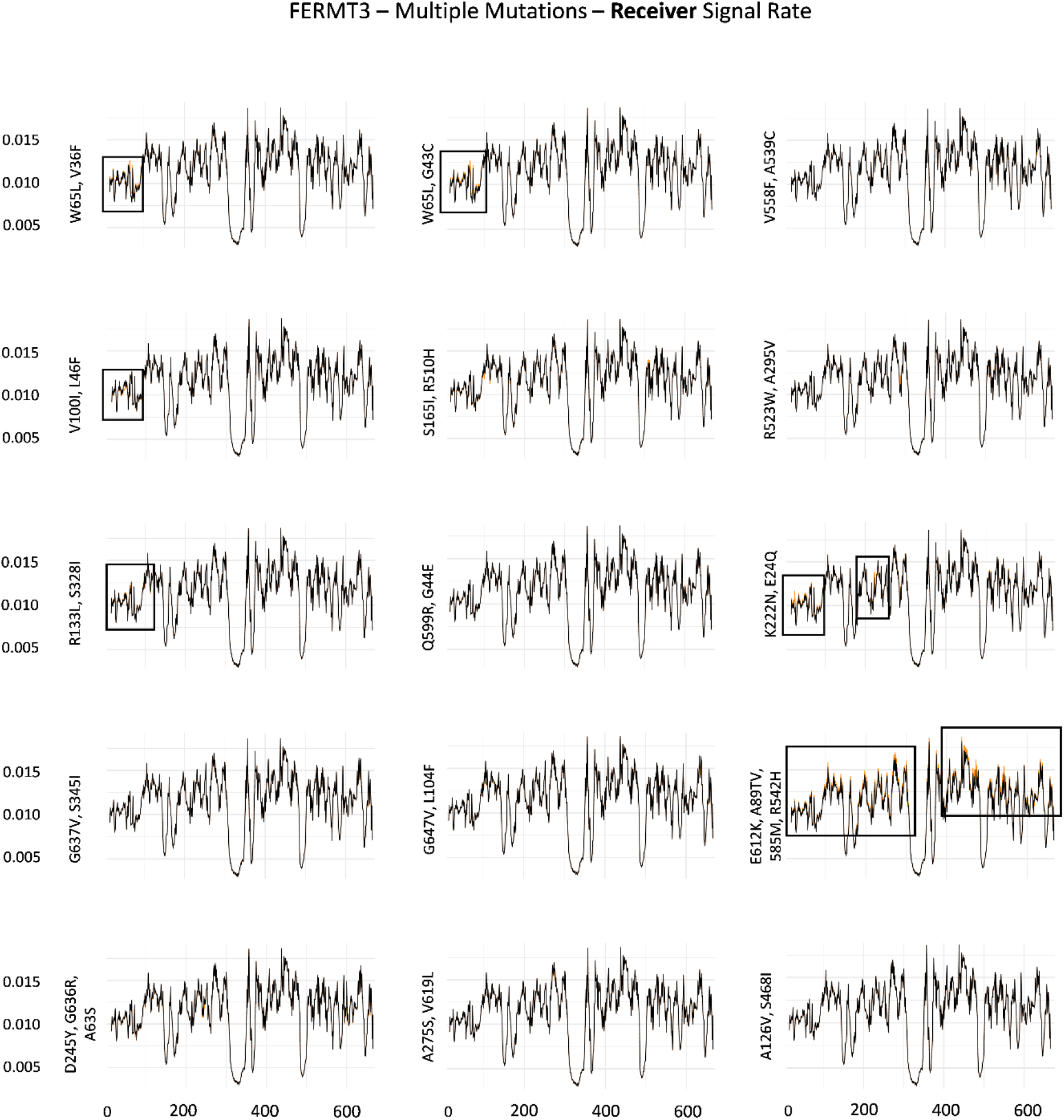
Perturbation in Signal Reception Ability in Multiple Mutated Kindlin3 with Respect to its Wild Type form. The receiver signal rate for every amino acid residue of both, the wild type kindlin3 and the mutated variants is plotted as a line plot. The x-axis represents the amino acid residue index, whereas the y-axis depicts the receiver signal rate (distance/hitting time), as calculated by Markov’s chain model. The mutations are enlisted on the adjacent left of each plot. Black, line plot for wild type kindlin3; orange, line plot for corresponding mutated kindlin3. The boxes within specific line plots highlight a region depicting a visually perturbed trend in signal reception.

**Fig.S40:**
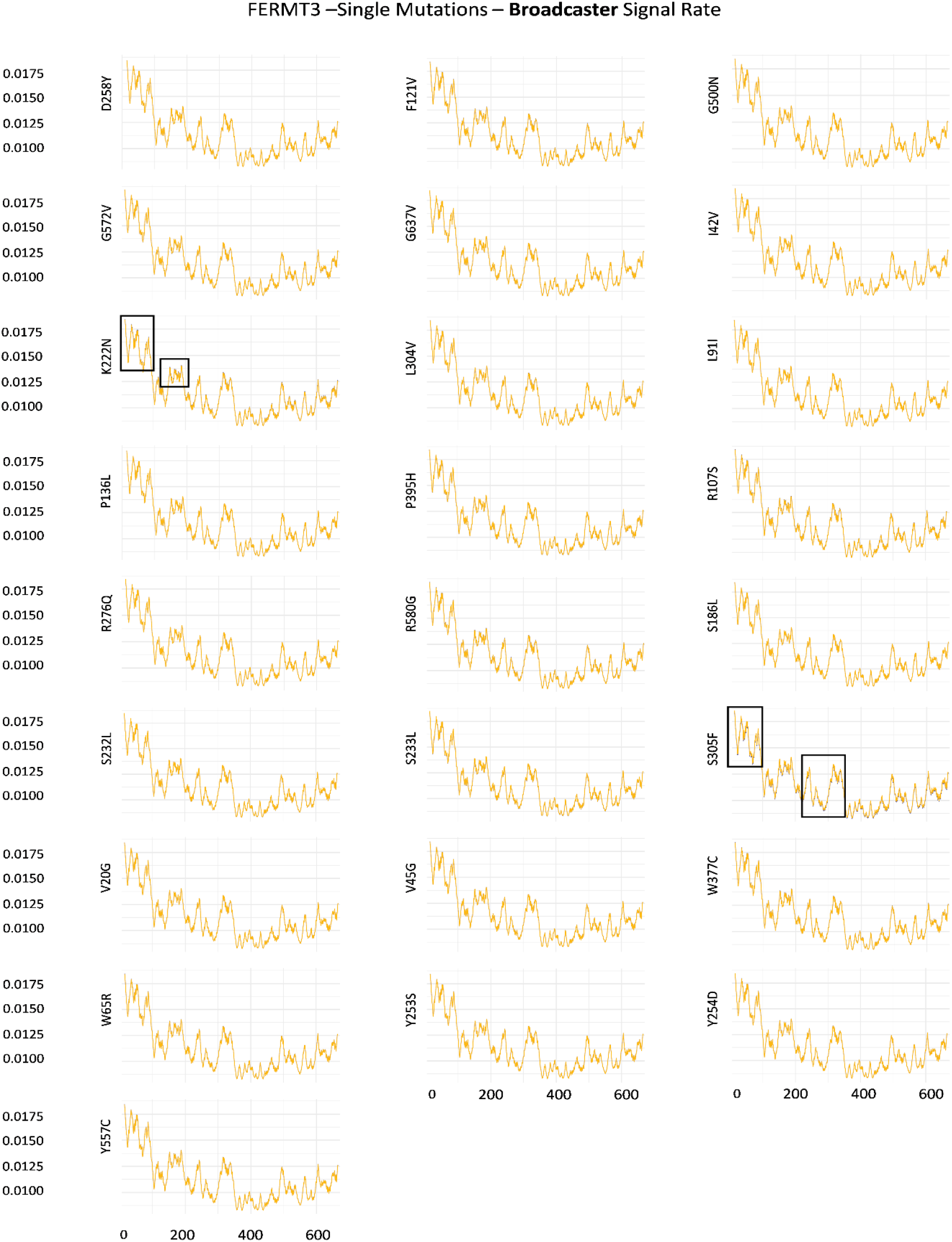
Perturbation in Signal Broadcasting Ability in Single Mutated Kindlin3 with Respect to its Wild Type form. The broadcasting signal rate for every amino acid residue of both, the wild type kindlin3 and the mutated variants is plotted as a line plot. The x-axis represents the amino acid residue index, whereas the y-axis depicts the broadcasting signal rate (distance/hitting time), as calculated by Markov’s chain model. The mutations are enlisted on the adjacent left of each plot. Yellow, line plot for wild type kindlin3; blue, line plot for corresponding mutated kindlin3. The boxes within specific line plots highlight a region depicting a visually perturbed trend in signal reception.

**Fig.S41:**
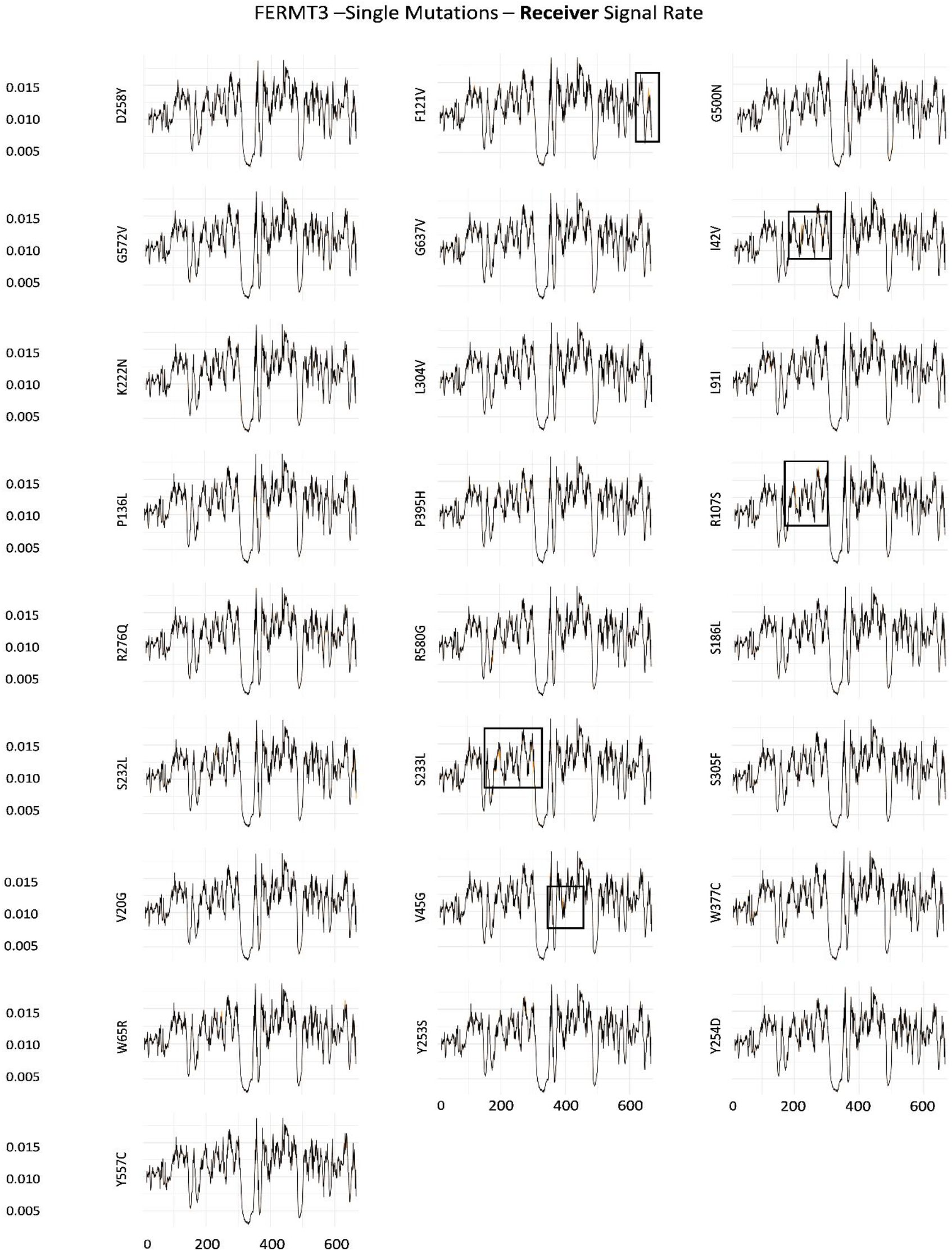
Perturbation in Signal Reception Ability in Single Mutated Kindlin3 with Respect to its Wild Type form. The receiver signal rate for every amino acid residue of both, the wild type kindlin3 and the mutated variants is plotted as a line plot. The x-axis represents the amino acid residue index, whereas the y-axis depicts the receiver signal rate (distance/hitting time), as calculated by Markov’s chain model. The mutations are enlisted on the adjacent left of each plot. Black, line plot for wild type kindlin3; Orange, line plot for corresponding mutated kindlin3. The boxes within specific line plots highlight a region depicting a visually perturbed trend in signal reception.

**Fig.S42:**
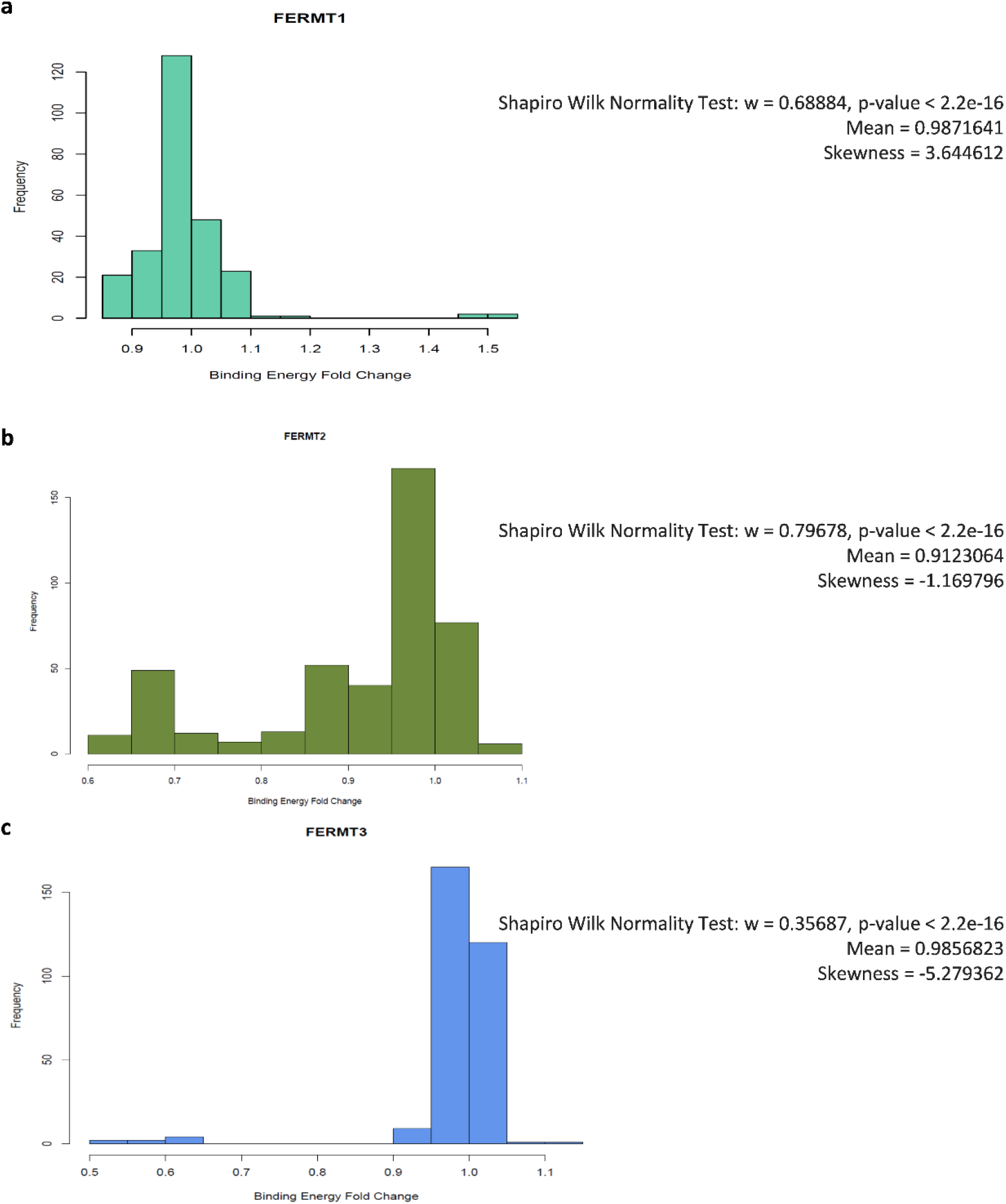
Histogram Showing Distribution in Fold Change in Binding Affinity of Kindlin Mutants for their Direct Interactors with respect to that of Wild Type. a. FERMT1; b. FERMT2; c. FERMT3. w, normality value; p-value indicative of all data being statistically significant; skewness: positive, right skewed; negative, left skewed. Binding affinity fold-change value > 1 signifies increased affinity, or interaction probability, whereas < 1 signifies decreased affinity towards interaction.

**Fig.S43:**
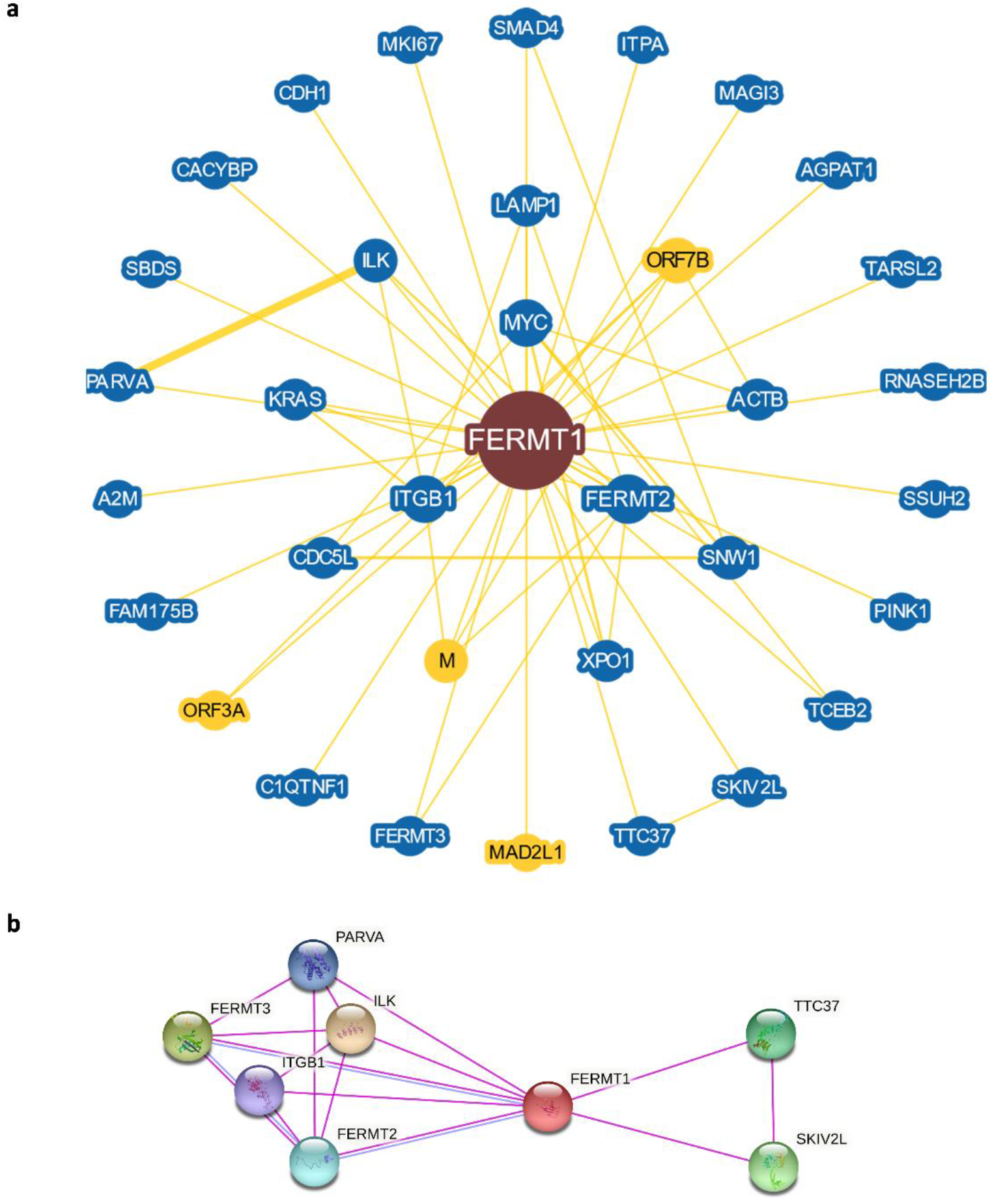
FERMT1 Interactome. a. FERMT1 protein associated network. Proteins represented by the different nodes with the query protein in the middle. Blue nodes signify associated gene from same organism while yellow shows association from different organism. Yellow lines represent connection through physical evidence. Larger node size represents a higher interaction and thicker connections show stronger evidence for association. b. Protein-protein interaction network for FERMT1. Nodes represent proteins while lines show interactions between the proteins. Associations sourced from experiments and databases denoted by the pink and blue lines respectively. Interaction score of medium confidence (0.400) employed. The distance of the lines indicates the interaction score in an inverse relation.

**Fig.S44:**
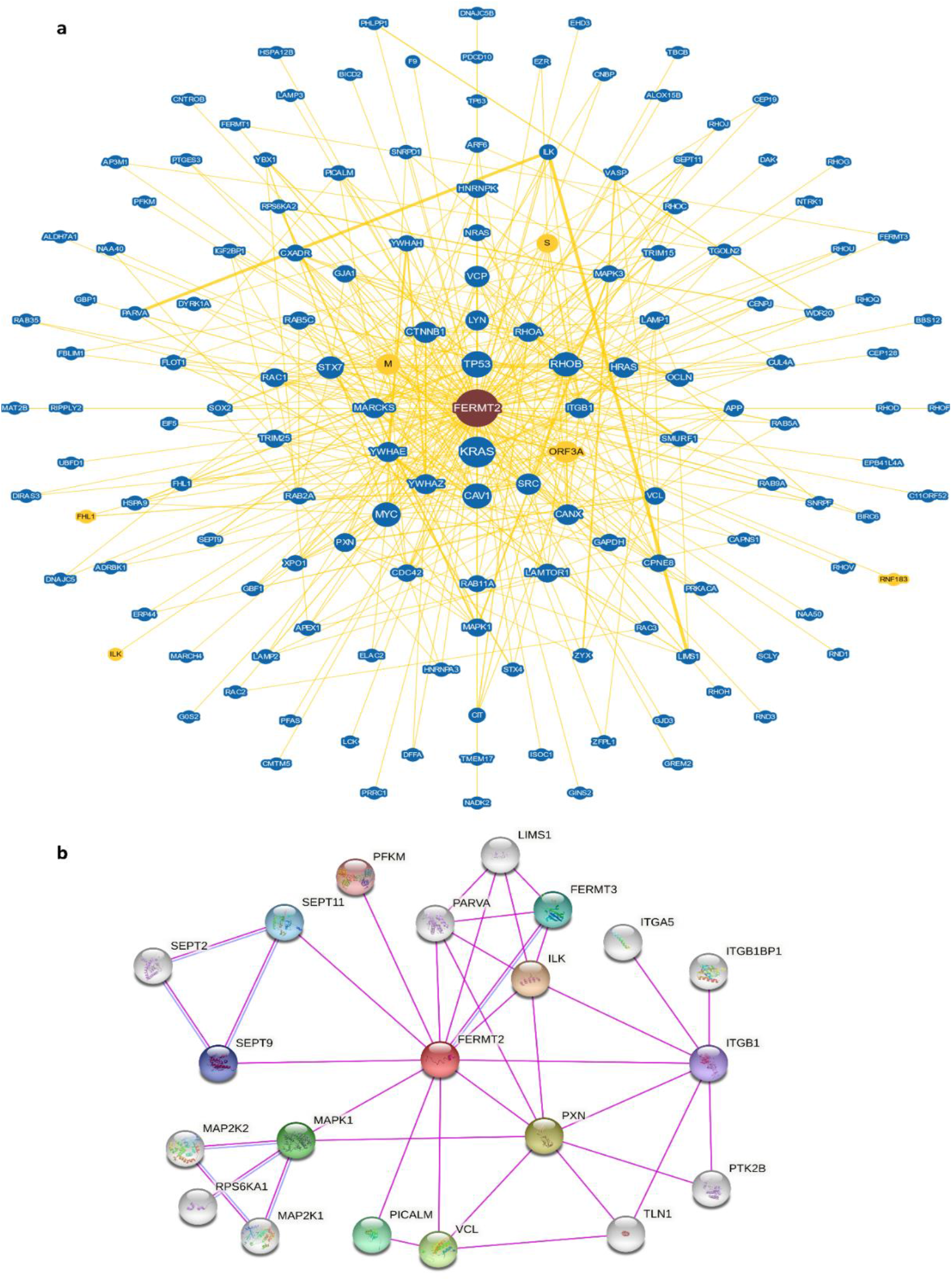
FERMT2 Interactome. a. FERMT2 protein associated network. Proteins represented by the different nodes with the query protein in the middle. Blue nodes signify associated gene from same organism while yellow shows association from different organism. Yellow lines represent connection through physical evidence. Larger node size represents a higher interaction and thicker connections show stronger evidence for association. b. Protein-protein interaction network for FERMT2. Nodes represent proteins while lines show interactions between the proteins. Associations sourced from experiments and databases denoted by the pink and blue lines respectively. Interaction score of medium confidence (0.400) employed. The distance of the lines indicates the interaction score in an inverse relation.

**Fig.S45:**
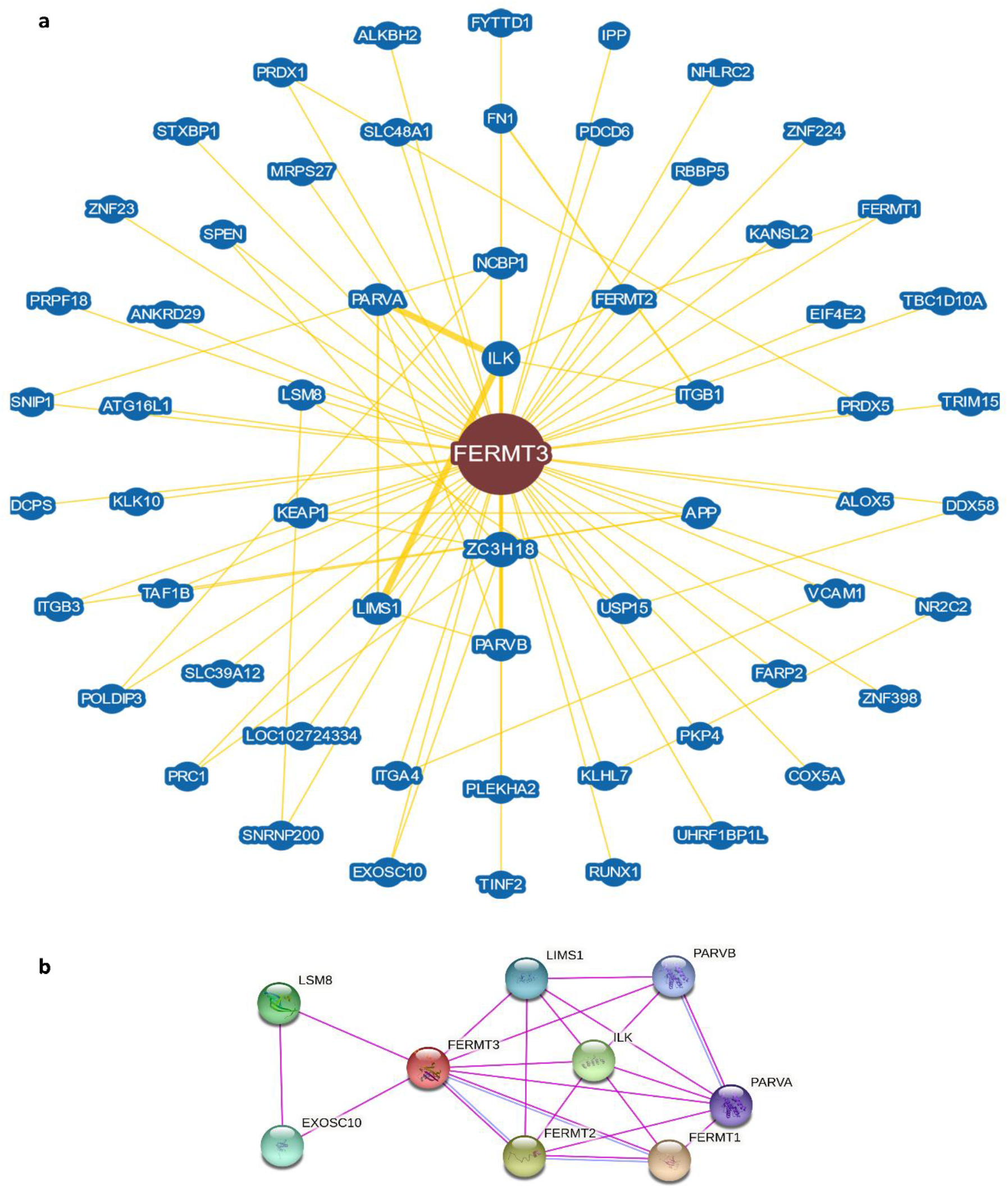
FERMT3 Interactome. a. FERMT3 protein associated network. Proteins represented by the different nodes with the query protein in the middle. Blue nodes signify associated gene from same organism while yellow shows association from different organism. Yellow lines represent connection through physical evidence. Larger node size represents a higher interaction and thicker connections show stronger evidence for association. b. Protein-protein interaction network for FERMT3. Nodes represent proteins while lines show interactions between the proteins. Associations sourced from experiments and databases denoted by the pink and blue lines respectively. Interaction score of medium confidence (0.400) employed. The distance of the lines indicates the interaction score in an inverse relation.

**Fig.S46:**
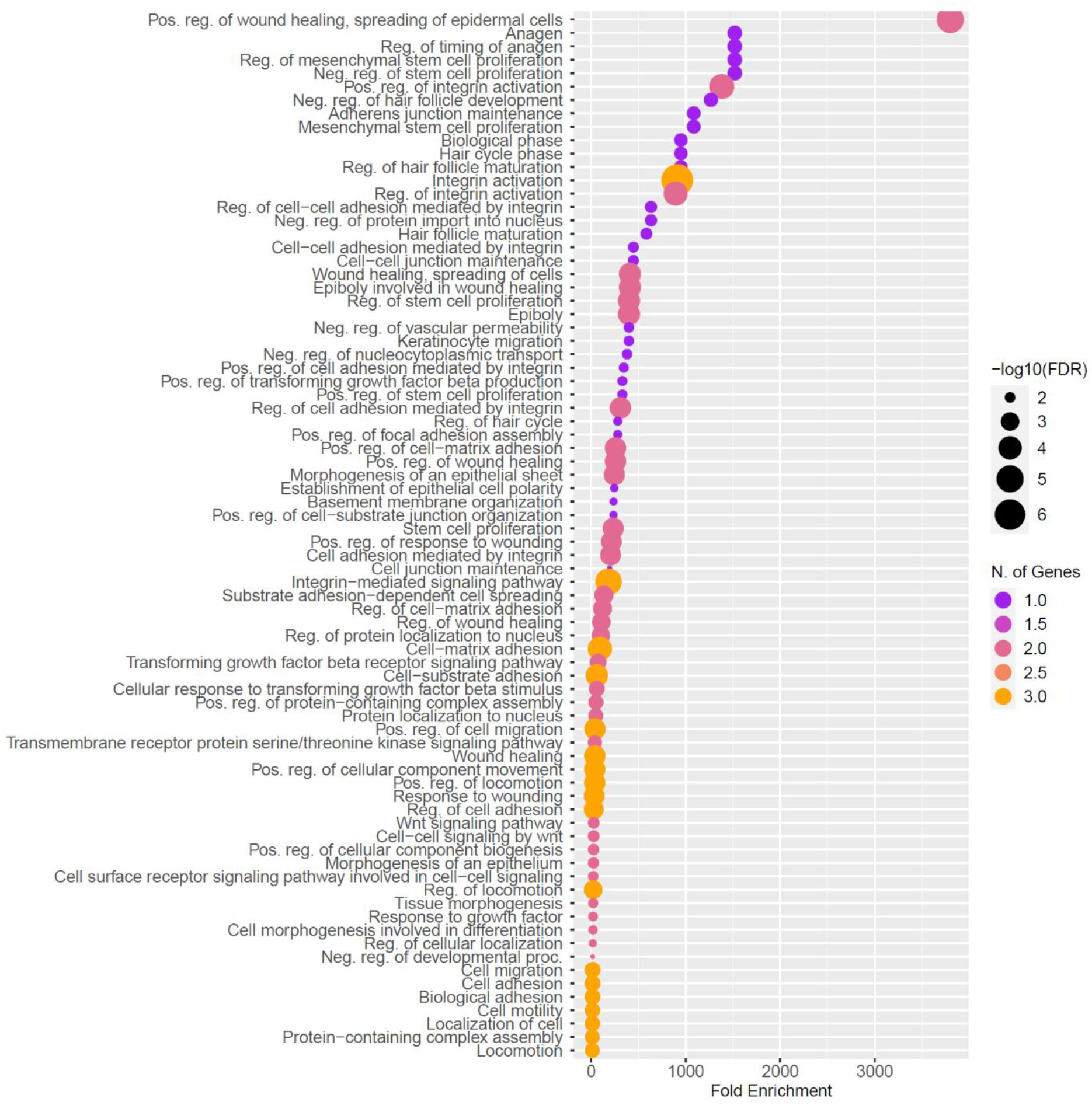
Pathway Enrichment Analysis for Kindlins. The biological processes are written towards the left, and their fold-enrichment are plotted in the x-axis. The color of the points indicates the number of genes (FERMT1, FERMT2, and FERMT3) involved. The size of the points is set according to –Log_10_ (FDR) value. Top 100 biological pathways are sorted based on the FDR cut-off (0.05).

**Fig.S47:**
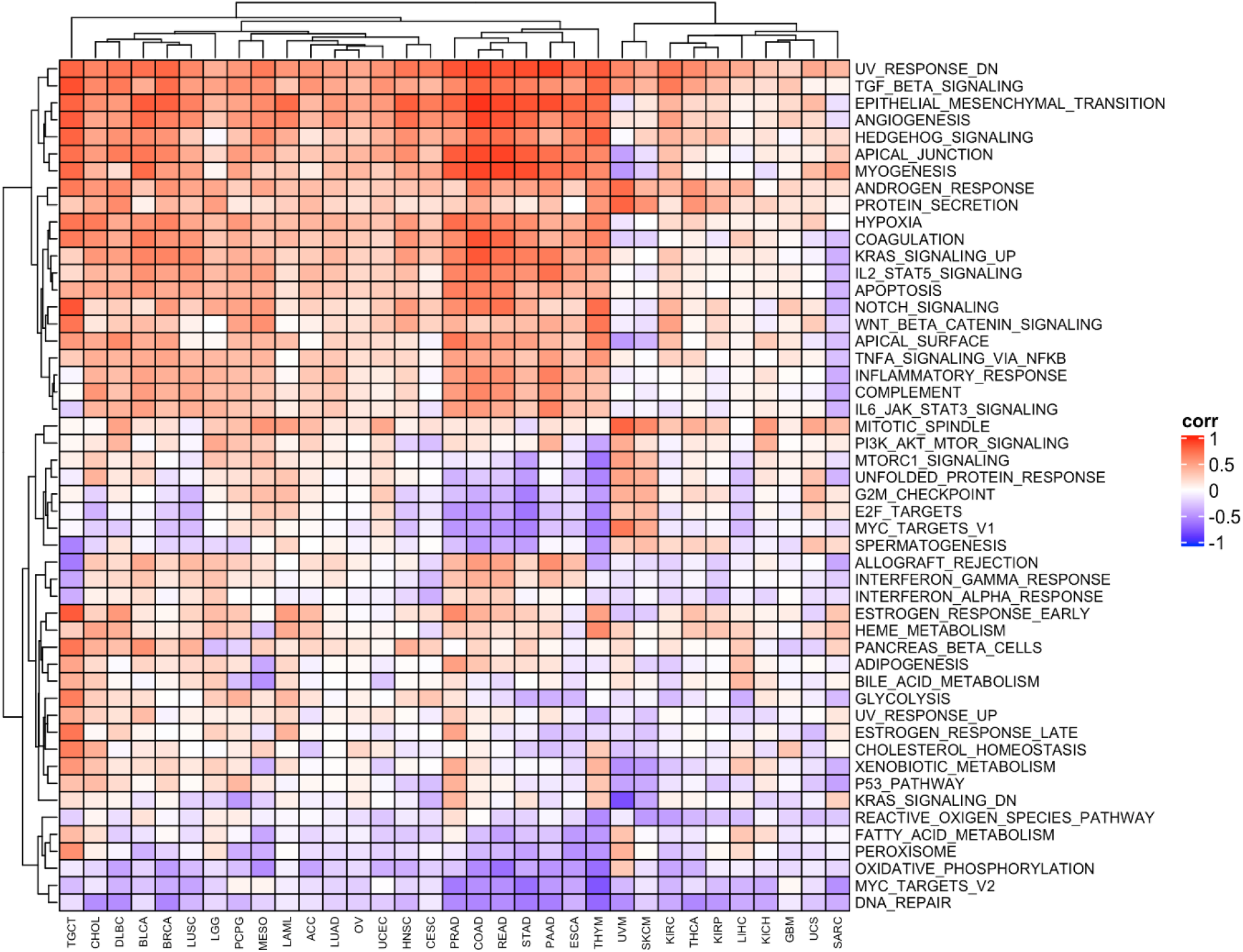
FERMT2 Gene Expression Association with EMT Signature Biological Events. Colors are indicative of the degree of correlation, where red signifies correlation, and purple signifies anti-correlation, respectively. The color intensity is proportional to the extent of correlation or anti-correlation, as represented by number, where 1 represents highest correlation, and -1 represents highest anti-correlation.

**Fig.S48:**
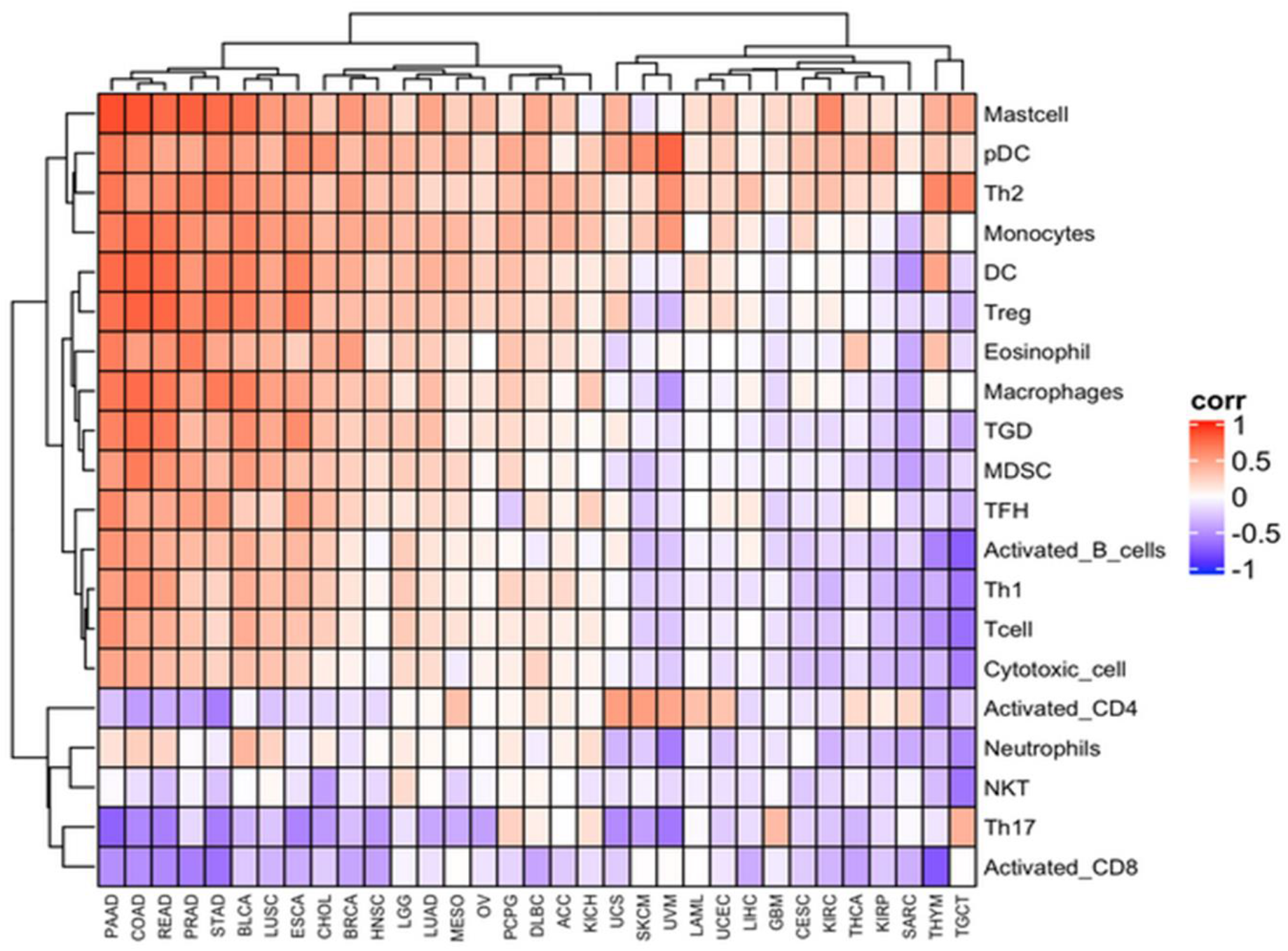
FERMT2 Gene Expression Association with Immune Cell Profiling Signature. Colors are indicative of the degree of correlation, where red signifies correlation, and purple signifies anti-correlation, respectively. The color intensity is proportional to the extent of correlation or anti-correlation, as represented by number, where 1 represents highest correlation, and -1 represents highest anti-correlation.

**Fig.S49:**
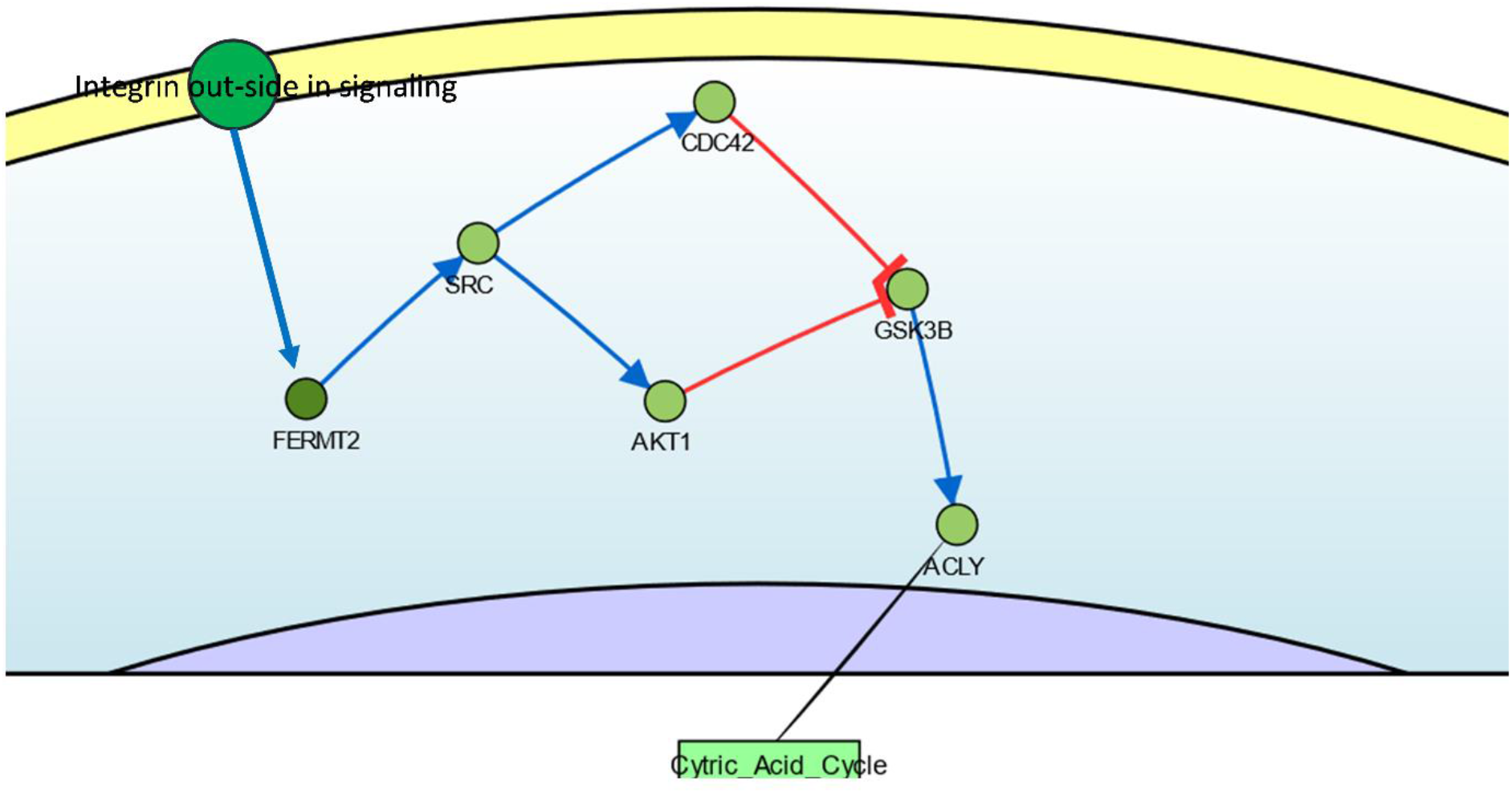
Kindlin2 as a connecting link between Integrin Outside-in Signaling & Citric Acid Cycle. The image is taken and modified from CancerGeneNet. Extra-cellular mechanical signals in cancer cells activate FERMT2 via Integrins outside-in signaling. Activated FERMT2 further activates SRC, and consequently CDC42 and AKT1 downstream. The activation of CDC42 and AKT1 inhibits GSK3B, which otherwise upregulates the citric acid cycle via ACLY. Blue arrows indicate activation; red lines indicate inhibition; the node distance between FERMT2-SRC = 0.332; SRC-CDC42 = 0.645; SRC-AKT1 = 0.683; CDC42-GSK3B = 0.359; AKT1-GSK3B = 0.834; GSK3B-ACLY = 0.379. The overall Z-score through CDC42 = -2.014; through AKT1 = -2.746

### Supplementary Note 1

#### Supplementary Methodology

##### Databases used in this study

The Cancer Genome Atlas is a comprehensive curation of cancer samples (from upwards of 11,000 patients, or upwards of 20,000 primary and matched normal samples) for over 15 years spanning through 33 cancer types. This data is primarily housed on cBioPortal (https://www.cbioportal.org/)^1,2^. Similarly, the Sanger Cancer Genome Project is based out of the Wellcome Trust Sanger Institute’s cancer, aging, and somatic mutations research group characterizing the role of genomic alterations in the life history of human cancers. The Catalogue of Somatic Mutations in Cancer (COSMIC) now collects data not just from the Wellcome Sanger Institute, but also from TCGA and upwards of 30,000 peer-reviewed primary literature (https://cancer.sanger.ac.uk/cosmic)^3^. The International Cancer Genome Consortium is an international body that coordinates with and houses the data from multiple, large-scale cancer genomic projects conducted worldwide on the ICGC Data Portal (https://dcc.icgc.org/)^4^. Within the ICGC, a project to identify and characterize mutations in ∼3000 cancer whole genomes was undertaken – The Pan-cancer Analysis of Whole Genomes (PCAWG) study^5^. The ICGC Data Portal and cBioportal houses all the PCAWG datasets.

Genomic data acquisition and analysis for our study was primarily from the PCAWG database, the Sanger Cancer Genome Project, and TCGA. Comparative transcriptomic analysis was performed using the TCGA and GTEx databases for cancer and non-cancer samples, respectively, along with the Gene Expression Omnibus (GEO)^6^ employing the GEPIA2 (http://gepia2.cancer-pku.cn/#index)^7^ and GENT2 portals (http://gent2.appex.kr/gent2/)^8^. Proteomics analysis for cancer and non-cancer cohorts was performed by employing the Genomic Data Commons’ (GDC)^9^ Clinical Proteomic Tumor Analysis Consortium (CPTAC, NCI/NIH) database, which incorporates TCGA and TARGET (Therapeutically Applicable Research to Generate Effective Therapies) libraries on cProSite (https://cprosite.ccr.cancer.gov/). Shortlisting of relevant MicroRNAs and their expression analysis in the context of cancer relied on the systematic analysis and text mining of high-throughput experimental data and primary literature on miRDB (http://www.mirdb.org/)^10^ and miRCancer (http://mircancer.ecu.edu/)^11^.

Protein-protein interactions and extended interactomes were characterized employing text-mining strategies across high-throughput datasets and primary literature, along with individual focused studies, and previously established databases on the STRING portal (https://string-db.org/)^12^ and BioGRID (https://thebiogrid.org/)^13^, respectively. The epithelial-to-mesenchymal-specific interactome was analysed employing EMT core signatures from existing literature, obtaining cell line transcriptomic data from the Cancer Cell Line Encyclopaedia (CCLE)^14^, and multi-omics profiles from the TCGA database and MET500^15^ on the EMTome portal (http://www.emtome.org/)^16^.

##### Copy Number Variation analysis

CNV data of FERMT1, 2 and 3 were collected from NCI GDC and ICGC PCAWG cohort. For the CNV module, the percentage of different types of CNV and CNV correlation with mRNA expression from gene in each cancer type were calculated. The raw data were processed with GISTICS2.0^17^ to get the Cancer-specific CNV statistics. However, we calculated the correlation between CNV and mRNA expression based on CNV raw data with sample specific mRNA expressions of concerned genes. The variations were divided into 2 subtypes-homozygous and heterozygous CNV. The later represent the CNV occurrence on only one chromosome whereas the former indicates CNV occurrence in both chromosomes. Percentage statistics were obtained based on CNV subtypes used, GISTIC processed data, and correlation calculations were done using raw CNV and mRNA RSEM data.

##### Methylation data analysis

Cancer-specific methylation data were obtained from NCI Genomic Data Commons (33 cancer types), across them only 14 cancer types contain paired tumor vs. normal data for FERMT1, FERMT2 and FERMT3. Based on these 14 cancer types we built the differential methylation analysis. Cancers with at least 10 tumor-normal pairs are considered for tumor vs normal methylation difference calculation, with only paired samples being included. Student t-test were performed to statistically define the methylation difference between tumor and normal samples, the p values in each case was adjusted by FDR, FDR ≤ 0.05 was considered as significance cut-off.

Methylation data of each patient and the available clinical overall survival data was combined to generate methylation-specific survivality analysis. Gene methylation level was classified into 2 groups by median methylation. Cox regression analysis was performed to estimate the hazards ratio. Condition: If Cox coef > 0, the hyper (high) methylation group shows a worse survival, otherwise low methylation with a better survivability. Log rank P test was performed to evaluate the distribution comparison of two groups where, p value <0.05 was considered significant.

The mRNA expression and methylation data for each available sample with FERMT1, 2, and 3 gene methylation were merged using sample specific TCGA barcode. Person’s product moment correlation coefficient method was used to test the association between paired mRNA expression and methylation. P-value was calculated by adjusting corresponding FDR and FDR ≤ 0.05 was considered.

##### Classification of stabilizing and destabilizing mutants

We calculated the stability of the mutants with respect to their respective wild-type structures using their change in free energy change. ΔΔG values of all the mutants were taken together and each data were repeated similar times to the frequency of the corresponding mutation found in all types of cancer. Separate subsets were prepared for stabilizing mutants (**ΔΔG (+ve)**) and destabilizing mutants ((**ΔΔG (-ve)**).

We classified this overall dataset by ranking the mutations in percentile according to their ΔΔG values.

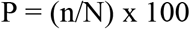

n = number of samples below a particular value; N = total number of samples; P = percentile of that particular value

In stabilizing data set 0-25 percentile region (P1) was considered as low, 25-75 percentile (P2) and 75-100 percentile (P3) region were considered as moderate and high respectively. In destabilizing data set 0-25 percentile region (P1) was considered as high, 25-75 percentile (P2) and 75-100 percentile (P3) region were considered as moderate and low respectively. As, in statistics median value is considered to be the divider between higher and lower half of a dataset, we calculated the median for P1 region for stabilizing and P3 region for destabilizing group and averaged their magnitude. Similarly, the median for P3 region for stabilizing and P1 region for destabilizing group were also calculated and their magnitude were averaged.

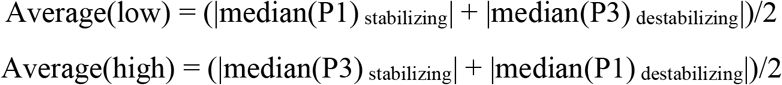

Mutants having ΔΔG values < Average(low) are considered as slightly stabilizing and ΔΔG values > -Average(low) were considered slightly destabilizing. Similarly, mutants having ΔΔG values > Average(high) are considered as very high stabilizing and ΔΔG values < - Average(high) were considered highly destabilizing. (Suppl. Method Fig. 1). Hence, both stabilizing and destabilizing single nucleotide mutants were classified into five categories-Very high, high, moderate, low and very low. We considered only very highly stabilizing and destabilizing mutations for further downstream analysis.

**Suppl. Method Fig. 1:**
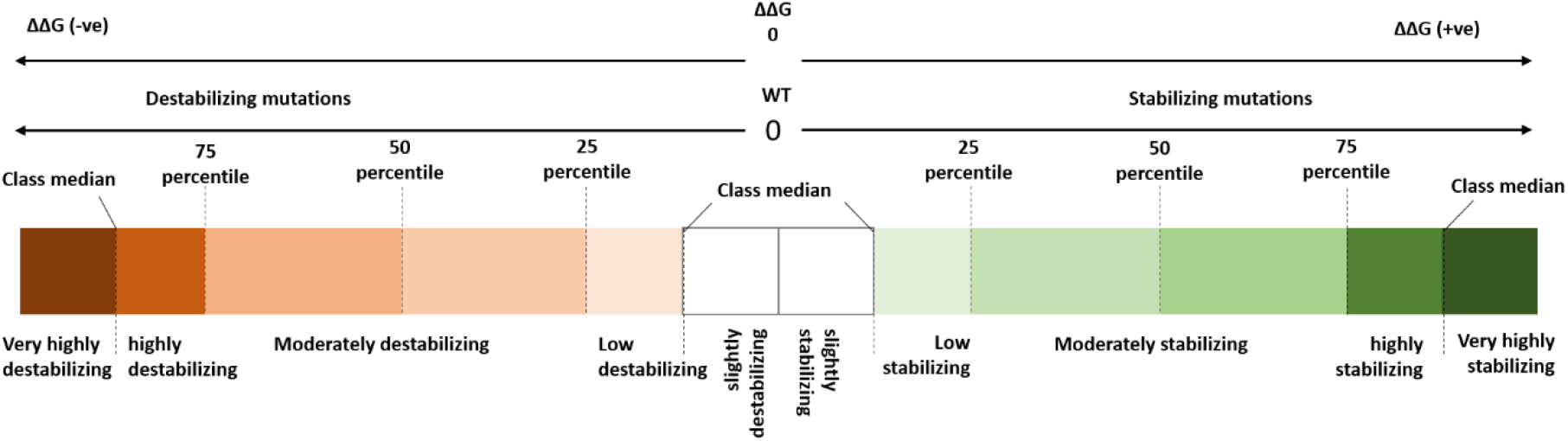
Classification of Kindlin mutations according to their stabilizing or destabilizing effect. The classifications are based on presently available data. The values (cut-offs) of percentiles and class medians are as follow:

| Status: | 0<sup>th</sup> percentile | 25<sup>th</sup> percentile | 50<sup>th</sup> percentile | 75<sup>th</sup> percentile |
| --- | --- | --- | --- | --- |
| **Stabilizing** | 0 | 0.24 | 0.52 | 0.89 |
| **Destabilizing** | 0 | -0.774 | -0.416 | -0.189 |
| .... |  |  |  |  |
| Class median | Stabilizing (High) | Stabilizing (Low) | Destabilizing (High) | Destabilizing (Low) |
|  | 1.23 | 0.12 | -1.23 | -0.12 |

##### Calculation of the signal transmission property

Assuming that one residue of a protein is perturbed based on a specific signal, it will pass-on the signal to another residue. The first passage time in Markov’s method, which is the average time for residue/node *i* to transfer the message from node *i* to *j* for the first time, is defined as hitting time *H*(*j*,*i*). If we assume *v_i_* and *v_j_* as the signal transducer (first perturbation site upon signal hitting) and receiver (response site) respectively, the passage from *v_i_* to *v_j_* can happen through an intermediate residue *v_k_* i.e., *v_i_* → *v_k_→ v_j_*.

The signal transmission property within a protein can be calculated by statistical-mechanical method based on the formula. Here, the hitting time is defined as-

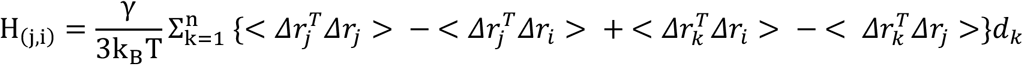

Where,

γ = spring constant

k_B_= Boltzmann constant

3k_B_T = Average total energy due to vibrational mode of a residue

(<*Δr_j_^T^Δr_j_*>) = Root mean square fluctuation of the response site.

(-<*Δr_j_^T^Δr_i_*>) = cross-correlation between the perturbation site and response site.

(<*Δr_k_^T^Δr_i_*> - <*Δr_k_^T^Δr_j_*>) = cross-correlations between intermediate residue *v_k_* and residues *v_i_* and *v_j_*.

*d_k_ = normalizing factor*

The efficiency of signal receiving or signal communication can be calculated as signal rate. It is defined as,

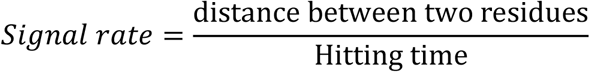

As validated by previous experiments, a low hitting time of a residue suggests its sensitivity to receive the signal. Therefore, signal rate being inversely proportional to hitting time, increases with increasing signal receiving efficiency of a residue.

##### Meta-analysis of Kindlin associated Mechanochemical signaling

We performed systematic electronic search on databases that contain research papers using the terms Mechanochemical signaling (Google Scholar, n=17800; PubMed, n=417; n, number of articles), mechanosensitive transcription factors (Google Scholar, n=31400; PubMed, n=325; n, number of articles), mechanosensitive receptors (Google Scholar, n=60700; PubMed, n=1443; n, number of articles) and mechanochemical ion channels (Google Scholar, n=49500; PubMed, n=2241; n, number of articles). Data was curated by considering the articles repeatedly coming as results, only once. Articles only containing proteins to be associated with mechano-chemical/mechanosensitive signals were considered for further analysis. The included study references were cross-searched for additional studies. Proteins with repeated mentions were considered only once. Abstracts found to be relevant to the topic of interest were shortlisted. The proteins were searched for association with kindlins using the extended interactome network of FERMT1, FERMT2 and FERMT3 obtained from Bio Grid We divided the mechanochemical proteins in two categories: Level-1 (directly associated with kindlins) and Level-2 (associated with kindlin interactors). This final list involves 20 mechanochemical transcription factors, 4 mechanochemical ion channels, 6 mechanosensitive receptor proteins, 13 mechanosensitive-cytoskeletal proteins and 14 proteins with other types of functions.

Major cellular processes where these genes are involved, were taken from GeneCards (https://www.genecards.org/). The data were extracted by all the authors individually and independently into a predefined form to enlist the type of protein, major cellular function and associated kindlin (Table S10).

**Suppl. Method Fig. 2:**
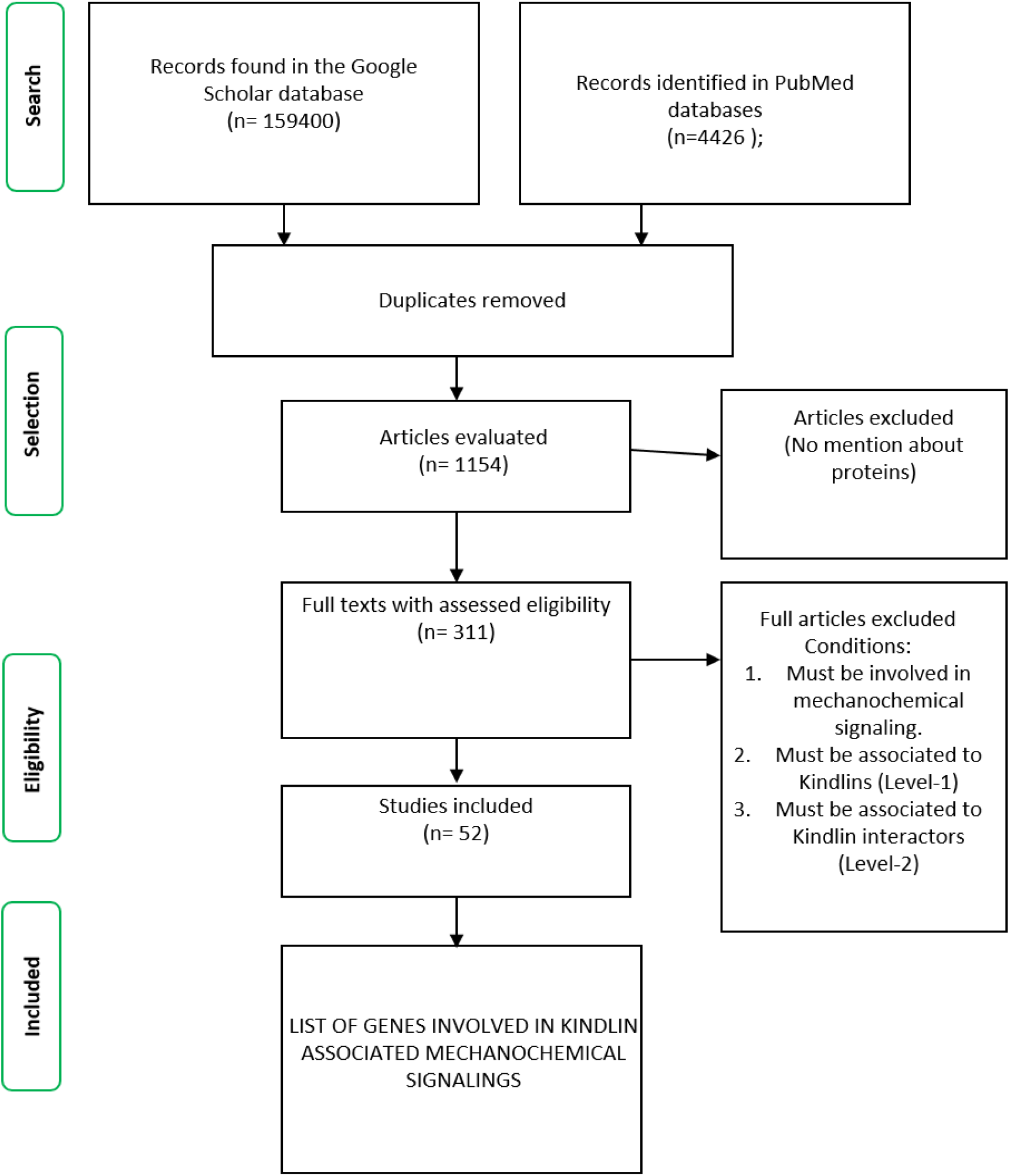
Methodological pipe-line for Meta-analysis of Kindlin associated mechanochemical signalings.

##### Cancer-specific pathway alteration analysis

We used Reverse phase protein array (RPPA) data obtained from TCPA cohort are used to calculate score for cancer samples across 32 cancer types. All of these TCPA RPPA data are from samples included in TCGA cohort. We analysed data for 10 mostly perturbed cancer related pathways including Apoptosis pathways, Cell Cycle progression, and DNA Damage Response, EMT, Hormone ER, Hormone AR, TSC/mTOR, RTK, RAS/MAPK, and PI3K/AKT pathway.

Replicates-based normalized (RBN) RPPA data were prepared as median-centered and were further normalized by standard deviation across all samples for each component. This help obtaining the relative protein level to compare pathway alteration. The pathway score is then the sum of the relative protein level of all positive regulatory components minus that of negative regulatory components in a particular pathway^18^.

We followed the same protocol followed by Ye, et al^19^. Gene expression was divided into 2 groups (High and Low) by median expression, the difference of pathway activity score (PAS) between groups is defined by student T-test whereas, p value was adjusted by FDR (cut-off FDR<=0.05). Assuming for a particular gene, Gene X, when PAS X(High) > PAS X(Low), gene X may render an activating effect to a pathway, otherwise it is considered to show an inhibiting effect to that pathway.

### Supplementary Note 2

#### Sources of Interactor-Specific GeneCards Data

##### FERMT1

*TTC37:* https://www.genecards.org/cgi-bin/carddisp.pl?gene=SKIC3

*SKIV2L:* https://www.genecards.org/cgi-bin/carddisp.pl?gene=SKIC2

*PARVA:* https://www.genecards.org/cgi-bin/carddisp.pl?gene=PARVA&keywords=PARVA

*ITGB1:* https://www.genecards.org/cgi-bin/carddisp.pl?gene=ITGB1&keywords=ITGB1

*ILK:* https://www.genecards.org/cgi-bin/carddisp.pl?gene=ILK&keywords=ILK

*FERMT3:* https://www.genecards.org/cgi-bin/carddisp.pl?gene=FERMT3&keywords=FERMT3

*FERMT2:* https://www.genecards.org/cgi-bin/carddisp.pl?gene=FERMT2&keywords=FERMT2

##### FERMT2

*VCL:* https://www.genecards.org/cgi-bin/carddisp.pl?gene=VCL&keywords=VCL

*TGFB1:* https://www.genecards.org/cgi-bin/carddisp.pl?gene=TGFB1&keywords=TGFB1

*SMAD3:* https://www.genecards.org/cgi-bin/carddisp.pl?gene=SMAD3&keywords=SMAD3

*SEPTIN9:* https://www.genecards.org/cgi-bin/carddisp.pl?gene=SEPTIN9&keywords=SEPTIN9

*SEPTIN11:* https://www.genecards.org/cgi-bin/carddisp.pl?gene=SEPTIN11&keywords=SEPTIN11

*PXN:* https://www.genecards.org/cgi-bin/carddisp.pl?gene=PXN&keywords=PXN

*PICALM:* https://www.genecards.org/cgi-bin/carddisp.pl?gene=PICALM&keywords=PICALM

*PFKM:* https://www.genecards.org/cgi-bin/carddisp.pl?gene=PFKM&keywords=PFKM

*PARVA:* https://www.genecards.org/cgi-bin/carddisp.pl?gene=PARVA&keywords=PARVA

*MAPK1:* https://www.genecards.org/cgi-bin/carddisp.pl?gene=MAPK1&keywords=MAPK1

*LIMS1:* https://www.genecards.org/cgi-bin/carddisp.pl?gene=LIMS1&keywords=LIMS1

*ITGB1:* https://www.genecards.org/cgi-bin/carddisp.pl?gene=ITGB1&keywords=ITGB1

*ILK:* https://www.genecards.org/cgi-bin/carddisp.pl?gene=ILK&keywords=ILK

*FERMT3:* https://www.genecards.org/cgi-bin/carddisp.pl?gene=FERMT3&keywords=FERMT3

*FERMT1:* https://www.genecards.org/cgi-bin/carddisp.pl?gene=FERMT1&keywords=FERMT1

##### FERMT3

*PARVB:* https://www.genecards.org/cgi-bin/carddisp.pl?gene=PARVB&keywords=PARVB

*PARVA:* https://www.genecards.org/cgi-bin/carddisp.pl?gene=PARVA&keywords=PARVA

*LSM8:* https://www.genecards.org/cgi-bin/carddisp.pl?gene=LSM8&keywords=LSM8

*LIMS1:* https://www.genecards.org/cgi-bin/carddisp.pl?gene=LIMS1&keywords=LIMS1

*ILK:* https://www.genecards.org/cgi-bin/carddisp.pl?gene=ILK&keywords=ILK

*FERMT2:* https://www.genecards.org/cgi-bin/carddisp.pl?gene=FERMT2&keywords=FERMT2

*FERMT1:* https://www.genecards.org/cgi-bin/carddisp.pl?gene=FERMT1&keywords=FERMT1

*EXOSC10:* https://www.genecards.org/cgi-bin/carddisp.pl?gene=EXOSC10&keywords=EXOSC10

**Table S1:** List of microRNAs associated with FERMT1 protein expression regulation in cancers, and their expression profile. Up, overexpressed; down, under-expressed.

| **MiRNA** | **Cancer** | **Profile** |
| --- | --- | --- |
| **hsa-miR-186-5p** | Colon | Up |
|  | Osteosarcoma | Down |
| **hsa-miR-493-3p** | Prostate | Down |
| **hsa-miR-16-1-3p** | Chronic lymphocytic leukaemia | Down |
|  | Gastric | Down |
| **hsa-miR-5692a** | Hepatocellular | Up |
| **hsa-miR-9-5p** | Colorectal | Down |
|  | Non-small cell lung | Up |
|  | Pancreatic | Down |
|  | Prostate | Up |
| **hsa-miR-660-5p** | Breast | Up |
|  | Non-small cell lung | Up |
| **hsa-miR-567** | Breast | Down |
|  | Osteosarcoma | Down |
| **hsa-miR-4723-5p** | Prostate | Down |
| **hsa-miR-577** | Colorectal | Down |
|  | Gastric | Down |
|  | Gastric | Up |
|  | Glioblastoma | Down |
|  | Hepatocellular carcinoma | Down |
|  | Non-small cell lung | Down |
|  | Papillary thyroid carcinoma | Down |
| **hsa-miR-490-5p** | Bladder | Down |
|  | Hepatocellular carcinoma | Down |
|  | Renal cell carcinoma | Down |
| **hsa-miR-539-5p** | Glioma | Down |

**Table S2:**
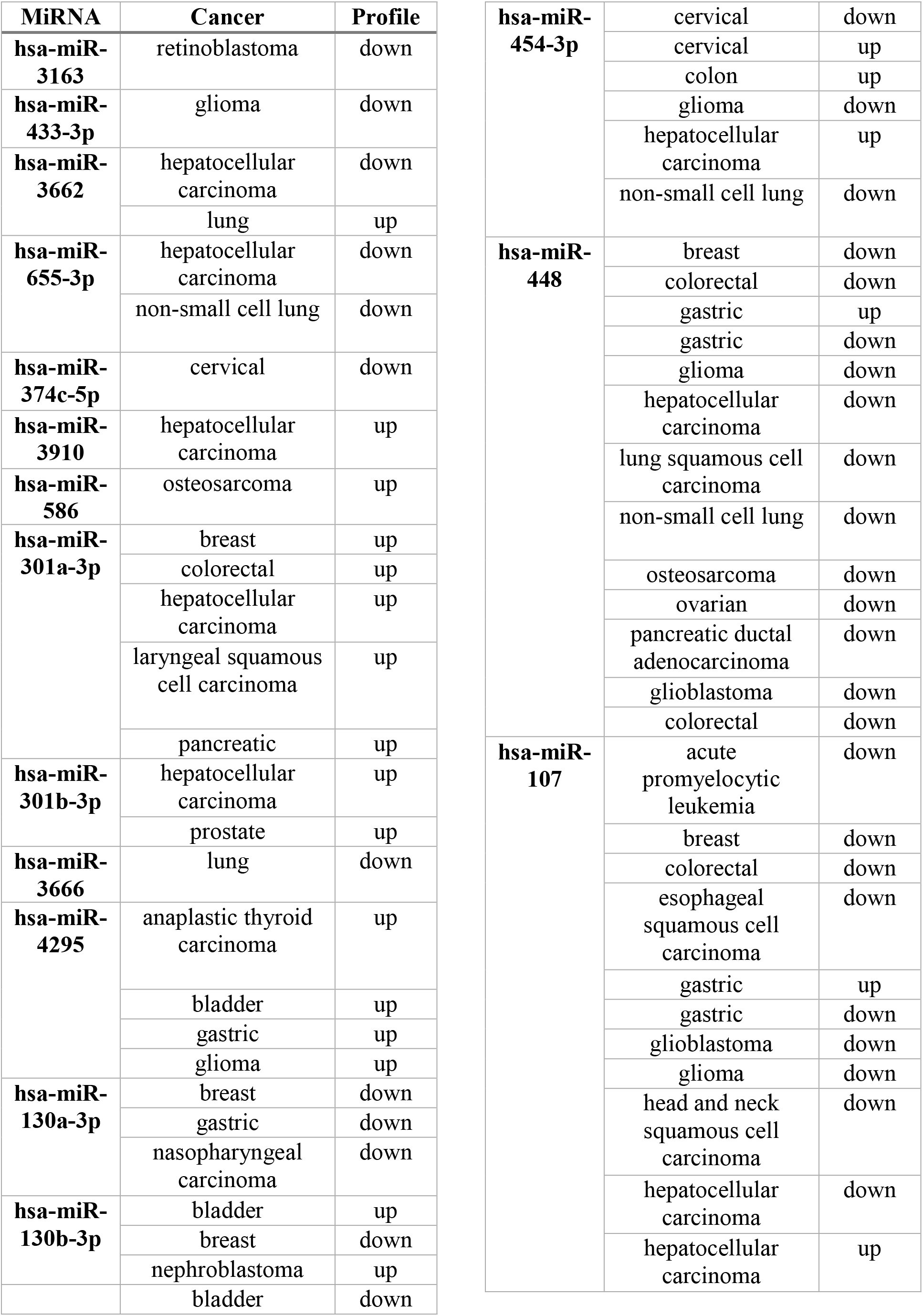
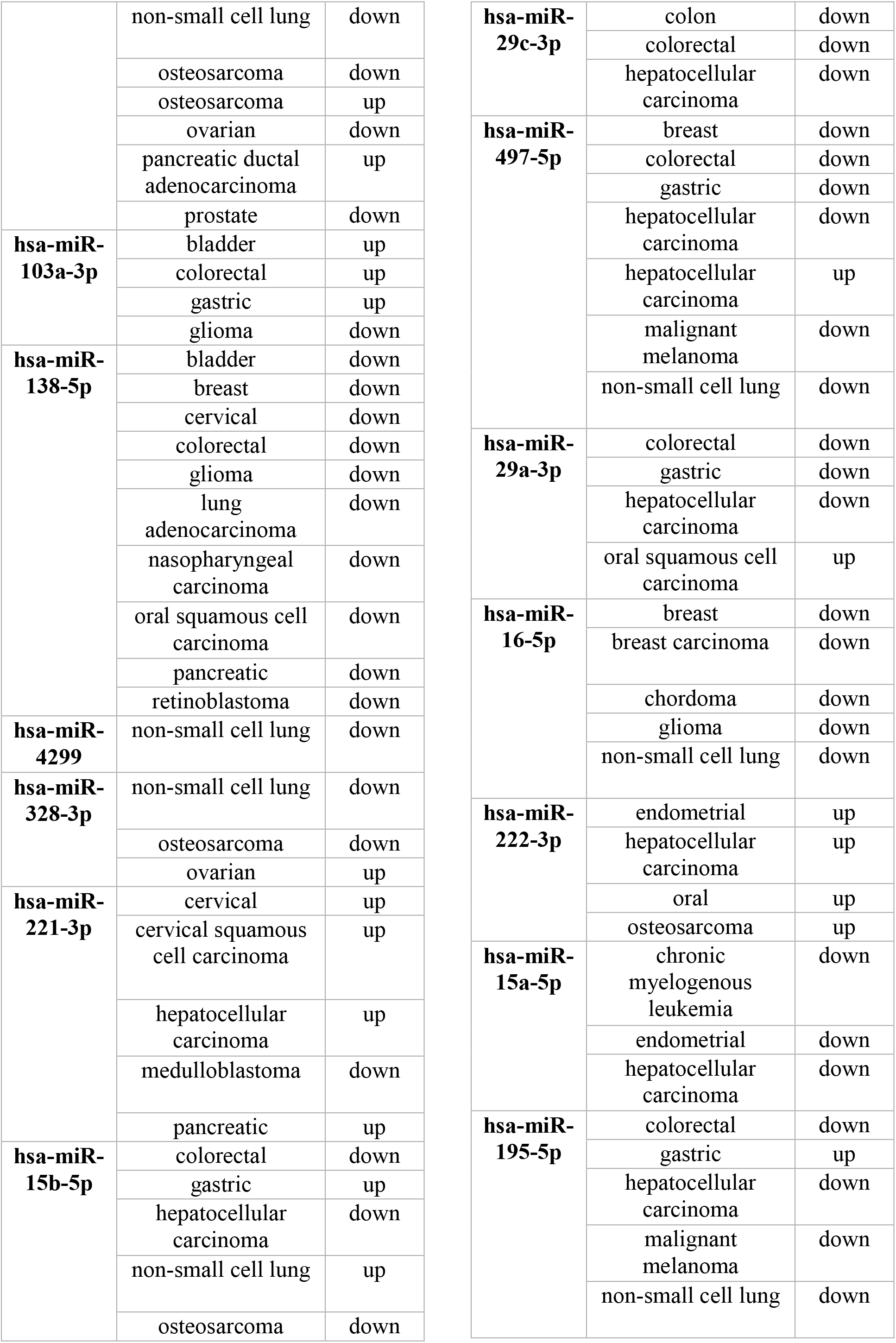
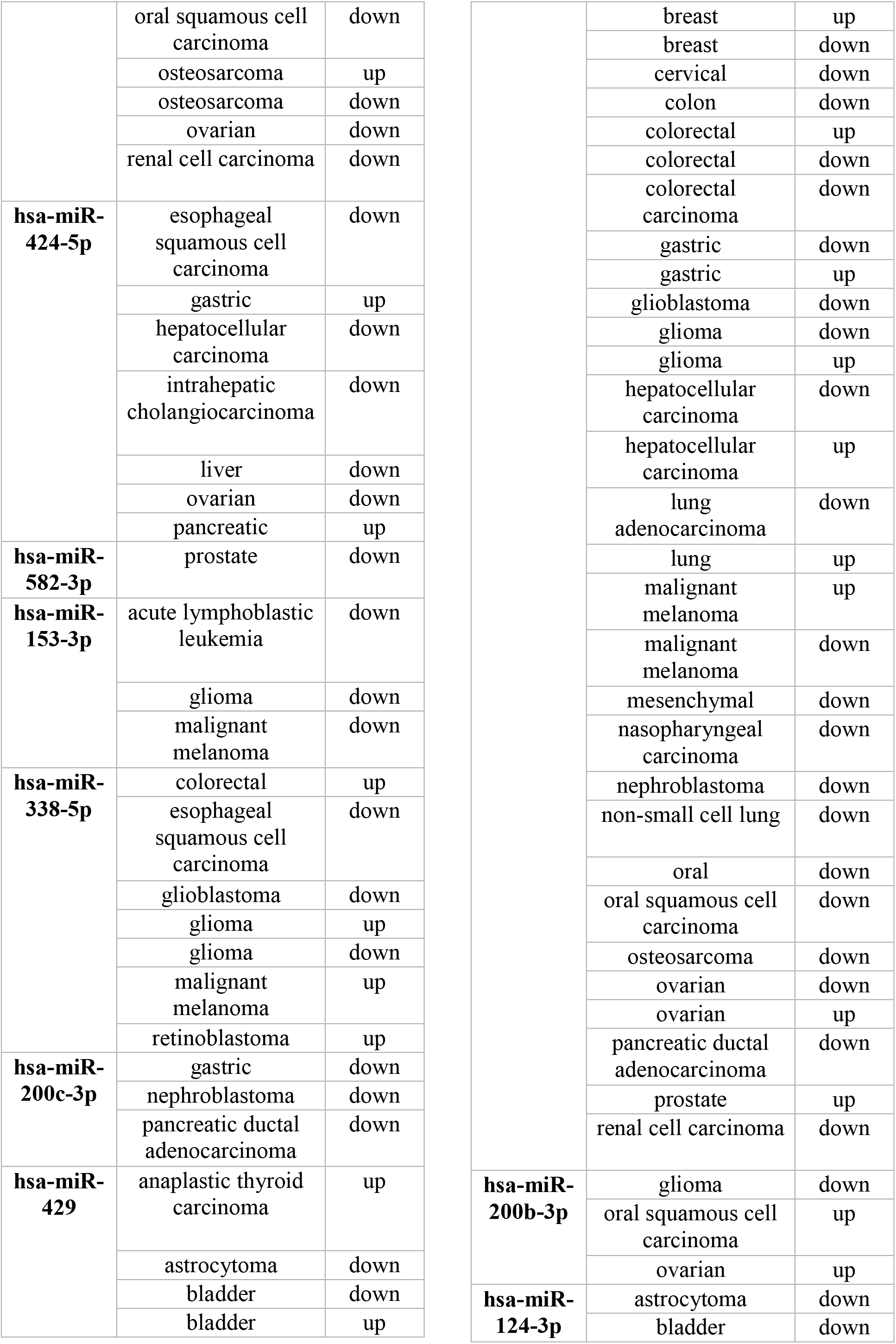
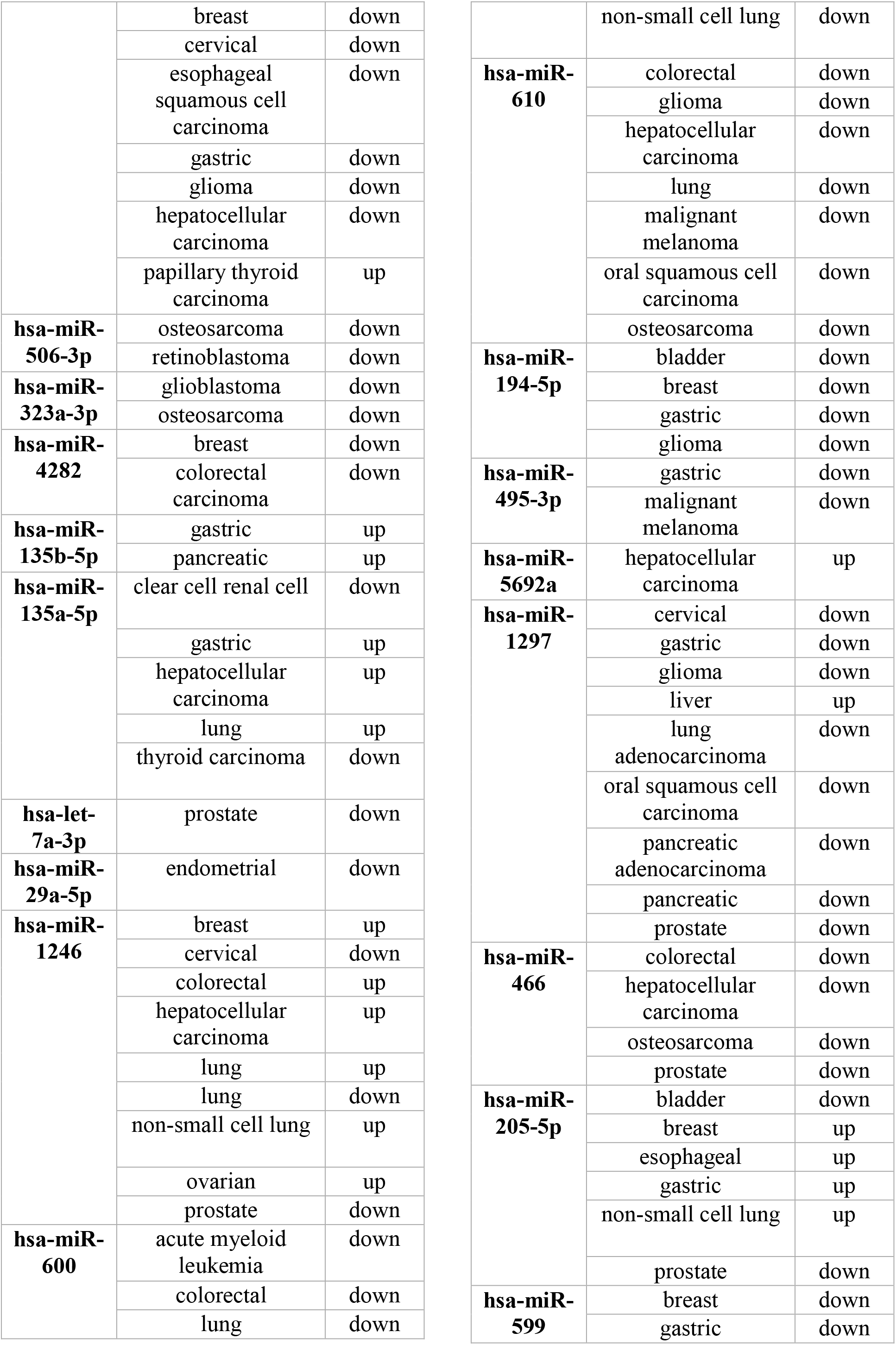
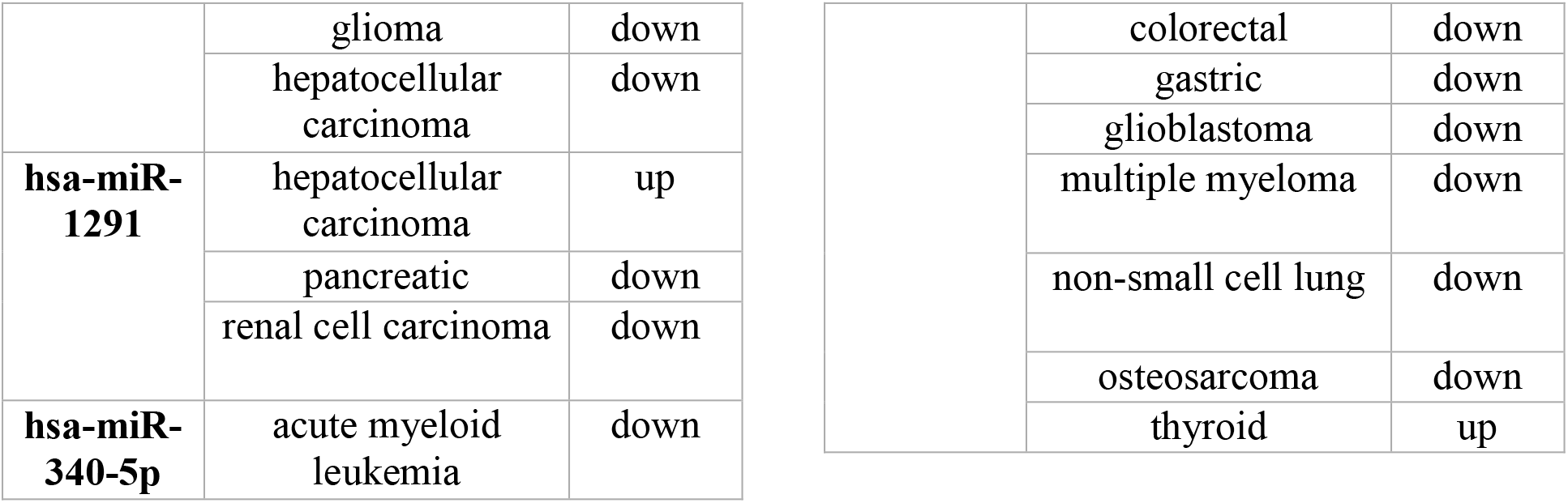
List of microRNAs associated with FERMT2 protein expression regulation in cancers, and their expression profile. Up, overexpressed; down, under-expressed.

**Table S3:** List of microRNAs associated with FERMT3 protein expression regulation in cancers, and their expression profile. Up, overexpressed; down, under-expressed.

| **MiRN A** | **Cancer** | **Profile** |
| --- | --- | --- |
| **hsa-miR-3163** | retinoblastoma | down |
| **hsa-miR-802** | breast | down |
|  | cervical | down |
|  | gastric | down |
|  | osteosarcoma | up |
|  | prostate | down |
| **hsa-miR-4775** | colorectal | up |
| **hsa-miR-542-3p** | astrocytoma | down |
|  | bladder | down |
|  | colorectal | down |
|  | gastric | down |
|  | glioblastoma | down |
|  | hepatocellular carcinoma | down |
|  | malignant melanoma | down |
|  | non-small cell lung | down |
|  | oral squamous cell carcinoma | up |
|  | oral squamous cell carcinoma | down |
|  | osteosarcoma | up |
|  | osteosarcoma | down |
| **hsa-miR-590-3p** | breast | down |
|  | colon | up |
|  | glioblastoma | down |
|  | glioblastoma | up |
|  | glioma | down |
|  | hepatocellular carcinoma | down |
|  | osteosarcoma | down |
|  | ovarian | up |
|  | prostate | up |
| **hsa-miR-509-5p** | non-small cell lung | down |
|  | pancreatic | down |
|  | papillary thyroid carcinoma | up |
|  | prostate | down |
|  | renal cell carcinoma | down |
| **hsa-miR-559** | gastric | down |
| **hsa-miR-548c-5p** | colorectal | down |
| **hsa-miR-548a-5p** | hepatocellular carcinoma | up |
| **hsa-miR-4455** | gastric | down |
| **hsa-miR-661** | glioma | down |
|  | non-small cell lung | up |
|  | ovarian | up |
|  | pancreatic ductal adenocarcinoma | up |

**Table S4:** Classification of Cancer-Associated Somatic Mutations in FERMT1 based on their impact on Structural Stability.

| **Mutation** | **Score** | **Classification** |
| --- | --- | --- |
| R17S | -1.779 | Very High Destabilizing |
| L31M | 0.455 | Moderate Stabilizing |
| L31Q | -1.567 | Very High Destabilizing |
| H38N | 0.113 | Slightly Stabilizing |
| G41E | 0.479 | Moderate Stabilizing |
| G41A | 0.381 | Moderate Stabilizing |
| M43V | -0.418 | Moderate Destabilizing |
| M43R | 0.712 | Moderate Stabilizing |
| N51K | 0.067 | Slightly Stabilizing |
| N51T | -0.337 | Moderate Destabilizing |
| S53F | 0.112 | Slightly Stabilizing |
| F59L | -0.134 | Low Destabilizing |
| F59Y | -0.21 | Moderate Destabilizing |
| A60T | 0.858 | Moderate Stabilizing |
| A60V | 0.999 | High Stabilizing |
| L61I | -0.11 | Slightly Destabilizing |
| L61F | -1.195 | High Destabilizing |
| W63L | -1.805 | Very High Destabilizing |
| W63C | -1.465 | Very High Destabilizing |
| G81A | -0.152 | Low Destabilizing |
| L88F | 0.175 | Low Stabilizing |
| R98H | -0.384 | Moderate Destabilizing |
| R98C | -0.253 | Moderate Destabilizing |
| S137F | 0.973 | High Stabilizing |
| Y145H | 0.116 | Slightly Stabilizing |
| K149H | 0.95 | High Stabilizing |
| I160T | 0.249 | Moderate Stabilizing |
| S169C | 0.476 | Moderate Stabilizing |
| S170Y | 0.781 | Moderate Stabilizing |
| P193L | 0.435 | Moderate Stabilizing |
| P193H | 0.52 | Moderate Stabilizing |
| G196E | 1.078 | High Stabilizing |
| A232V | 0.586 | Moderate Stabilizing |
| R238Q | 0.064 | Slightly Stabilizing |
| R255C | -0.161 | Low Destabilizing |
| R255H | 0.455 | Moderate Stabilizing |
| E264Q | -0.743 | Moderate Destabilizing |
| R271Q | -0.506 | Moderate Destabilizing |
| S276F | 0.62 | Moderate Stabilizing |
| S276Y | 1.162 | High Stabilizing |
| F277I | -0.37 | Moderate Destabilizing |
| D279N | -0.555 | Moderate Destabilizing |
| Y284H | 0.082 | Slightly Stabilizing |
| D285V | -0.605 | Moderate Destabilizing |
| A286V | 0.068 | Slightly Stabilizing |
| A286T | -0.494 | Moderate Destabilizing |
| R288Q | -1.024 | High Destabilizing |
| E304K | -0.232 | Moderate Destabilizing |
| Y320C | -1.6 | Very High Destabilizing |
| Y320H | -1.335 | Very High Destabilizing |
| E342K | 0.499 | Moderate Stabilizing |
| A345V | 0.255 | Moderate Stabilizing |
| A346V | -0.179 | Low Destabilizing |
| A359E | -0.061 | Slightly Destabilizing |
| L376R | -1.23 | Very High Destabilizing |
| L378V | 0.665 | Moderate Stabilizing |
| A402V | 1.894 | Very High Stabilizing |
| P427L | 0.767 | Moderate Stabilizing |
| M448I | -0.381 | Moderate Destabilizing |
| A467V | 0.473 | Moderate Stabilizing |
| A467K | -0.113 | Slightly Destabilizing |
| M495I | -0.811 | High Destabilizing |
| E509K | 0.132 | Low Stabilizing |
| D512Y | 0.214 | Low Stabilizing |
| M513T | -1.538 | Very High Destabilizing |
| R522W | -0.498 | Moderate Destabilizing |
| R522Q | -0.001 | Slightly Destabilizing |
| K525E | -0.223 | Moderate Destabilizing |
| R526K | -0.17 | Low Destabilizing |
| R535W | 0.469 | Moderate Stabilizing |
| R535P | -0.378 | Moderate Destabilizing |
| A558V | 1.025 | High Stabilizing |
| E564K | 0.708 | Moderate Stabilizing |
| R573I | -0.469 | Moderate Destabilizing |
| R605K | -0.613 | Moderate Destabilizing |
| I609V | 0.446 | Moderate Stabilizing |
| R619W | 0.612 | Moderate Stabilizing |
| H644R | -0.263 | Moderate Destabilizing |
| E645K | -1.549 | Very High Destabilizing |
| G649S | -0.89 | High Destabilizing |
| G674V | 1.188 | High Stabilizing |
| S4F | 0.363 | Moderate Stabilizing |
| T5A | -0.197 | Moderate Destabilizing |
| V18I | -0.165 | Low Destabilizing |
| D19G | -0.561 | Moderate Destabilizing |
| E23K | 0.136 | Low Stabilizing |
| Q25K | -0.542 | Moderate Destabilizing |
| Q26K | -0.1 | Slightly Destabilizing |
| V29I | 0.014 | Slightly Stabilizing |
| T30S | -0.277 | Moderate Destabilizing |
| R32K | -0.219 | Moderate Destabilizing |
| I52V | -0.202 | Moderate Destabilizing |
| D55H | -0.014 | Slightly Destabilizing |
| W75C | -0.822 | High Destabilizing |
| V82I | 0.858 | Moderate Stabilizing |
| T91I | 1.655 | Very High Stabilizing |
| P92R | 1.916 | Very High Stabilizing |
| M96I | -0.753 | Moderate Destabilizing |
| S112R | 0.288 | Moderate Stabilizing |
| S114A | -0.032 | Slightly Destabilizing |
| F118V | -0.646 | Moderate Destabilizing |
| I124T | -2.764 | Very High Destabilizing |
| R131T | -1.008 | High Destabilizing |
| S133L | 1.392 | Very High Stabilizing |
| E135D | -0.98 | High Destabilizing |
| P141S | -1.118 | High Destabilizing |
| F146Y | 0.285 | Moderate Stabilizing |
| K148N | 0.257 | Moderate Stabilizing |
| K154Q | -0.14 | Low Destabilizing |
| N156D | -0.189 | Low Destabilizing |
| P159L | 1.247 | Very High Stabilizing |
| I161N | 0.297 | Moderate Stabilizing |
| T172A | 0.051 | Slightly Stabilizing |
| A173T | -0.437 | Moderate Destabilizing |
| P180L | 1.066 | High Stabilizing |
| T186I | -0.041 | Slightly Destabilizing |
| M187I | 0.36 | Moderate Stabilizing |
| S201P | 0.76 | Moderate Stabilizing |
| P210S | 1.183 | High Stabilizing |
| T212M | 0.52 | Moderate Stabilizing |
| Q226K | 0.517 | Moderate Stabilizing |
| L231F | 0.05 | Slightly Stabilizing |
| Y235S | -1.522 | Very High Destabilizing |
| A244S | -1.534 | Very High Destabilizing |
| G249C | 1.245 | Very High Stabilizing |
| D252Y | 1.29 | Very High Stabilizing |
| S253P | 0.538 | Moderate Stabilizing |
| L257I | -0.05 | Slightly Destabilizing |
| Q263R | 0.178 | Low Stabilizing |
| P282L | 0.716 | Moderate Stabilizing |
| Q291E | 0.057 | Slightly Stabilizing |
| I300V | -1.585 | Very High Destabilizing |
| D306Y | 1.361 | Very High Stabilizing |
| T308R | 0.575 | Moderate Stabilizing |
| E309K | 0.374 | Moderate Stabilizing |
| E310K | 1.16 | High Stabilizing |
| E311K | 0.66 | Moderate Stabilizing |
| H321Y | 0.654 | Moderate Stabilizing |
| S328F | 0.894 | High Stabilizing |
| D333V | 0.706 | Moderate Stabilizing |
| E339K | 0.332 | Moderate Stabilizing |
| D341N | 0.252 | Moderate Stabilizing |
| G356D | 0.668 | Moderate Stabilizing |
| G357E | 0.282 | Moderate Stabilizing |
| L384V | 0.006 | Slightly Stabilizing |
| P386L | -0.307 | Moderate Destabilizing |
| F394C | -0.67 | Moderate Destabilizing |
| S400F | 0.649 | Moderate Stabilizing |
| L420I | -0.384 | Moderate Destabilizing |
| E424K | -0.346 | Moderate Destabilizing |
| D428N | -0.488 | Moderate Destabilizing |
| V431L | 0.632 | Moderate Stabilizing |
| A432V | 0.892 | High Stabilizing |
| P443S | -0.584 | Moderate Destabilizing |
| V444A | 0.213 | Low Stabilizing |
| N449T | 0.118 | Slightly Stabilizing |
| E450K | 0.265 | Moderate Stabilizing |
| K475R | 0.793 | Moderate Stabilizing |
| M477I | 0.256 | Moderate Stabilizing |
| N488S | 0.075 | Slightly Stabilizing |
| S506Y | 0.068 | Slightly Stabilizing |
| M511I | -0.719 | Moderate Destabilizing |
| E516Q | -0.543 | Moderate Destabilizing |
| S520L | 1.121 | High Stabilizing |
| L532Q | -0.705 | Moderate Destabilizing |
| A533T | -1.338 | Very High Destabilizing |
| A534T | -0.528 | Moderate Destabilizing |
| A544V | 0.544 | Moderate Stabilizing |
| R554Q | -0.224 | Moderate Destabilizing |
| Q560H | 1.154 | High Stabilizing |
| S561L | 1.093 | High Stabilizing |
| V572F | -0.393 | Moderate Destabilizing |
| G576R | -1.239 | Very High Destabilizing |
| S577G | -0.563 | Moderate Destabilizing |
| S586L | 1.331 | Very High Stabilizing |
| D594N | -0.173 | Low Destabilizing |
| T597S | 0.256 | Moderate Stabilizing |
| G598R | -0.21 | Moderate Destabilizing |
| E617K | -0.049 | Slightly Destabilizing |
| E624K | -0.574 | Moderate Destabilizing |
| D639Y | 1.168 | High Stabilizing |
| Y646C | -0.979 | High Destabilizing |
| L653F | 0.295 | Moderate Stabilizing |
| S657F | 0.602 | Moderate Stabilizing |
| K658R | 0.512 | Moderate Stabilizing |
| D659N | 0.045 | Slightly Stabilizing |
| Q660H | 0.346 | Moderate Stabilizing |
| E662V | -0.554 | Moderate Destabilizing |
| L664R | -0.004 | Slightly Destabilizing |
| D665N | 0.491 | Moderate Stabilizing |
| E666A | -0.488 | Moderate Destabilizing |
| K671R | 0.657 | Moderate Stabilizing |
| Q676H | 1.134 | High Stabilizing |

**Table S5:**
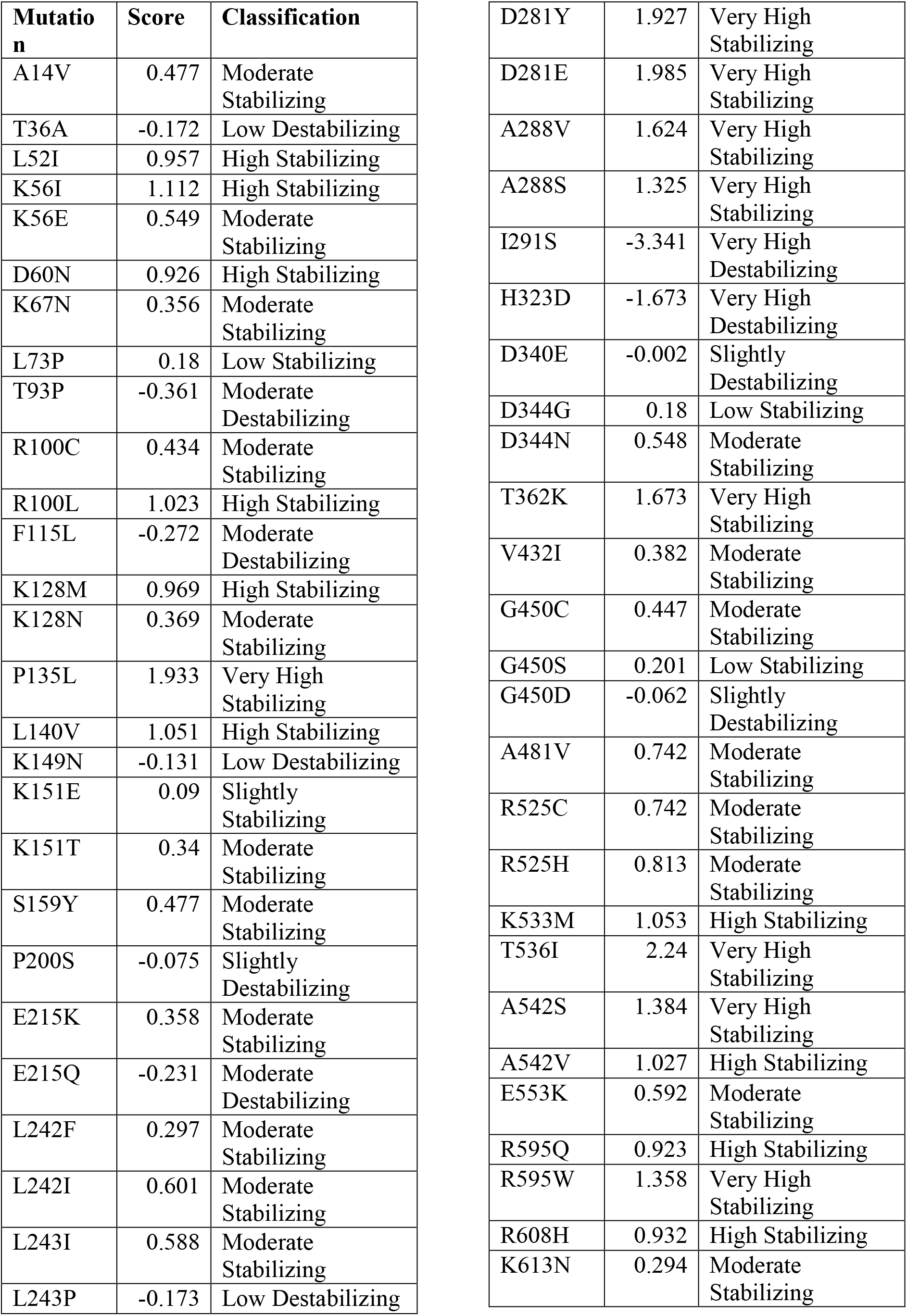
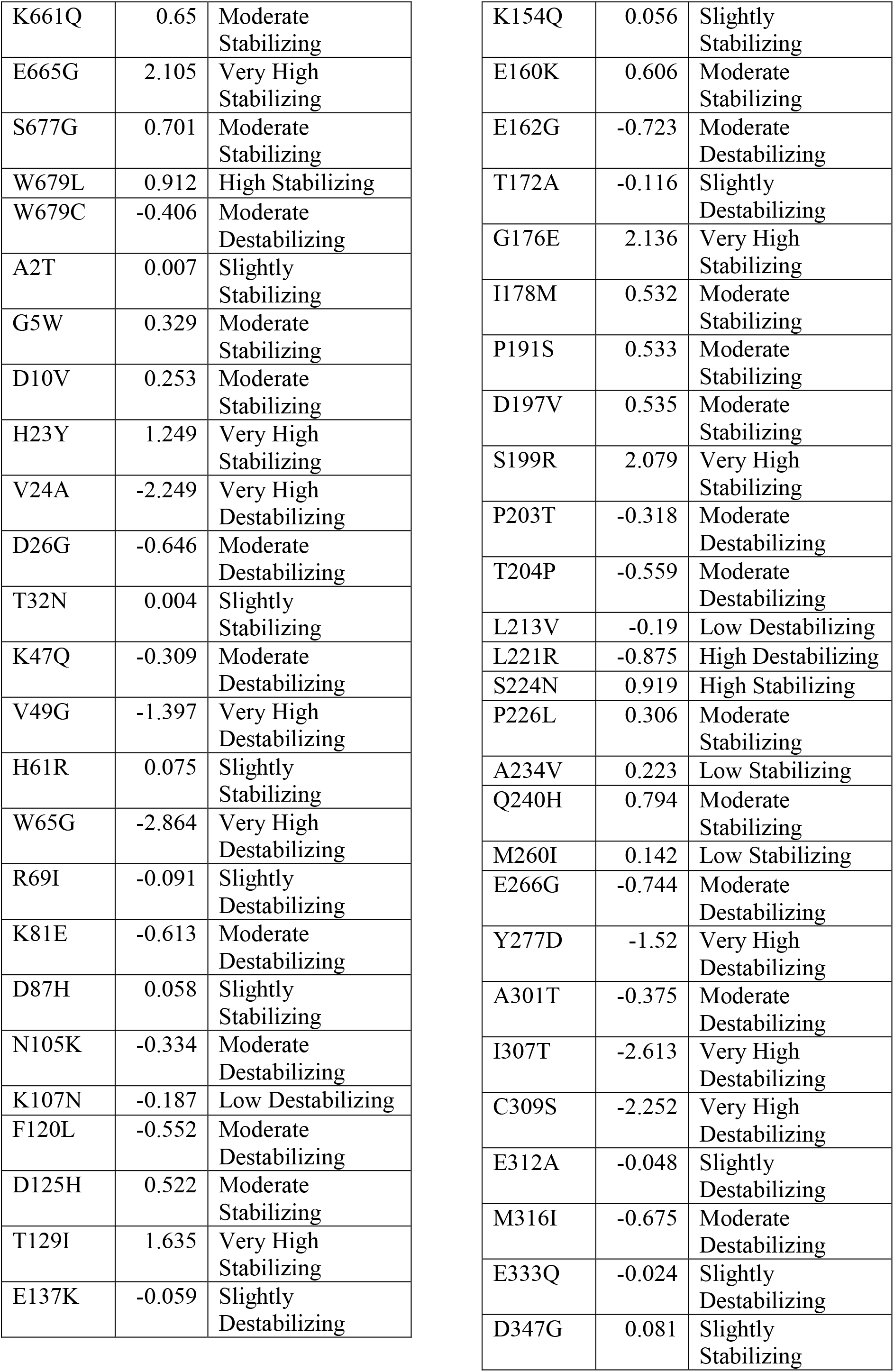
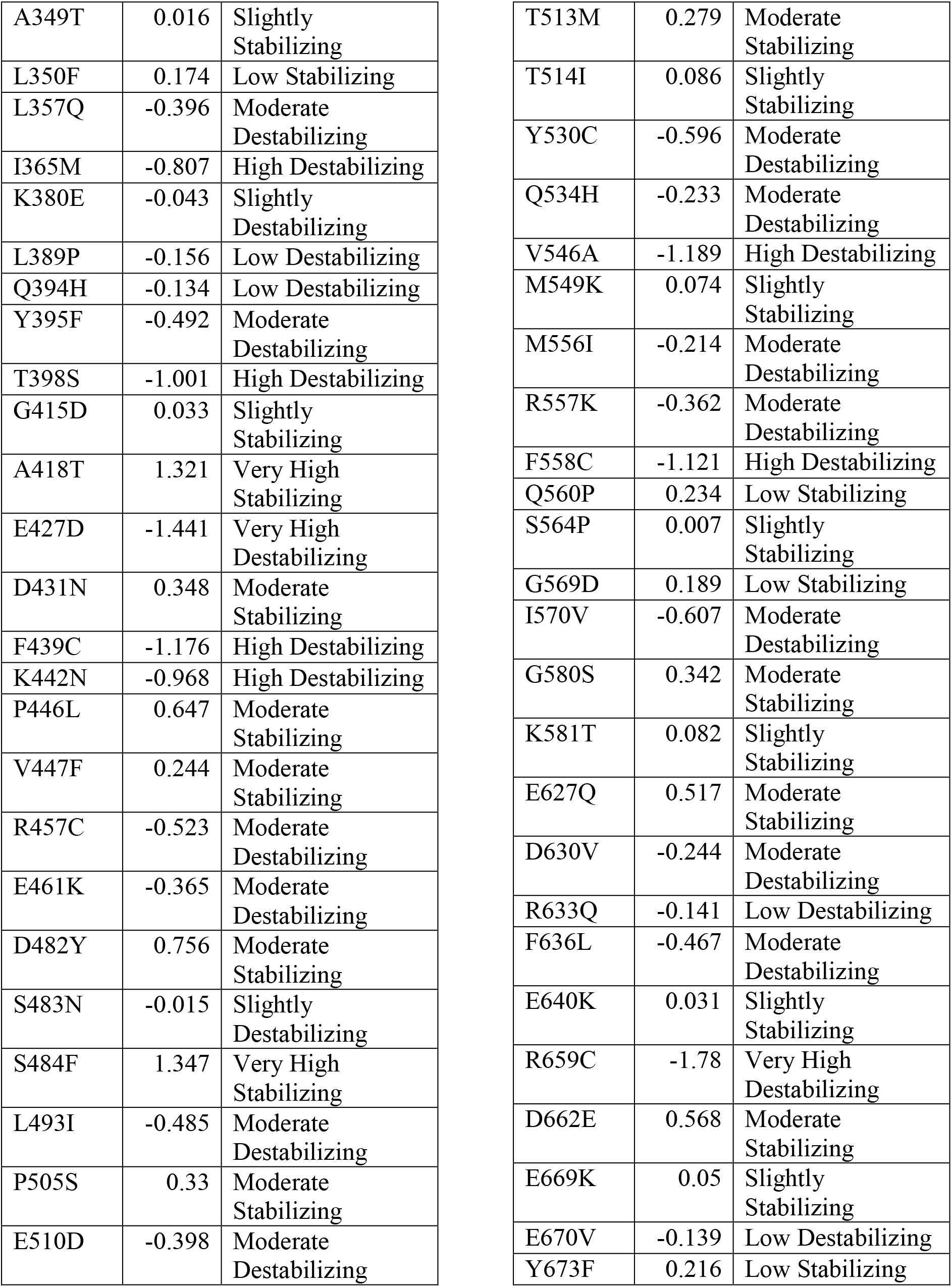
Classification of Cancer-Associated Somatic Mutations in FERMT2 based on their impact on Structural Stability.

| **Mutation** | **Score** | **Classification** |
| --- | --- | --- |
| A14V | 0.477 | Moderate Stabilizing |
| T36A | -0.172 | Low Destabilizing |
| L52I | 0.957 | High Stabilizing |
| K56I | 1.112 | High Stabilizing |
| K56E | 0.549 | Moderate Stabilizing |
| D60N | 0.926 | High Stabilizing |
| K67N | 0.356 | Moderate Stabilizing |
| L73P | 0.18 | Low Stabilizing |
| T93P | -0.361 | Moderate Destabilizing |
| R100C | 0.434 | Moderate Stabilizing |
| R100L | 1.023 | High Stabilizing |
| F115L | -0.272 | Moderate Destabilizing |
| K128M | 0.969 | High Stabilizing |
| K128N | 0.369 | Moderate Stabilizing |
| P135L | 1.933 | Very High Stabilizing |
| L140V | 1.051 | High Stabilizing |
| K149N | -0.131 | Low Destabilizing |
| K151E | 0.09 | Slightly Stabilizing |
| K151T | 0.34 | Moderate Stabilizing |
| S159Y | 0.477 | Moderate Stabilizing |
| P200S | -0.075 | Slightly Destabilizing |
| E215K | 0.358 | Moderate Stabilizing |
| E215Q | -0.231 | Moderate Destabilizing |
| L242F | 0.297 | Moderate Stabilizing |
| L242I | 0.601 | Moderate Stabilizing |
| L243I | 0.588 | Moderate Stabilizing |
| L243P | -0.173 | Low Destabilizing |
| D281Y | 1.927 | Very High Stabilizing |
| D281E | 1.985 | Very High Stabilizing |
| A288V | 1.624 | Very High Stabilizing |
| A288S | 1.325 | Very High Stabilizing |
| I291S | -3.341 | Very High Destabilizing |
| H323D | -1.673 | Very High Destabilizing |
| D340E | -0.002 | Slightly Destabilizing |
| D344G | 0.18 | Low Stabilizing |
| D344N | 0.548 | Moderate Stabilizing |
| T362K | 1.673 | Very High Stabilizing |
| V432I | 0.382 | Moderate Stabilizing |
| G450C | 0.447 | Moderate Stabilizing |
| G450S | 0.201 | Low Stabilizing |
| G450D | -0.062 | Slightly Destabilizing |
| A481V | 0.742 | Moderate Stabilizing |
| R525C | 0.742 | Moderate Stabilizing |
| R525H | 0.813 | Moderate Stabilizing |
| K533M | 1.053 | High Stabilizing |
| T536I | 2.24 | Very High Stabilizing |
| A542S | 1.384 | Very High Stabilizing |
| A542V | 1.027 | High Stabilizing |
| E553K | 0.592 | Moderate Stabilizing |
| R595Q | 0.923 | High Stabilizing |
| R595W | 1.358 | Very High Stabilizing |
| R608H | 0.932 | High Stabilizing |
| K613N | 0.294 | Moderate Stabilizing |
| K661Q | 0.65 | Moderate Stabilizing |
| E665G | 2.105 | Very High Stabilizing |
| S677G | 0.701 | Moderate Stabilizing |
| W679L | 0.912 | High Stabilizing |
| W679C | -0.406 | Moderate Destabilizing |
| A2T | 0.007 | Slightly Stabilizing |
| G5W | 0.329 | Moderate Stabilizing |
| D10V | 0.253 | Moderate Stabilizing |
| H23Y | 1.249 | Very High Stabilizing |
| V24A | -2.249 | Very High Destabilizing |
| D26G | -0.646 | Moderate Destabilizing |
| T32N | 0.004 | Slightly Stabilizing |
| K47Q | -0.309 | Moderate Destabilizing |
| V49G | -1.397 | Very High Destabilizing |
| H61R | 0.075 | Slightly Stabilizing |
| W65G | -2.864 | Very High Destabilizing |
| R69I | -0.091 | Slightly Destabilizing |
| K81E | -0.613 | Moderate Destabilizing |
| D87H | 0.058 | Slightly Stabilizing |
| N105K | -0.334 | Moderate Destabilizing |
| K107N | -0.187 | Low Destabilizing |
| F120L | -0.552 | Moderate Destabilizing |
| D125H | 0.522 | Moderate Stabilizing |
| T129I | 1.635 | Very High Stabilizing |
| E137K | -0.059 | Slightly Destabilizing |
| K154Q | 0.056 | Slightly Stabilizing |
| E160K | 0.606 | Moderate Stabilizing |
| E162G | -0.723 | Moderate Destabilizing |
| T172A | -0.116 | Slightly Destabilizing |
| G176E | 2.136 | Very High Stabilizing |
| I178M | 0.532 | Moderate Stabilizing |
| P191S | 0.533 | Moderate Stabilizing |
| D197V | 0.535 | Moderate Stabilizing |
| S199R | 2.079 | Very High Stabilizing |
| P203T | -0.318 | Moderate Destabilizing |
| T204P | -0.559 | Moderate Destabilizing |
| L213V | -0.19 | Low Destabilizing |
| L221R | -0.875 | High Destabilizing |
| S224N | 0.919 | High Stabilizing |
| P226L | 0.306 | Moderate Stabilizing |
| A234V | 0.223 | Low Stabilizing |
| Q240H | 0.794 | Moderate Stabilizing |
| M260I | 0.142 | Low Stabilizing |
| E266G | -0.744 | Moderate Destabilizing |
| Y277D | -1.52 | Very High Destabilizing |
| A301T | -0.375 | Moderate Destabilizing |
| I307T | -2.613 | Very High Destabilizing |
| C309S | -2.252 | Very High Destabilizing |
| E312A | -0.048 | Slightly Destabilizing |
| M316I | -0.675 | Moderate Destabilizing |
| E333Q | -0.024 | Slightly Destabilizing |
| D347G | 0.081 | Slightly Stabilizing |
| A349T | 0.016 | Slightly Stabilizing |
| L350F | 0.174 | Low Stabilizing |
| L357Q | -0.396 | Moderate Destabilizing |
| I365M | -0.807 | High Destabilizing |
| K380E | -0.043 | Slightly Destabilizing |
| L389P | -0.156 | Low Destabilizing |
| Q394H | -0.134 | Low Destabilizing |
| Y395F | -0.492 | Moderate Destabilizing |
| T398S | -1.001 | High Destabilizing |
| G415D | 0.033 | Slightly Stabilizing |
| A418T | 1.321 | Very High Stabilizing |
| E427D | -1.441 | Very High Destabilizing |
| D431N | 0.348 | Moderate Stabilizing |
| F439C | -1.176 | High Destabilizing |
| K442N | -0.968 | High Destabilizing |
| P446L | 0.647 | Moderate Stabilizing |
| V447F | 0.244 | Moderate Stabilizing |
| R457C | -0.523 | Moderate Destabilizing |
| E461K | -0.365 | Moderate Destabilizing |
| D482Y | 0.756 | Moderate Stabilizing |
| S483N | -0.015 | Slightly Destabilizing |
| S484F | 1.347 | Very High Stabilizing |
| L493I | -0.485 | Moderate Destabilizing |
| P505S | 0.33 | Moderate Stabilizing |
| E510D | -0.398 | Moderate Destabilizing |
| T513M | 0.279 | Moderate Stabilizing |
| T514I | 0.086 | Slightly Stabilizing |
| Y530C | -0.596 | Moderate Destabilizing |
| Q534H | -0.233 | Moderate Destabilizing |
| V546A | -1.189 | High Destabilizing |
| M549K | 0.074 | Slightly Stabilizing |
| M556I | -0.214 | Moderate Destabilizing |
| R557K | -0.362 | Moderate Destabilizing |
| F558C | -1.121 | High Destabilizing |
| Q560P | 0.234 | Low Stabilizing |
| S564P | 0.007 | Slightly Stabilizing |
| G569D | 0.189 | Low Stabilizing |
| I570V | -0.607 | Moderate Destabilizing |
| G580S | 0.342 | Moderate Stabilizing |
| K581T | 0.082 | Slightly Stabilizing |
| E627Q | 0.517 | Moderate Stabilizing |
| D630V | -0.244 | Moderate Destabilizing |
| R633Q | -0.141 | Low Destabilizing |
| F636L | -0.467 | Moderate Destabilizing |
| E640K | 0.031 | Slightly Stabilizing |
| R659C | -1.78 | Very High Destabilizing |
| D662E | 0.568 | Moderate Stabilizing |
| E669K | 0.05 | Slightly Stabilizing |
| E670V | -0.139 | Low Destabilizing |
| Y673F | 0.216 | Low Stabilizing |

**Table S6:** Classification of Cancer-Associated Somatic Mutations in FERMT3 based on their impact on Structural Stability.

| **Mutation** | **Score** | **Classification** |
| --- | --- | --- |
| S14L | 0.589 | Moderate Stabilizing |
| R19L | -0.304 | Moderate Destabilizing |
| G44E | 0.399 | Moderate Stabilizing |
| G44R | 0.66 | Moderate Stabilizing |
| K48N | -0.389 | Moderate Destabilizing |
| Q52E | -0.233 | Moderate Destabilizing |
| Q52H | 0.521 | Moderate Stabilizing |
| R55C | -0.363 | Moderate Destabilizing |
| Q75H | 0.403 | Moderate Stabilizing |
| A89T | 0.134 | Low Stabilizing |
| R90C | -1.096 | High Destabilizing |
| R90H | -0.83 | High Destabilizing |
| P95L | 0.606 | Moderate Stabilizing |
| R103W | 0.544 | Moderate Stabilizing |
| R103Q | -0.349 | Moderate Destabilizing |
| R107S | -1.283 | Very High Destabilizing |
| R107H | -0.624 | Moderate Destabilizing |
| R108H | -0.594 | Moderate Destabilizing |
| R111C | -0.525 | Moderate Destabilizing |
| R113C | -0.115 | Slightly Destabilizing |
| R113H | 0.619 | Moderate Stabilizing |
| F121L | -0.763 | Moderate Destabilizing |
| F121V | -1.349 | Very High Destabilizing |
| R129S | -0.719 | Moderate Destabilizing |
| P136L | 1.559 | Very High Stabilizing |
| E137D | 0.227 | Low Stabilizing |
| E137K | 0.308 | Moderate Stabilizing |
| A176S | -0.039 | Slightly Destabilizing |
| R179Q | 0.01 | Slightly Stabilizing |
| R179L | 0.226 | Low Stabilizing |
| S186L | 1.554 | Very High Stabilizing |
| R200Q | -0.185 | Low Destabilizing |
| Q202H | 0.521 | Moderate Stabilizing |
| D206N | -0.093 | Slightly Destabilizing |
| L210I | -0.142 | Low Destabilizing |
| R212C | -0.208 | Moderate Destabilizing |
| L219Q | 0.858 | Moderate Stabilizing |
| L225I | -0.427 | Moderate Destabilizing |
| S232L | 2.233 | Very High Stabilizing |
| S233L | 1.673 | Very High Stabilizing |
| Q239H | 0.061 | Slightly Stabilizing |
| Q239E | -0.677 | Moderate Destabilizing |
| A246P | 0.077 | Slightly Stabilizing |
| R250H | -0.774 | High Destabilizing |
| R250C | -1.032 | High Destabilizing |
| D258Y | 1.585 | Very High Stabilizing |
| R267Q | -0.358 | Moderate Destabilizing |
| R276Q | -1.631 | Very High Destabilizing |
| R276W | 0.117 | Slightly Stabilizing |
| A295V | -0.051 | Slightly Destabilizing |
| A296S | -0.906 | High Destabilizing |
| A296T | -0.599 | Moderate Destabilizing |
| S305F | 1.677 | Very High Stabilizing |
| S337L | 0.614 | Moderate Stabilizing |
| D344V | -0.169 | Low Destabilizing |
| D344E | 0.323 | Moderate Stabilizing |
| R360W | -0.468 | Moderate Destabilizing |
| R360Q | -0.446 | Moderate Destabilizing |
| R366W | 0.573 | Moderate Stabilizing |
| R366Q | 0.457 | Moderate Stabilizing |
| G372D | 0.149 | Low Stabilizing |
| D392G | 0.186 | Low Stabilizing |
| G396V | 0.611 | Moderate Stabilizing |
| G396W | 0.771 | Moderate Stabilizing |
| G396R | 0.732 | Moderate Stabilizing |
| E434K | 0.648 | Moderate Stabilizing |
| E434Q | -0.026 | Slightly Destabilizing |
| S468C | 0.083 | Slightly Stabilizing |
| S468N | 0.626 | Moderate Stabilizing |
| E469K | -0.126 | Low Destabilizing |
| L474M | -0.923 | High Destabilizing |
| R481C | -0.264 | Moderate Destabilizing |
| R481H | -0.016 | Slightly Destabilizing |
| H490Y | 0.1111 | Slightly Stabilizing |
| A498V | 0.386 | Moderate Stabilizing |
| E499K | 0.109 | Slightly Stabilizing |
| R510S | -0.859 | High Destabilizing |
| R510H | -0.649 | Moderate Destabilizing |
| R513Q | -0.101 | Slightly Destabilizing |
| R523W | -0.294 | Moderate Destabilizing |
| R523Q | -0.384 | Moderate Destabilizing |
| A527V | 0.293 | Moderate Stabilizing |
| A527S | -0.219 | Moderate Destabilizing |
| A546S | -0.317 | Moderate Destabilizing |
| E569K | 0.652 | Moderate Stabilizing |
| G572C | 0.371 | Moderate Stabilizing |
| G572V | 1.321 | Very High Stabilizing |
| R577Q | -0.688 | Moderate Destabilizing |
| R580H | -0.464 | Moderate Destabilizing |
| R580L | -0.02 | Slightly Destabilizing |
| R580C | -0.442 | Moderate Destabilizing |
| R580G | -1.599 | Very High Destabilizing |
| V588M | -0.161 | Low Destabilizing |
| V619M | -0.203 | Moderate Destabilizing |
| R629Q | -0.74 | Moderate Destabilizing |
| R644W | -0.324 | Moderate Destabilizing |
| R644Q | -0.928 | High Destabilizing |
| R644L | -0.558 | Moderate Destabilizing |
| R646Q | 0 | No Effect |
| R646W | 0.234 | Low Stabilizing |
| L658M | -0.699 | Moderate Destabilizing |
| G3E | -0.222 | Moderate Destabilizing |
| M4I | -0.472 | Moderate Destabilizing |
| T6A | -0.626 | Moderate Destabilizing |
| G9E | -0.92 | High Destabilizing |
| Y11C | -0.394 | Moderate Destabilizing |
| V20G | -3.147 | Very High Destabilizing |
| E24Q | -0.422 | Moderate Destabilizing |
| E25D | -0.636 | Moderate Destabilizing |
| D26N | 0.871 | Moderate Stabilizing |
| P27L | 0.502 | Moderate Stabilizing |
| R35W | 0.613 | Moderate Stabilizing |
| T37N | 0.174 | Low Stabilizing |
| I42V | -1.251 | Very High Destabilizing |
| V45G | -2.117 | Very High Destabilizing |
| L46F | 0.838 | Moderate Stabilizing |
| W65R | -1.531 | Very High Destabilizing |
| K69T | -0.112 | Slightly Destabilizing |
| K82N | -0.206 | Moderate Destabilizing |
| G84R | -0.345 | Moderate Destabilizing |
| L91I | 1.344 | Very High Stabilizing |
| G94V | 1.045 | High Stabilizing |
| V100I | 0.232 | Low Stabilizing |
| N106S | -0.828 | High Destabilizing |
| A114S | -1.055 | High Destabilizing |
| P119A | -0.312 | Moderate Destabilizing |
| L131F | -0.206 | Moderate Destabilizing |
| S132N | 0.643 | Moderate Stabilizing |
| A144V | 0.979 | High Stabilizing |
| K150N | -0.468 | Moderate Destabilizing |
| K151T | -0.216 | Moderate Destabilizing |
| S165I | 1.161 | High Stabilizing |
| M181V | -0.736 | Moderate Destabilizing |
| H184Y | 1.09 | High Stabilizing |
| F185L | -0.374 | Moderate Destabilizing |
| A189T | -0.023 | Slightly Destabilizing |
| Q190K | 0.271 | Moderate Stabilizing |
| C194W | -0.586 | Moderate Destabilizing |
| H196R | -0.505 | Moderate Destabilizing |
| P203L | 0.558 | Moderate Stabilizing |
| P205H | 0.893 | High Stabilizing |
| K222N | -1.472 | Very High Destabilizing |
| R234L | -0.317 | Moderate Destabilizing |
| K242N | -0.124 | Low Destabilizing |
| G244W | -0.287 | Moderate Destabilizing |
| K252N | 0.092 | Slightly Stabilizing |
| Y253S | -2.944 | Very High Destabilizing |
| Y254D | -1.886 | Very High Destabilizing |
| P261H | 0.158 | Low Stabilizing |
| P265T | 0.546 | Moderate Stabilizing |
| Q298H | 0.138 | Low Stabilizing |
| L304V | -1.409 | Very High Destabilizing |
| E309K | 0.367 | Moderate Stabilizing |
| D317H | 0.56 | Moderate Stabilizing |
| D322V | -0.168 | Low Destabilizing |
| V332M | -0.166 | Low Destabilizing |
| S345I | 0.009 | Slightly Stabilizing |
| R357Q | -0.126 | Low Destabilizing |
| R374H | 0.119 | Slightly Stabilizing |
| W377C | -1.314 | Very High Destabilizing |
| V379M | -0.04 | Slightly Destabilizing |
| Y387C | 0.136 | Low Stabilizing |
| P395H | 1.507 | Very High Stabilizing |
| D397N | 0.288 | Moderate Stabilizing |
| D412N | -0.193 | Moderate Destabilizing |
| S416F | 0.498 | Moderate Stabilizing |
| S428P | 0.055 | Slightly Stabilizing |
| P429S | 0.396 | Moderate Stabilizing |
| R438Q | -0.147 | Low Destabilizing |
| G451D | 0.156 | Low Stabilizing |
| S456F | 1.041 | High Stabilizing |
| R459C | -0.373 | Moderate Destabilizing |
| G485W | -0.166 | Low Destabilizing |
| D495H | 0.347 | Moderate Stabilizing |
| A496S | -0.114 | Slightly Destabilizing |
| G500N | -1.382 | Very High Destabilizing |
| V507I | 0.033 | Slightly Stabilizing |
| A517S | -0.065 | Slightly Destabilizing |
| H528R | -0.242 | Moderate Destabilizing |
| L540V | -0.416 | Moderate Destabilizing |
| Q548H | -0.04 | Slightly Destabilizing |
| G554D | 0.63 | Moderate Stabilizing |
| Y557C | -1.211 | High Destabilizing |
| D568N | -0.245 | Moderate Destabilizing |
| A574T | 0.999 | High Stabilizing |
| N575H | 1.141 | High Stabilizing |
| I579M | -0.021 | Slightly Destabilizing |
| D582H | 0.338 | Moderate Stabilizing |
| R593H | -0.699 | Moderate Destabilizing |
| R598C | -0.298 | Moderate Destabilizing |
| Q599R | -0.182 | Low Destabilizing |
| N601Y | 0.564 | Moderate Stabilizing |
| Q608P | 0.121 | Low Stabilizing |
| E612Q | 0.195 | Low Stabilizing |
| V631I | 0.284 | Moderate Stabilizing |
| G636W | -0.503 | Moderate Destabilizing |
| G637V | 1.269 | Very High Stabilizing |
| S642W | 0.65 | Moderate Stabilizing |
| T643M | 0.714 | Moderate Stabilizing |
| R648H | -0.026 | Slightly Destabilizing |
| G649E | -0.075 | Slightly Destabilizing |
| D655N | 0.354 | Moderate Stabilizing |
| A666S | 0.374 | Moderate Stabilizing |

**Table S7:** Phosphorylation Status and Scores for Wild Type and Mutated FERMT1 for Experimental Phosphorylation Sites. Y, phosphorylated; N, non-phosphorylated.

| **Mutation** | **T8** | **Score** | **T30** | **Score** |
| --- | --- | --- | --- | --- |
| **FERMT1 WT** | Y | 0.641 | Y | 0.228 |
| **A155D, S502Y** | N | — | Y | 0.225 |
| **A244S** | N | — | Y | 0.225 |
| **A345V, F436S, E624K** | N | — | Y | 0.225 |
| **A402V** | N | — | Y | 0.225 |
| **A533T** | N | — | Y | 0.225 |
| **D252Y** | N | — | Y | 0.225 |
| **D306Y** | N | — | Y | 0.225 |
| **E509K, R288Q** | N | — | Y | 0.225 |
| **E645K** | N | — | Y | 0.225 |
| **F146Y, S53F** | N | — | Y | 0.225 |
| **G249C** | N | — | Y | 0.225 |
| **G576R** | N | — | Y | 0.225 |
| **I124T** | N | — | Y | 0.225 |
| **I300V** | N | — | Y | 0.225 |
| **L31Q** | N | — | Y | 0.303 |
| **L61F, T91I** | N | — | Y | 0.223 |
| **L376R** | N | — | Y | 0.225 |
| **M477I, P210S, A60V** | N | — | Y | 0.225 |
| **M513T** | N | — | Y | 0.225 |
| **P92R** | N | — | Y | 0.225 |
| **P159L** | N | — | Y | 0.225 |
| **Q26K, R17S, H38N** | N | — | Y | 0.184 |
| **R17S** | N | — | Y | 0.224 |
| **R255C, I160T** | N | — | Y | 0.225 |
| **R255C, R526K, I160T** | N | — | Y | 0.225 |
| **R526K, I300V** | N | — | Y | 0.225 |
| **R554W, R297W** | N | — | Y | 0.225 |
| **S133L** | N | — | Y | 0.225 |
| **S323R, T673N** | N | — | Y | 0.225 |
| **S586L** | N | — | Y | 0.225 |
| **T30K, A316E** | N | — | N | — |
| **T212K, P210T** | N | — | Y | 0.225 |
| **W63C** | N | — | Y | 0.221 |
| **W63L** | N | — | Y | 0.226 |
| **Y235S** | N | — | Y | 0.225 |
| **Y244S** | N | — | Y | 0.225 |
| **Y320C** | N | — | Y | 0.225 |
| **Y320H** | N | — | Y | 0.225 |
| **D265EDEL** | N | — | Y | 0.227 |
| **F118LDEL** | N | — | Y | 0.229 |
| **F206LINS** | N | — | Y | 0.227 |
| **N497TDEL** | N | — | Y | 0.231 |
| **Q226PINS** | N | — | Y | 0.226 |
| **Q226SDEL** | N | — | Y | 0.228 |
| **S586HDEL** | N | — | Y | 0.23 |

**Table S8:** Phosphorylation Status and Scores for Wild Type and Mutated FERMT2 for Experimental Phosphorylation Sites. Y, phosphorylated; N, non-phosphorylated.

| **Structure** | **Y193** | **Score** | **S159** | **Score** | **S181** | **Score** | **S666** | **Score** |
| --- | --- | --- | --- | --- | --- | --- | --- | --- |
| **FERMT2** | Y | 0.317 | Y | 0.079 | Y | 0.509 | Y | 0.447 |
| **D57DEL** | N | — | N | — | N | — | N | — |
| **D459A** | N | — | Y | 0.059 | Y | 0.509 | N | — |
| **E583KDEL** | N | — | Y | 0.079 | Y | 0.407 | N | — |
| **G5DEL** | N | — | N | — | N | — | N | — |
| **I324DEL** | N | — | Y | 0.079 | Y | 0.44 | N | — |
| **K152DEL** | N | — | N | — | N | — | N | — |
| **K153INS** | N | — | N | — | N | — | N | — |
| **K154DEL** | N | — | N | — | N | — | N | — |
| **K386DEL** | Y | 0.317 | Y | 0.079 | Y | 0.509 | N | — |
| **L387INS** | N | — | N | — | N | — | N | — |
| **R290DEL** | Y | 0.317 | Y | 0.079 | Y | 0.509 | N | — |
| **Y530DEL** | Y | 0.317 | Y | 0.079 | Y | 0.509 | N | — |
| **A288S** | Y | 0.311 | Y | 0.079 | Y | 0.509 | Y | 0.463 |
| **A288V** | Y | 0.311 | Y | 0.079 | Y | 0.509 | Y | 0.463 |
| **A418T** | Y | 0.311 | Y | 0.079 | Y | 0.509 | Y | 0.463 |
| **A542C** | Y | 0.311 | Y | 0.079 | Y | 0.509 | Y | 0.463 |
| **C309S** | Y | 0.311 | Y | 0.079 | Y | 0.509 | Y | 0.463 |
| **D281Y** | Y | 0.311 | Y | 0.079 | Y | 0.509 | Y | 0.463 |
| **D281E** | Y | 0.311 | Y | 0.079 | Y | 0.509 | Y | 0.463 |
| **E427D** | Y | 0.311 | Y | 0.079 | Y | 0.509 | Y | 0.463 |
| **E665G** | Y | 0.311 | Y | 0.079 | Y | 0.509 | Y | 0.451 |
| **F439C** | Y | 0.311 | Y | 0.079 | Y | 0.509 | Y | 0.463 |
| **G176E** | Y | 0.311 | Y | 0.079 | Y | 0.51 | Y | 0.463 |
| **H23Y** | Y | 0.311 | Y | 0.079 | Y | 0.509 | Y | 0.463 |
| **H323D** | Y | 0.311 | Y | 0.079 | Y | 0.509 | Y | 0.463 |
| **I291S** | Y | 0.311 | Y | 0.079 | Y | 0.509 | Y | 0.463 |
| **I307T** | Y | 0.311 | Y | 0.079 | Y | 0.509 | Y | 0.463 |
| **P135L** | Y | 0.311 | Y | 0.079 | Y | 0.509 | Y | 0.463 |
| **R595W** | Y | 0.311 | Y | 0.079 | Y | 0.509 | Y | 0.463 |
| **R659C** | Y | 0.311 | Y | 0.079 | Y | 0.509 | Y | 0.463 |
| **S199R** | Y | 0.311 | Y | 0.079 | Y | 0.509 | Y | 0.461 |
| **S484F** | Y | 0.311 | Y | 0.079 | Y | 0.509 | Y | 0.463 |
| **T129I** | Y | 0.311 | Y | 0.079 | Y | 0.509 | Y | 0.463 |
| **T362K** | Y | 0.311 | Y | 0.079 | Y | 0.509 | Y | 0.463 |
| **T536I** | Y | 0.311 | Y | 0.079 | Y | 0.509 | Y | 0.463 |
| **V24A** | Y | 0.311 | Y | 0.079 | Y | 0.509 | Y | 0.463 |
| **V49G** | Y | 0.311 | Y | 0.079 | Y | 0.509 | Y | 0.463 |
| **W65G** | Y | 0.311 | Y | 0.079 | Y | 0.509 | Y | 0.463 |
| **Y277D** | Y | 0.311 | Y | 0.079 | Y | 0.509 | Y | 0.463 |

**Table S9:** Phosphorylation Status and Scores for Wild Type and Mutated FERMT3 for Experimental Phosphorylation Sites. Y, phosphorylated; N, non-phosphorylated.

| **Mutation** | **S484** | **Score** | **T482** | **Score** |
| --- | --- | --- | --- | --- |
| **FERMT3 WT** | N | — | Y | 0.454 |
| **F121V** | N | — | Y | 0.454 |
| **D258Y** | N | — | Y | 0.454 |
| **G500N** | N | — | Y | 0.454 |
| **G572V** | N | — | Y | 0.454 |
| **G637V** | N | — | Y | 0.454 |
| **I42V** | N | — | Y | 0.454 |
| **K222N** | N | — | Y | 0.454 |
| **L91I** | N | — | Y | 0.454 |
| **L304V** | N | — | Y | 0.454 |
| **P136L** | N | — | Y | 0.454 |
| **P395H** | N | — | Y | 0.454 |
| **R107S** | N | — | Y | 0.454 |
| **R276Q** | N | — | Y | 0.454 |
| **R580G** | N | — | Y | 0.454 |
| **S186L** | N | — | Y | 0.454 |
| **S232L** | N | — | Y | 0.454 |
| **S233L** | N | — | Y | 0.454 |
| **S305F** | N | — | Y | 0.454 |
| **V20G** | N | — | Y | 0.454 |
| **V45G** | N | — | Y | 0.454 |
| **W65R** | N | — | Y | 0.454 |
| **W377C** | N | — | Y | 0.454 |
| **Y253S** | N | — | Y | 0.454 |
| **Y254D** | N | — | Y | 0.454 |
| **Y557C** | N | — | Y | 0.454 |
| **A126V, S468I** | N | — | Y | 0.454 |
| **A275S, V619L** | N | — | Y | 0.454 |
| **D245Y, G636R, A63S** | N | — | Y | 0.454 |
| **E612K, A89T, V585M, R542H** | N | — | Y | 0.453 |
| **G637V, S345I** | N | — | Y | 0.454 |
| **G637V, L104F** | N | — | Y | 0.454 |
| **K222N, E24Q** | N | — | Y | 0.454 |
| **Q599R, G44E** | N | — | Y | 0.454 |
| **R113L, S328I** | N | — | Y | 0.454 |
| **R523W, A295V** | N | — | Y | 0.438 |
| **S165I, R510H** | N | — | Y | 0.454 |
| **V100I, L46F** | N | — | Y | 0.454 |
| **V558F, A539S** | N | — | Y | 0.455 |
| **W65L, G43C** | N | — | Y | 0.454 |
| **W65L, V36F** | N | — | Y | 0.454 |

**Table S10:** List of Kindlin-associated Mechanosensitive Proteins that are Components of Major Mechano-chemical Signaling. TF, transcription factor; MIC, mechano-chemical ion channels; MRP, mechano-chemical receptor proteins; MCP, mechano-sensitive cytoskeletal proteins; OTHERS, proteins with other cellular functions; indicated by different colors.

| Name of protein | Type | Major Cellular function | Kindlin involved | Refs. |
| --- | --- | --- | --- | --- |
| SRF | TF | EMT | Kindlin 3, Kindlin 2 | 20 |
| MYOCD | TF | EMT | Kindlin 2 | 21 |
| MRTF-A | TF | EMT | Kindlin 3 | 22 |
| MRTF-B | TF | EMT | Kindlin 2 | 22 |
| YY1AP1 | TF | Invasion | Kindlin 2 | 23 |
| TAFAZZIN | TF | Invasion | Kindlin 2 | 24 |
| CTNNB | TF | EMT | Kindlin 2 | 25 |
| NFKB | TF | EMT | Kindlin 2 | 26 |
| FOXP3 | TF | Invasion | Kindlin 2 | 27 |
| SMAD | TF | EMT (involved in TGF-beta pathway) | Kindlin 2 | 28 |
| AP1 | TF | EMT/ Cell proliferation/apoptosis | Kindlin 2 | 29 |
| KLF2 | TF | Quiescence/proliferation | Kindlin 2 | 30 |
| KLF4 | TF | Cell cycle arrest/apoptosis | Kindlin 2 | 31 |
| GATA4 | TF | Survival (anti-apoptotic) | Kindlin 2 | 32 |
| CEBP | TF | Proliferation/apoptosis/differentiation | Kindlin 2 | 33 |
| ROCK1 | TF | Cell migration/adhesion/proliferation | Kindlin 2 | 34 |
| MURF1 | TF | Protein breakdown – muscle atrophy | Kindlin 2 | 35 |
| P53 | TF | Tumor suppressor/controls proliferation | Kindlin 2 | 36 |
| P21 | TF | Cell cycle inhibitor/ anti-proliferation | Kindlin 2 | 36 |
| TCF4 | TF | Regulate gene expression in immune system and brain during development | Kindlin 2 | 37 |
| TRAAK | **MIC** | Nervous system – signalling | Kindlin 1 | 38 |
| PIEZO1 | **MIC** | Mechanotransduction processes | Kindlin 2 | 39 |
| PIEZO2 | **MIC** | Mechanotransduction processes | Kindlin 2 | 40 |
| TRPV | **MIC** | Signalling – homeostasis | Kindlin 1 | 41 |
| INTEGRIN | **MRP** | Signalling/adhesion | Kindlin 1,<br>Kindlin 2,<br>Kindlin 3 | 42–44 |
| EMR2/ADGRE2 | **MRP** | Adhesion/migration of myeloid cells | Kindlin 2 (Based on ITGB2) | 45 |
| CD97/ADGRE5 | **MRP** | Invasion | Kindlin 3 | 46 |
| GPR56/ADGRG1 | **MRP** | Inhibits over-migration of neurons | Kindlin 2 | 47 |
| TGF-BETA | **MRP** | EMT, immunosuppression | Kindlin 1,<br>Kindlin 2 | 48,49 |
| TMX4 | **MRP** | ER reductase – protein folding | Kindlin 1,<br>Kindlin 2 | 50 |
| Talin | **MCP** | Cell-ECM communication | Kindlin 1,<br>Kindlin 2,<br>Kindlin 3 | 42,51,52 |
| Actin | **MCP** | Cytoskeletal structure- focal adhesions (migration, contractility, shape) | Kindlin 2 | 53 |
| Vinculin | **MCP** | Cell adhesion | Kindlin 2 | 54 |
| Paxillin | **MCP** | Signalling | Kindlin 2 | 51,55 |
| CTNNA | **MCP** | Adherens junction | Kindlin 2 | 37 |
| Vimentin | **MCP** | EMT | Kindlin 2 | 56 |
| Alpha Tubulin | **MCP** | Cell division | Kindlin 2 | 57 |
| Beta Tubulin | **MCP** | Cell division | Kindlin 2 | 58 |
| Arp2 | **MCP** | Cytoskeleton- branched nucleation of F-actin | Kindlin 2 | 55 |
| Arp3 | **MCP** | Cytoskeleton- branched nucleation of F-actin | Kindlin 2 | 55 |
| ILK | **MCP** | Invasion | Kindlin 1, Kindlin 2, Kindlin 3 | 59,60 |
| GIV | **MCP** | Signalling | Kindlin 2 | 61 |
| PARVA | **MCP** | Cell survival/invasion | Kindlin 2 | 62 |
| LMNA | **MCP** | Cell proliferation | Kindlin2 (via F-actin) | 63 |
| FYB | **OTHERS** | Cell motility, proliferation, and activation | Kindlin 3 | 64 |
| APBB1IP | **OTHERS** | Immune response regulator and cancer cell invasion | Kindlin 3 | 65 |
| FBLIM | **OTHERS** | Cell-ECM adhesion | Kindlin 1, Kindlin 2 | 66 |
| GNB2L1 | **OTHERS** | Cell signalling | Kindlin 3 | 67 |
| SKI2W | **OTHERS** | DNA replication/repair and translation (helicase activity) | Kindlin 2 | Via the STRING Database |
| CLATHRIN | **OTHERS** | Cell interactions, signalling, homeostasis- endocytosis | Kindlin 2 | 68 |
| DNMT1 | **OTHERS** | DNA methylation | Kindlin 2 | 69 |
| DNMT3a | **OTHERS** | DNA methylation | Kindlin 2 | 70 |
| Bardet-Biedl syndrome proteins | **OTHERS** | Ciliary trafficking | Kindlin 2 | 71 |
| RAC1 | **OTHERS** | Cytoskeleton regulation | Kindlin 3 | 72 |
| RAC3 | **OTHERS** | Invasion | Kindlin 3 | 73 |
| GSK3BETA | **OTHERS** | Apoptosis/cell motility | Kindlin 2 | 37 |
| COL1A1 | **OTHERS** | Cytoskeleton regulation, cell motility | Kindlin2 | 74,75 |
| FN1 | **OTHERS** | Cell adhesion, migration | Kindlin2 | 76,77 |

